# GWAS and PheWAS of Red Blood Cell Components in a Northern Nevadan Cohort

**DOI:** 10.1101/475467

**Authors:** Karen A. Schlauch, Robert W. Read, Gai Elhanan, William J Metcalf, Anthony D. Slonim, Ramsey Aweti, Robert Borkowski, Joseph J. Grzymski

## Abstract

In this study, we perform a full genome-wide association study (GWAS) to identify statistically significantly associated single nucleotide polymorphisms (SNPs) with three red blood cell (RBC) components and follow it with two independent PheWASs to examine associations between phenotypic data (case-control status of diagnoses or disease), significant SNPs, and RBC component levels. We first identified associations between the three RBC components: mean platelet volume (MPV), mean corpuscular volume (MCV), and platelet counts (PC), and the genotypes of approximately 500,000 SNPs on the Illumina Infimum^®^ DNA Human OmniExpress-24 BeadChip using a single cohort of 4,700 Northern Nevadans. Twenty-one SNPs in five major genomic regions were found to be statistically significantly associated with MPV, two regions with MCV, and one region with PC, with *p*<5x10^-8^. Twenty-nine SNPs and nine chromosomal regions were identified in 30 previous GWASs, with effect sizes of similar magnitude and direction as found in our cohort. The two strongest associations were SNP rs1354034 with MPV (*p*=2.4x10^-13^) and rs855791 with MCV (*p*=5.2x10^-12^). We then examined possible associations between these significant SNPs and incidence of 1,488 phenotype groups mapped from International Classification of Disease version 9 and 10 (ICD9 and ICD10) codes collected in the extensive electronic health record (EHR) database associated with Healthy Nevada Project consented participants. Further leveraging data collected in the EHR, we performed an additional PheWAS to identify associations between continuous red blood cell (RBC) component measures and incidence of specific diagnoses. The first PheWAS illuminated whether SNPs associated with RBC components in our cohort were linked with other hematologic phenotypic diagnoses or diagnoses of other nature. Although no SNPs from our GWAS were identified as strongly associated to other phenotypic components, a number of associations were identified with *p*-values ranging between 1x10^-3^ and 1x10^-4^ with traits such as respiratory failure, sleep disorders, hypoglycemia, hyperglyceridemia, GERD and IBS. The second PheWAS examined possible phenotypic predictors of abnormal RBC component measures: a number of hematologic phenotypes such as thrombocytopenia, anemias, hemoglobinopathies and pancytopenia were found to be strongly associated to RBC component measures; additional phenotypes such as (morbid) obesity, malaise and fatigue, alcoholism, and cirrhosis were also identified to be possible predictors of RBC component measures.

**Author Summary:** The combination of electronic health records and genomic data have the capability to revolutionize personalized medicine. Each separately contains invaluable data; however, combined, the two are able to identify new discoveries that may have long-term health benefits. The Healthy Nevada Project is a non-profit initiative between Renown Medical Center and the Desert Research Institute in Reno, NV. The project has so far collected a cohort of 6,500 Northern Nevadans, with extensive medical electronic health records in the Renown Health database. Combining the genotypes of these participants with the clinical data, this study’s aim is to find associations between genotypes (genes) and phenotypes (diagnoses and lab records). Here, we identify and examine clinical associations with red blood cell components such as platelet counts and mean platelet volume. These are components that have clinical relevance for several diseases, such as anemia, atherothrombosis and cancer. Our results from genome wide association studies mirror previous studies, and identify new associations. The extensive electronic health records enabled us to perform phenome wide associations to discover strong associations with hematologic components, as well as other important traits and diagnoses.

## Introduction

The complete blood count (CBC) is a widely used medical diagnostic test that is a compilation of the number, size, and composition of various components of the hematopoietic system. Abnormal CBC measures may indicate illness or disease. Mean corpuscular volume (MCV), platelet count (PC), and mean platelet volume (MPV) are specific CBC characteristics (hereby called RBC components), and linked to complex disorders such as anemia, alpha thalassemia and cardiovascular disease [1–5]. Platelets are involved in vascular integrity, wound healing, immune and inflammatory responses, and tumor metastasis; the role of platelets is also paramount in hemostasis and in the pathophysiology of atherothrombosis and cancer [6–12]. Additionally, abnormally high mean platelet volumes (MPV) are considered a predictor of post event outcome in coronary disease and myocardial infarction [13]. Furthermore, studies have shown that individuals living in higher altitudes have noted differences in red blood cell components than at sea level. At approximately 4,400 feet above sea level, Northern Nevada, where this study is conducted, is considered a high desert in the Sierra Nevada foothills. Alper showed that mean platelet volume (MPV) is 7.5% higher at altitudes greater than 4,000 feet than at sea level [14]. Similarly, Hudson showed a notable and statistically significant positive correlation with platelet counts (PC) and altitude [15], while mean corpuscular volume (MCV) was recorded as lower at higher altitudes than at sea level [16]. As RBCs help transport oxygen throughout the entire body, the identification of RBC-related genotypic mutations, especially in an RBC high-turnover environment is valuable. Lastly, the identification of genomic regions with roles in megakaryopoiesis and platelet formation, as well as neoplastic conditions like polycythemia vera and essential thrombocytosis (ET) [17,18], may help identify those that have a higher risk of certain complex RBC diseases.

Given the importance of these three RBC components, we conducted a study to identify both genetic and phenotypic associations with all three characteristics via GWASs and PheWASs. Our study begins with the Healthy Nevada Project, a single cohort formed in 2016 to investigate factors that may contribute to health outcomes in Northern Nevada. Its first phase provided 10,000 individuals in Northern Nevada with genotyping on the 23andMe 2016 Illumina Human OmniExpress-24 BeadChip platform at no cost. Renown Hospital is the largest hospital in the area, and 75% of these 10,000 individuals are cross-referenced in its extensive EHR database As noted above, previous GWASs have identified significant genetic links with all three RBC components we examine in this study, MPV, MCV and PC [13,17–45]. Lin et al. 2007 identified a strong genetic link with MCV in region 11p15 using the Framingham cohort [19]; Kullo et al. 2010 leveraged EHR data from the Mayo Clinic to detect four genes strongly associated with at least one of the three RBC components [27]. Similarly, a number of regions were linked with PC in an African American cohort [35] and MPV [35]; Shameer detected five regions associated with PC and eight with MPV [18].

Our study first performed a genome-wide association study (GWAS) of 4,700 genotyped Northern Nevadans who have at least one recorded value for one of the three RBC components MPV, MCV and PC to examine the genetic component of these components. We found 38 SNPs to be statistically significantly associated (*p*<5x10^-8^) to one of the three RBC components. Many of these associations were previously reported, yet our study did identify nine novel SNPs in six different regions. While there were few new associations discovered in our cohort, we identified several SNPs that fall within genes influencing megakaryocytes maturation, platelet volume, platelet signaling and diseases such as anemia. Further, with extensive linked electronic medical record (EMR) data, we had the ability to perform a PheWAS of 1,488 standard lab results (phenotypes) against each SNP found to be associated to RBC components in the Northern Nevadan cohort to examine pleiotropy. Additionally, we then examined the RBC components phenotypically, using linked electronic medical record (EMR) data to determine the relationship between measures of each component and a variety of clinical conditions recorded in patients. Many relevant and strongly statistically significant associations were identified, especially with hematologic components; other traits not currently shown to be linked to RBC components, such as obesity, alcoholism and cirrhosis, were also detected.

## Results

### Characteristics of cohort

We examined 4,700 genotyped individuals with at least one recorded RBC measure; 4,590 individuals in the cohort had measures for all three components. Table 1 describes the cohort with respect to gender, age, ethnic origin, and standardized value of each RBC component. Note that all values for each component were standardized to the most current lab test administered for that component via linear transformation. The normal (healthy) reference values to which all individual records were standardized are also presented in Table 1. The mean standardized RBC component values for each individual are available in Supplementary Table S1.

**Table 1.** Cohort Characteristics

|  |  |  |
| --- | --- | --- |
| Age (years) | 47.24 ± 15.82 |  |
| Male (%) | 1328 (28.24) |  |
| African American (%) | 53 (1.12) |  |
| Asian (%) | 100 (2.12) |  |
| Caucasian (%) | 4,202 (89.35) |  |
| Latino (%) | 138 (2.93) |  |
| Native American (%) | 30 (0.64) |  |
| Pacific Islander (%) | 11 (0.23) |  |
| Unknown (%) | 168 (3.6) |  |
|  | **Standardized Component levels** | **Normal Reference Ranges** |
| MPV (fL) | 10.58 ± 0.98 | [9, 12.9] (fL) |
| MCV (fL) | 91.53 ± 4.47 | [81.4, 97.8] (fL) |
| PC (K/uL) | 251.57 ± 62.23 | [164, 446] (K/uL) |

### GWAS of RBC components

Using the average measures of each individual’s MPV, PC and MCV lab records, a standard GWAS under the additive model with adjustments for gender, age and the first four principal components was performed using *PLINK* 1.9. Genomic inflation coefficients (lambda) were computed for each cohort: 1.031 for MPV, 1.027 for PC, and 1.045 for MCV.

Any SNP with an association *p*-value of *p*<5x10^-8^ was considered a statistically significant association, following current standards [28,32,46,47]. The percentage of phenotypic variance attributed to genetic variation was computed with a combination of *PLINK* and GCTA [48]: genetic variance was 35.3% for MCV; 32.2% for MPV; 20.7% for PC. The three individual GWAS studies identified a total of 38 SNPs that associated with a RBC component with statistical significance. Manhattan plots of the three GWAS results are presented in Supplemental Figure S1(A-C). As an example for the reader, we include in the manuscript (Fig. 1), a Manhattan plot for MCV.

**Fig 1:**
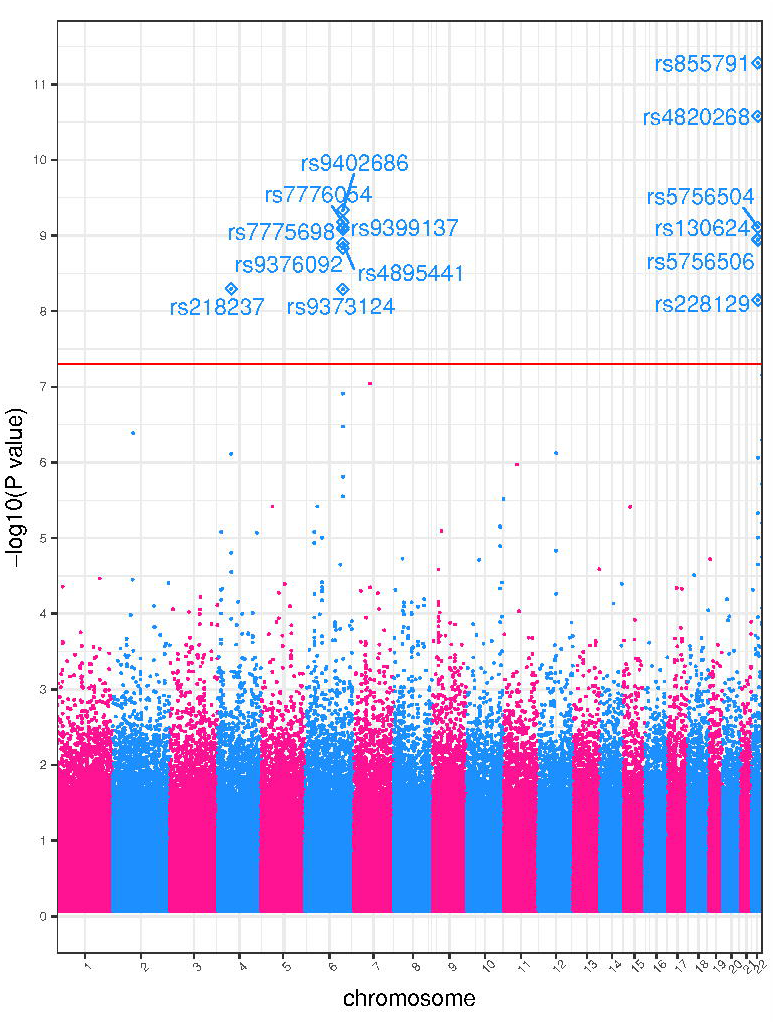
MCV GWAS Manhattan plot. Genome-wide association study results for MCV. The x-axis represents the genomic position of 498,916 SNPs. The y-axis represents -log_10_-transformed raw *p*-values of each genotypic association. The red horizontal line indicates the threshold of significance *p*=5x10^-8^.

### MPV

A GWAS was performed on a cohort of 4,591 genotyped participants with MPV laboratory measures. We identified 21 SNPs across five different chromosomal regions that reached genome-wide significance (*p* < 5x10^-8^; Table 2). Of these, 13 demonstrated previous associations in at least one other study, with six associated with RBC components (Supplementary Table S2) [13,17,18,25,28,30,33,35,49-58]. All five significant chromosomal regions were previously associated with MPV[17,18]. The fifth region 18q22.2, contains three SNPs associated in our cohort with average *p*-value *p*=3.86x10^-9^, however none of the individual SNPs have been previously associated with MPV. Results are presented in Table 2.

**Table 2.** Statistically Significant GWAS SNPs

| rsID | Chrom | Cyto Region | Associated Gene | Minor Allele | MAF | $\beta$ | (SE) | GWAS $p$-value | Mutation Classification | RBC |
| --- | --- | --- | --- | --- | --- | --- | --- | --- | --- | --- |
| rs10274553 | chr3 | p14.3 | ARHGEF3 | C | 49.74 | -0.1198 | 0.020 | $3.82 \times 10^{-9}$ | intron | MPV |
| rs10509186 | chr3 | p14.3 | ARHGEF3 | T | 45.55 | -0.1187 | 0.021 | $7.75 \times 10^{-9}$ | intron | MPV |
| rs10822186 | chr7 | q22.3 | NA | G | 49.22 | -0.1107 | 0.020 | $4.38 \times 10^{-8}$ | unknown | MPV |
| rs11130549 | chr7 | q22.3 | NA | C | 34.11 | -0.1238 | 0.022 | $9.94 \times 10^{-9}$ | unknown | MPV |
| rs12355784 | chr7 | q22.3 | NA | A | 45.46 | -0.1184 | 0.021 | $9.32 \times 10^{-9}$ | unknown | MPV |
| rs1354034 | chr7 | q22.3 | NA | T | 41.14 | 0.1546 | 0.021 | 2.39x10<sup>-13</sup> | unknown | MPV |
| rs1788103 | chr7 | q22.3 | NA | G | 48.18 | -0.1261 | 0.020 | 5.15x10<sup>-10</sup> | unknown | MPV |
| rs1790588 | chr7 | q22.3 | NA | C | 48.01 | -0.1273 | 0.020 | 3.31x10<sup>-10</sup> | unknown | MPV |
| rs1790974 | chr10 | q21.3 | JMJD1C | T | 43.82 | -0.1128 | 0.020 | 3.32x10<sup>-8</sup> | intron | MPV |
| rs1935 | chr10 | q21.3 | JMJD1C | G | 45.58 | -0.1135 | 0.021 | 3.57x10<sup>-8</sup> | intron | MPV |
| rs201979226 | chr10 | q21.3 | JMJD1C | C | 48.78 | 0.1183 | 0.020 | 5.89x10<sup>-9</sup> | intron, near-gene-5 | MPV |
| rs342240 | chr10 | q21.3 | JMJD1C | A | 41.36 | 0.129 | 0.021 | 3.49x10<sup>-10</sup> | intron, untranslated-3 | MPV |
| rs342275 | chr10 | q21.3 | JMJD1C | T | 41.9 | 0.1292 | 0.020 | 2.96x10<sup>-10</sup> | intron | MPV |
| rs342293 | chr10 | q21.3 | JMJD1C | G | 44.31 | 0.1325 | 0.020 | 6.61x10<sup>-11</sup> | missense | MPV |
| rs342296 | chr10 | q21.3 | REEP3 | A | 44.03 | 0.131 | 0.020 | 1.04x10<sup>-10</sup> | intron | MPV |
| rs34818942 | chr12 | q24.31 | WDR66 | T | 7.29 | 0.254 | 0.039 | 7.77x10<sup>-11</sup> | intron | MPV |
| rs386614085 | chr12 | q24.31 | RHOF | G | 45.45 | -0.1172 | 0.021 | 1.21x10<sup>-8</sup> | intron | MPV |
| rs4379723 | chr18 | q22.2 | CD226 | C | 45.45 | -0.1172 | 0.021 | 1.29x10<sup>-8</sup> | missense | MPV |
| rs763361 | chr18 | q22.2 | CD226 | T | 47.24 | -0.1273 | 0.020 | 3.26x10<sup>-10</sup> | intron | MPV |
| rs7910927 | chr18 | q22.2 | CD226 | G | 45.52 | -0.1146 | 0.021 | 2.68x10<sup>-8</sup> | intron | MPV |
| rs7961894 | chr18 | q22.2 | DOK6 | T | 10.24 | 0.2221 | 0.033 | 2.68x10<sup>-11</sup> | untranslated-3 | MPV |
| rs218237 | chr4 | q12 | NA | T | 15.19 | 0.740 | 0.126 | 5.07x10<sup>-9</sup> | unknown | MCV |
| rs9402686 | chr6 | q23.3 | NA | A | 24.89 | 0.647 | 0.104 | 4.60x10<sup>-10</sup> | unknown | MCV |
| rs7776054 | chr6 | q23.3 | NA | G | 24.28 | 0.645 | 0.104 | 6.65x10<sup>-10</sup> | unknown | MCV |
| rs9399137 | chr6 | q23.3 | NA | C | 23.83 | 0.648 | 0.105 | 7.82x10<sup>-10</sup> | unknown | MCV |
| rs7775698 | chr6 | q23.3 | NA | T | 24.23 | 0.642 | 0.104 | 8.24x10<sup>-10</sup> | unknown | MCV |
| rs4895441 | chr6 | q23.3 | NA | G | 25.08 | 0.628 | 0.103 | 1.27x10<sup>-9</sup> | unknown | MCV |
| rs111194878 | chr6 | q23.3 | NA | A | 25.41 | 0.622 | 0.103 | 1.47x10<sup>-9</sup> | unknown | MCV |
| rs9373124 | chr6 | q23.3 | NA | C | 26.16 | 0.599 | 0.102 | 5.14x10<sup>-9</sup> | unknown | MCV |
| rs855791 | chr22 | q12.3 | TMPRSS6 | A | 44.2 | -0.621 | 0.090 | 5.23x10<sup>-12</sup> | missense | MCV |
| rs4820268 | chr22 | q12.3 | TMPRSS6 | G | 46.56 | -0.604 | 0.090 | 2.65x10<sup>-11</sup> | coding-synon | MCV |
| rs5756504 | chr22 | q12.3 | TMPRSS6 | T | 36.93 | 0.567 | 0.092 | 7.77x10<sup>-10</sup> | intron | MCV |
| rs130624 | chr22 | q12.3 | NA | G | 42.77 | 0.549 | 0.090 | 1.13x10<sup>-9</sup> | unknown | MCV |
| rs5756506 | chr22 | q12.3 | TMPRSS6 | C | 36.92 | 0.563 | 0.092 | 1.15x10<sup>-9</sup> | intron | MCV |
| rs386563505 | chr22 | q12.3 | NA | A | 40.75 | 0.525 | 0.091 | 7.12x10<sup>-9</sup> | unknown | MCV |
| rs385893 | chr9 | p24.1 | NA | T | 49 | -7.744 | 1.258 | 8.04x10<sup>-10</sup> | unknown | PC |
| rs10974808 | chr9 | p24.1 | RCL1 | G | 11.48 | 11.490 | 1.943 | 3.53x10<sup>-9</sup> | intron | PC |
| rs423955 | chr9 | p24.1 | NA | C | 34.17 | -7.387 | 1.325 | 2.64x10<sup>-8</sup> | near-gene-5 | PC |

### MCV

A GWAS was performed on a cohort of 4,699 genotyped participants with MCV laboratory measures. There were 14 SNPS that were significantly associated with MCV (Table 2). These SNPs lie in three chromosomal regions: predominantly in 6q23.3 and 22q12.3. These two regions have detailed annotation and were linked previously with MCV (Supplementary Table S2) [20,27,32]. All but four of the SNPs are in non-coding regions. These four SNPs lie in *TMPRSS6*. The gene *TMPRSS6* codes for the protein matriptase-2, which is part of a signaling pathway that regulates blood iron levels [31]. The two SNPS rs855791 and rs4820268 showed the strongest association with MCV (*p*<1x10^-11^). These two SNPS also lie in *TMPRSS6* and cause a missense and synonymous mutation, respectively. Results are presented in Table 2.

### PC

A GWAS was performed on a cohort of 4,700 genotyped participants with PC laboratory measures. Three SNPs were identified with statistically significant (*p*<5x10^-8^) links to PC values in our cohort, two of which were previously identified in other studies (Supplementary Table S2) [17,25,26,34]. The SNP rs10974808 is in the same cytoband region (9p24.1) as the others but has not been linked to PC. The three SNPs have different effects on PC: rs385893 and rs423955 have negative effect size (β=-7.744 and -7.387, respectively), while rs10974808 has a positive effect (β=11.490). The minor allele frequency of rs10974808 is much rarer (MAF=11.48%) compared to 49% for rs385893 and 31.17% for rs423955. Results are presented in Table 2.

### Comparison to other GWAS studies

The Northern Nevada cohort had mean standardized MPV values of 10.58 ± 0.98 fL, comparable to levels reported in the Health ABC cohort described in Qayyum (10.9 ± 1.6 fL), and two European cohorts investigated in Geiger (10.53 ± 1.08, 10.83 ± 0.87) [28,35]. The Nevadan cohort had MCV values of 91.53 ± 4.5 fL, also comparable to those described in Kullo (90.5 ± 4.2 fL) and Ding in the Mayo and Johns Hopkins Group Health Cooperative cohorts (90.53 ± 4.17 and 91.56 ± 4.49, respectively), as well as several European cohorts in Geiger (e.g., 91.5 ± 4.2, 91.4 ± 4.41, 91.1 ± 4.44, 92.0 ± 4.3) [27,28,32]. Mean standardized PC values in the Nevadan cohort (251 ± 62.23 K/uL) were very similar to many of the cohorts examined in Geiger (e.g., 258.6 ± 63.1, 252 ±71.7, 250.9 ±64.8, 247 ± 64.7) [28].

Our three GWAS results were in close correlation with many of the other studies. For example, the locus rs7961894 in the WDR66 gene on q24.31 was found associated to MPV in our cohort and in Meisinger as a top hit [24]. Effect sizes in Meisinger were larger than ours (1.03 vs. 0.22), but the number of minor alleles predicted an increase in MPV for both studies. Another SNP, rs342240, was one of our cohort’s top associations with MPV, and was also identified by Shameer and Soranzo as significant links to MPV [17,18]. Similarly, locus rs385893 was identified as a possible predictor of PC by Soranzo and our cohort, with very similar notably large negative effect sizes (-6.24 and -7.74, respectively). Kullo also found SNP rs7775698 to be significantly associated to MCV, with similar positive effect sizes as our study (0.92 vs 0.56) [27]. Soranzo et al. identified rs9402686 as a top link with MCV, and again, effect sizes were similar to ours (0.82 vs 0.65) [17].

### ANOVA

The mean component values across genotypes presented in Supplementary Table S2 correlate with negative and positive effect sizes: SNPs showing a negative effect size have a decrease in component values across the genotypes from left to right (homozygous in major allele, heterozygous, homozygous in minor allele). All ANOVA *p*-values of the significant SNPs identified in this study are significant, even after a simple Bonferonni correction (.05/38=0.001). A box and whisker figure of ANOVA results for the top hit SNP rs7961894 are shown in Supplementary Fig S2.

### PheWAS of RBC components

The first PheWAS examined possible associations between significant SNPs identified in each RBC trait GWAS and 1,488 phenotypic groups. At significance levels 1x10^-4^<*p*<1x10^-3^, putative associations of MCV-specific SNPs included respiratory failure; those with PC included GERD and other diseases of the esophagus. Our study also showed links with MPV-associated SNPs and skin cancer, hypoglycemia, hyperglyceridemia, IBS, among others. These associations are outlined in Supplementary Figs S3(A-C).

The second PheWAS investigated whether the 1,488 phenotype groups were associated with the levels of each RBC component; more specifically, the analysis identified whether the number of cases in a phenotype group was a predictor of the level of the component. For example, the PheWAS examining associations of MPV levels presented significant links with thrombocytopenia and purpura (*p*<1x10^-8^). Interestingly, Vitamin D deficiency was also shown to be a predictor of MPV levels, although at a lower significance level (*p*<1x10^-6^). Incidence of malaise and fatigue was also found to be a potential predictor of MPV in our cohort.

Associations with MCV included hemoglobinopathies and hemolytic anemias (*p*<1x10^-35^), as well as iron deficient anemias (*p*<1x10^-20^). Again, association with (morbid) obesity was evident (*p*<1x10^-20^). Alcoholism and related liver diseases were associated with MCV at a significance level of *p*<1x10^-8^; abnormal glucose and diabetes were also linked to MCV at *p*<1x10^-5^. We identified a strong association in our cohort between platelet counts and thrombocytopenia and purpura (*p*<1x10^-30^). Associations with other hematologic phenotypes such as various anemias and pancytopenia also reached significance (*p*<1x10^-8^). Additionally, (morbid) obesity and cirrhosis were statistically significantly associated with PC with *p*<1x10^-8^ significance level. These three PheWAS results are shown in Supplementary Fig S4(A-C). As an example for the reader, we include the PheWAS results for MCV in Fig 2.

**Fig 2:**
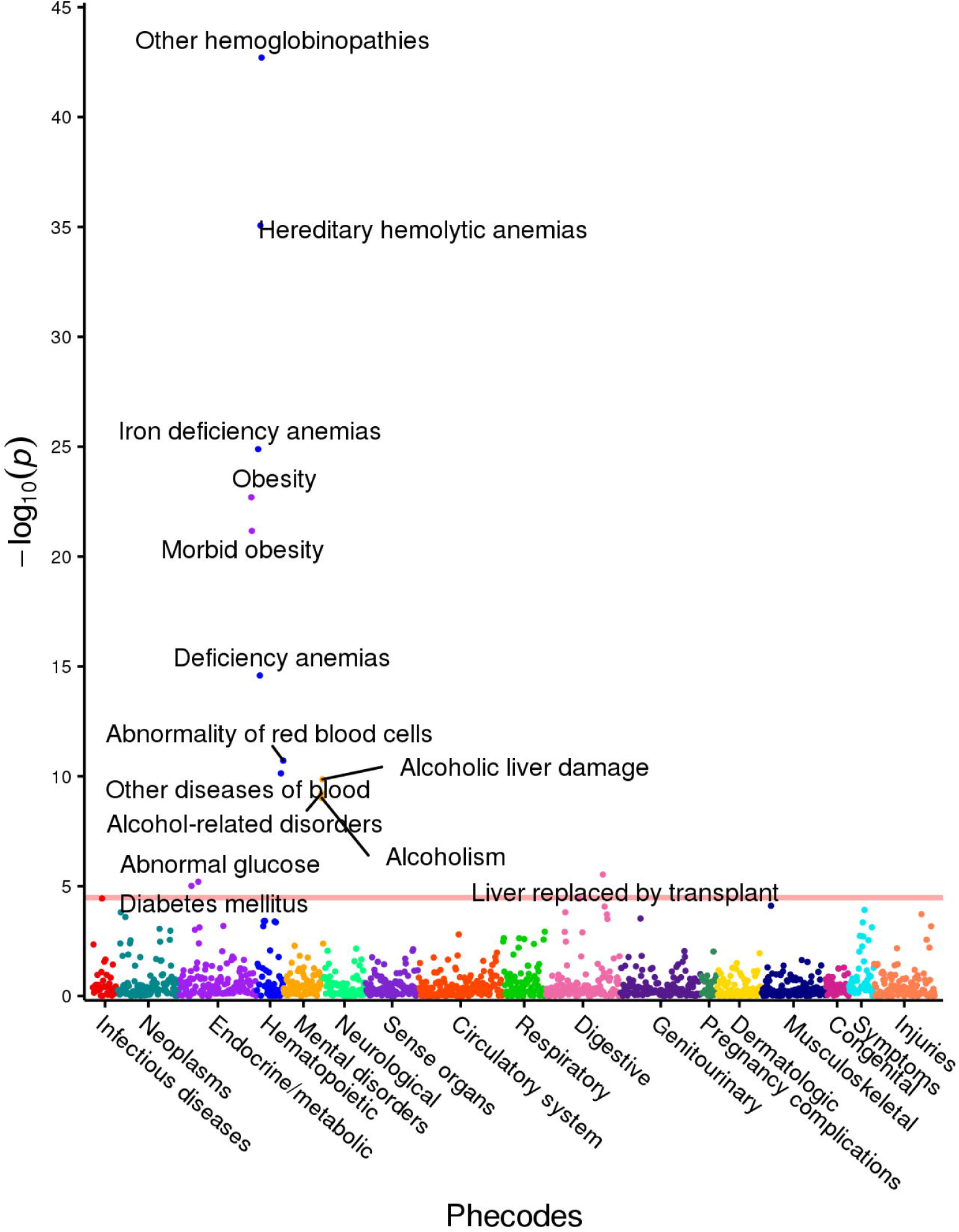
MCV PheWAS plot. This figure illustrates the results of individual linear regression between incidence of phenotype groups (phecodes) and continuous MCV component measures. The model includes age, gender and ethnicity as covariates. Each point represents the *p*-value of the association between one of 1,488 phecodes with at least 20 cases assigned to it, and the MCV component measure. The horizontal red line represents the significance level *p*=3.4x10^-5^.

## Discussion

In this study, we first performed three independent GWASs of 4,700 Healthy Nevada Project participants with 500,000 genotypes against the RBC components: platelet count, mean platelet volume and mean corpuscular volume. We followed these with two independent PheWASs for each component to identify additional phenotypic associations with each blood component-significant SNP, and phenotypic associations with measures of each blood component.

Our genome-wide association analysis identified ten different chromosomal cytoband regions associated with at least one RBC component. Nine of those regions were previously associated to RBC components in other studies; the region 22q13.33 represents a novel region in our study [17,18,20,25,27,28,30,32,49,59,60]. Nine genes lie in the cytoband regions: their functions are outlined in Table 3.

**Table 3.** Table Presenting Gene Functions

| Gene | Gene Description | Region | RBC | Function | Reference |
| --- | --- | --- | --- | --- | --- |
| *ARHGEF3* | Rho Guanine Nucleotide Exchange Factor 3 | p14.3 | MPV | Increases activity of Rho GTPases by catalyzing the release of bound GDP; may have a role in megakaryocytes maturation | [61] |
| *JMJD1C* | Histone Demethylase | q21.3 | MPV | Possible hormone-dependent transcriptional activation | [17] |
| *REEP3* | Receptor Accessory Protein 3 | q21.3 | MPV | Membrane protein | [18] |
| *WDR66* | WD Repeat Domain 66 | q24.31 | MPV | May create and alter platelet volumes | [24] |
| *RHOF* | Ras Homolog Family Member F | q24.31 | MPV | May regulate platelet filopodia formation | [62] |
| *CD226* | Cluster of Differentiation 226 | q22.22 | MPV | Catalyzes binding of activated platelets to endothelial cells; may have a role in in platelet signal transduction | [63] |
| *DOK6* | Docking Protein 6 | q22.2 | MPV | Protein scaffolding | [64] |
| *TMPRSS6* | Transmembrane Protease,<br>Serine 6 | q12.13 | MCV | Acts by cleaving hemojuvelin | [27,31,38] |
| *RCL1* | RNA Terminal Phosphate<br>Cyclase Like 1 | p24.1 | PC | rRNA processing | [28] |

Our GWAS results were very similar to previous MPV GWAS associations. The most significant genetic association with MPV (rs1354034, *p* = 2.39x10^-13^) is found in an intronic region within *ARHGEF3* on chromosome 3p14.3. The gene *ARHGEF3* codes for a Rho guanine nucleotide exchange factor 3 protein and was associated to MPV in previous studies [13,17,18,28,33,61], further demonstrating that our study was able to replicate associations with RBC components in prior single-cohort studies. The mechanism by which rs1354034 affects MPV values is still ambiguous. As it lies in a DNase I hypersensitive region within open chromatin, it could directly affect *ARHGEF3* expression in human megakaryocytes maturation [61]. Our second most significant association (rs7961894, *p*= 2.68x10^-11^) was also previously linked with MPV [13,18,24,28]. This SNP lies in intron 3 of *WDR66* on chromosome 12q24.31. Expression levels of *WDR66* have been directly tied to MPV, possibly indicating that *WDR66* is involved in the establishment of platelet volumes. SNP rs7961894 is not directly correlated with *WDR66* expression levels, implying an indirect role possibly through other regulatory mechanisms [24].

We also identified several SNPs on chromosome 10q21.3 to be associated with MPV in our cohort that were linked to sex hormone levels in previous studies [53]. This may imply a possible relationship between sex hormone levels and MPV. These SNPs almost exclusively lie in *JMJD1C*, a gene that encodes as a probable histone demethylase, and may have a function in hormone-dependent transcriptional activation [17]. This could indicate that the transcription of certain hematopoietic target genes may be enhanced or repressed when specific sex hormones are present; however, the exact targets and mechanisms have yet to be studied and clinical evidence for such association is scant.

Further, the chromosomal region 18q22.2 was shown to be associated with MPV [13], although the significant SNPs in this region have not been linked to MPV in previous studies. Three out of the four SNPs in this region are intronic to *CD226*, while one is in an untranslated region of *DOK6*. *CD226* codes for a protein, which mediates the binding of activated platelets to endothelial cells and may participate in platelet signal transduction [63]. Soranzo et al. also identified this gene as having a possible role in megakaryocyte (MK) development, thus these SNPs in *CD226* may influence platelet development [17]. *DOK6* encodes a docking protein, necessary for protein scaffolding, but to our knowledge has no known relation to platelet function; therefore, the functional relevance of a SNP in this gene is ambiguous. The mechanism by which these SNPS within 18q22.2 affect *CD226, DOK6* or MPV is also currently unknown.

The majority of SNPs associated with MCV and PC are in non-coding regions, and most were previously associated with these components in previous studies [17,27,32,45]. Our two strongest associations with MCV (rs855791, *p* = 5.23x10^-12^) and (rs4820268, *p* = 2.65x10^-11^) are in the gene *TMPRSS6* and could cause an altered or loss of function for the matriptase-2 protein. Altered function of the protein will likely influence iron status within the body, demonstrating why these SNPS are highly associated with anemia caused by iron deficiency [31,38]. PC was associated with only a single gene in our GWAS. This gene, *RCL1,* which encodes an RNA terminal phosphate cyclase-like 1 protein, was previously associated with PC [28]. The SNP associated to PC in this gene (rs10974808, *p*= 3.53x10^-09^) in our cohort has not been linked to PC by other studies to the best of our knowledge. Our strongest association (rs385893, *p* = 8.04×10^−10^) was previously found to affect *JAK2*, a gene 400 kb downstream of the locus and a key regulator of megakaryocyte maturation, illustrating that these SNPs may influence changes over large genetic regions [17]. This also highlights the difficulty determining the exact mechanisms by which these SNPS alter components, such as RBC, given their large theoretical range of influence.

We present here two comprehensive PheWAS analyses of RBC components. The first examines whether additional phenotypic associations exist between SNPs associated to an RBC component in our cohort. The second groups extensive EHR phenotypic data from the Healthy Nevada Project clinical database into 1,488 different phenotype groups and examines the association (predictive value) between their incidence rate with continuous RBC component values. To the best of our knowledge, this is the first PheWAS targeted at RBC components. Not surprisingly, many of our strongest associations were with hematopoietic phenotypes, indicating that the incidence of having one (or more) abnormal hematopoietic characteristics is a potential predictor of RBC component levels. Interestingly, the incidence of having vitamin D deficiency may be linked to MPV levels and requires further study, as incident solar radiation in the Northern Nevadan location of the study is high. Also of interest is that MCV and PC levels could be associated to the occurrence of (morbid) obesity, alcoholism and cirrhosis which are linked to poor vitamin D synthesis [65].

The identified associations between the RBD indices and hematopoietic findings and pathologies are mostly expected due to their known physiologic association and reconfirm previously reported findings. Iron deficiency anemia is often microcytic and characterized by reduced MCV [66]. Iron deficiency also affects megakaryocytes and may induce changes in megakaryocyte differentiation as well as increased platelet counts and volume [67]. As noted earlier, one of the strongest associations reported here is in the vicinity of JAK2, a known regulator of megakaryocytes maturation [68].

While thrombocytopenias are clearly synonymous with reduced PC, associated platelet volume and size changes can be used to differentiate between inherited macrothrombocytopenias and idiopathic thrombocytopenic purpura (ITP) [69], thus establishing an association with MPV that may be positive or inverse. While this study demonstrated a strong negative association between PC and purpura, and a positive association with MPV, it is important to note that not all purpuras are necessarily caused by platelet deficiency. However, phenotypic groupings were not specific enough to identify associations with respect to specific etiologies (See Supplementary Table S3).

Vitamin D, independently, and in association with platelet activity and increased platelet indices, has been associated with cardiovascular disease [70]. The positive association between vitamin D deficiency and MPV levels is intriguing and follows other findings. Cumhur et al. [71] observed an inverse correlation between vitamin D levels and MPV and hypothesized that this may be due to increased release of proinflammatory cytokines present with vitamin D deficiency. Park et al. also reported an inverse association between PC and MPV and vitamin D levels in adults [72].

Platelet activation, as evidenced by platelet indices, is a recognized phenomenon in metabolic syndrome [73,74]. This study resulted in a positive association between PC and morbid obesity, and a negative association between MCV and obesity and morbid obesity. While previous evidence [75] does not necessarily support all-gender association between obesity and increased platelet counts, our finding may reflect an association between the central obesity of metabolic syndrome and the associated platelet activation of metabolic syndrome. However, the phenotype groups were not specific enough to allow for specific differentiation between obesity types (See Supplementary Table S3).

Thrombocytopenia is often observed in chronic liver disease and cirrhosis and platelet activation may play a role in liver regeneration [76,77]. Alcoholism is also associated with thrombocytopenia [78]. However, evidence of an association between liver disease or alcoholism and platelet activation indices is lacking. Moreover, evidence points to platelet function defects in chronic alcoholism [79]. Thus, the negative effect of PC on cirrhosis and positive effect of MCV on cirrhosis, alcoholism, and alcohol-related disorders found in this study is intriguing and merits further confirmation and research.

## Materials and Methods

### The Renown EHR Database

The Renown Health EHR system was instated in 2007 on the EPIC system (EPIC System Corporation, Verona, Wisconsin, USA), and currently contains lab results, diagnosis codes (ICD9 and ICD10) and demographics of more than 1 million patients seen in the hospital system since 2005.

### Sample Collection

Saliva as a source of DNA was collected from 10,000 adults in Northern Nevada as the first phase of the Healthy Nevada Project to contribute to comprehensive population health studies in Nevada. The personal genetics company 23andMe^®^ was used to genotype these individuals. using the Orogene^®^ DX OGD-500.001 saliva kit [DNA Genotek, Ontario, Canada]. Genotypes are based on the Illumina Human OmniExpress-24 BeadChip platform [San Diego, CA, USA] including approximately 570,000 SNPs.

### IRB and Ethics Statement

The study was approved by our local Institutional Review Board (IRB, project 956068-12). Participants in the Healthy Nevada Project consent to having genetic information associated with electronic health information in a de-identified manner. Neither researchers nor participants have access to the complete EHR data and cannot map participants to patient identifiers. These data are not incorporated into the HER; rather, EHR and genetic data are linked in a separate environment via a unique identifier as approved by the IRB.

### Processing of EHR data

Most cohort participants had multiple RBC recordings across thirteen years; in these cases, the mean age of each participant across those records was computed and later used as a covariate for each component in GWAS and PheWAS analyses. Normalization of test values was necessary as lab tests were updated across the 13 years of data collection. Many of the participants had lab results (for the same RBC component) recorded across different tests with different healthy reference ranges. For example, the 4,700 participants had measurements for MCV with respect to one or more of ten different MCV lab tests and corresponding healthy reference ranges. Many participants had records across several of these ten different tests. Only those tests/reference ranges having records for more than one individual were used in analyses. To standardize the RBC values across different normal reference ranges, a simple linear transform was computed using each test’s reference range and the most recent test’s range. All component measures within each separate test were then transformed into ranges of the most recent via each range’s specific linear transform. The most recent healthy normal reference range for each component is listed in Table 1. Distributions of raw and transformed laboratory test values can be found in Supplementary Fig S5(A-C).

### Genotyping and Quality Control

Genotyping was performed by 23andMe using the Illumina Infimum^®^ DNA Human OmniExpress-24 BeadChip V4. This genotyping platform (Illumina, San Diego, CA) consists of approximately 570,000 SNPs. DNA extraction and genotyping were performed on saliva samples by the National Genetics Institute (NG1), a CLIA licensed clinical laboratory and a subsidiary of the Laboratory Corporation of America.

Raw genotype data were processed through a standard quality control process [46,47,80-82]. SNPs with a minor allele frequency (MAF) less than 0.01 were removed. SNPS that were out of HWE (*p*-value < 1x10^-6^) were also excluded. Any SNP with call rate less than 95% was removed; any individual with a call rate less than 95% was also excluded from further study. Two pairs of participants were excluded due to high IBS (Identical by State) in all three cohorts). Additionally, twelve people were excluded due to high autosomal heterozygosity (FDR < 1%). This left 498,916 high-quality SNPS and 4,699 participants in the MCV cohort with mean autosomal heterozygosity of 0.321. The same process yielded 4,591 participants for MPV with the same mean autosomal heterozygosity of 0.321. Similarly, the PC cohort consisted of 4,700 participants with same mean autosomal heterozygosity.

Additionally, a principal component analysis (PCA) was performed to identify principal components to correct for population substructure. Genotype data was pruned to exclude SNPs with high linkage disequilibrium using *PLINK* and standard pruning parameters of 50 SNPs per sliding window; window size of five SNPs; *r*^2^=0.5 [80]. Regression models were adjusted by the first four components, decreasing the genomic inflation factor of all RBC components to λ≤ 1.04, well within standard ranges [17,27,83].

### GWAS

Using *PLINK* v1.9 [84], we performed a simple linear regression analysis with an assumed additive model (number of copies of the minor allele) including age, gender and the first four principal components as covariates to correct for any bias generated by these variables. Standardized values of all three components followed approximate normal distributions (Supplementary Fig S5(A-C) (row 2). Total phenotypic variance explained by the SNPs was calculated by first producing a genetic relationship matrix of all SNPs on autosomal chromosomes in *PLINK*. Subsequently, a restricted maximum likelihood analysis was conducted using GTCA on the relationship matrix to estimate the variance explained by the SNPS.

A simple one-way ANOVA was performed on the mean RBC component values across the three genotypes. The raw *p*-values associated to the F-test statistic are included in Supplementary Table S2. QUANTO [85] was used to calculate power in our study. We found that for every combination of a SNP’s effect size and its MAF, power was greater than 90% based on the approximately 4,500 participants used for each SNP’s analysis. Specifically, effect sizes ranged between [-0.62 11.49] (note that greater effect sizes are associated with greater mean component values) and MAFs ranged between [.073, 0.497]. These values are included in Table 2.

### PheWAS

The **R** package PheWAS [86] was used to perform two independent PheWAS analyses. The first examined associations between statistically significant SNPs identified in an RBC GWAS and EHR phenotypes based on ICD9 codes. The second identified associations between RBC levels in our cohort and ICD9-based diagnoses only. ICD9 and ICD10 codes for each individual in the cohort recorded in the Renown EHR were aggregated via a mapping from the Center for Medicare and Medicaid services (https://www.cms.gov/Medicare/Coding/ICD10/2018-ICD-10-CM-and-GEMs.html). A total of 34,555 individual diagnoses mapped to 6,632 documented ICD9 codes. ICD9 codes were aggregated and converted into 1,814 individual phenotype groups (“phecodes”) using the PheWAS package as described in Carroll and Denny [86,87]. Of these, only the phecodes that included at least 20 cases were used for downstream analyses, following Carroll’s protocol [86]: there were 1,488 phecodes with more than 20 cases in each PheWAS. Age, gender, and ethnicity were included in all PheWAS models. The first PheWAS detected associations between statistically significant SNPs (*p*<5x10^-8^) identified in each of the three GWASs above and case/control status of EHR phenotypes represented by ICD9 codes. Specifically, a logistic regression between the incidence (number of cases) of each phenotype group (phecode) and the additive genotypes of each statistically significant SNP was performed, using age and gender as covariates. Possible associations of 1,488 phecodes with each previously detected SNP were assessed. The level of statistical significance was computed as a Bonferroni correction for all possible associations per component: *p*=0.05/ *N_p_* /*N_s_*, where *N_p_* is the number of phecodes tested and *N_s_* is the number of SNPs examined in the specific blood component. This significance level is represented by a red line in Supplementary Fig S3(A-C).

A second PheWAS, as outlined in Carroll et al. (2014) [86], was performed to examine associations between each of the three quantitative RBC components and the phecode categories. Specifically, a linear regression between the RBC measure and the case/control status of a phecode was performed (with age and gender as covariates) for each of 1,488 phecodes. This analysis resulted in a number of hematologic phenotypes that associated with RBC component levels (Table 4). A single-SNP Bonferroni correction 3.4x10^-5^=0.05/*N_p_* (with *N_p_*=1,488) was used to compute the level of statistical significance. Phecodes with association levels *p*<3.4x10^-5^ are highlighted in Supplementary Fig S4 (A-C). Note that two associations with MPV at slightly higher *p*-values (*p*=5.43x10^-5^ and *p*=6.55x10^-5^) are also included; these are presented in the Discussion.

**Table 4.** PheWAS Results for MPV, MCV and PC.

| Phecode | Description | Group | RBC | $\beta$ | SE | $p$ | N |
| --- | --- | --- | --- | --- | --- | --- | --- |
| 287.3 | Thrombocytopenia | hematopoietic | MPV | 0.75 | 0.11 | $9.06 \times 10^{-12}$ | 4104 |
| 287 | Purpura and other hemorrhagic conditions | hematopoietic | MPV | 0.62 | 0.10 | $2.86 \times 10^{-10}$ | 4124 |
| 286.3 | Coagulation defects complicating pregnancy or postpartum | hematopoietic | MPV | 2.47 | 0.39 | $3.66 \times 10^{-10}$ | 4029 |
| 655 | Known or suspected fetal abnormality affecting mother | pregnancy complications | MPV | 0.56 | 0.11 | $8.85 \times 10^{-7}$ | 4455 |
| 798 | Malaise and fatigue | symptoms | MPV | 0.16 | 0.03 | $5.21 \times 10^{-6}$ | 4162 |
| 261 | Vitamin deficiency | endocrine/metabolic | MPV | 0.14 | 0.04 | $5.43 \times 10^{-5}$ | 4049 |
| 61.4 | Vitamin D deficiency | endocrine/metabolic | MPV | 0.14 | 0.04 | $6.55 \times 10^{-5}$ | 3992 |
| 282.8 | Other hemoglobinopathies | hematopoietic | MCV | -12.18 | 0.87 | $1.97 \times 10^{-43}$ | 3751 |
| 282 | Hereditary hemolytic anemias | hematopoietic | MCV | -10.33 | 0.82 | $8.62 \times 10^{-36}$ | 3754 |
| 280 | Iron deficiency anemias | hematopoietic | MCV | -3.76 | 0.36 | $1.30 \times 10^{-25}$ | 3854 |
| 278 | Overweight, obesity and other hyperalimentation | endocrine/metabolic | MCV | -1.48 | 0.14 | $3.56 \times 10^{-25}$ | 4365 |
| 278.1 | Obesity | endocrine/metabolic | MCV | -1.71 | 0.17 | $2.01 \times 10^{-23}$ | 3874 |
| 280.1 | Iron deficiency anemias unspecified or not due to blood loss | hematopoietic | MCV | -3.81 | 0.38 | $5.03 \times 10^{-23}$ | 3837 |
| 278.11 | Morbid obesity | endocrine/metabolic | MCV | -2.04 | 0.21 | $6.83 \times 10^{-22}$ | 3540 |
| 281.9 | Deficiency anemias | hematopoietic | MCV | 8.73 | 1.10 | $2.60 \times 10^{-15}$ | 3743 |
| 289.9 | Abnormality of red blood cells | hematopoietic | MCV | -7.30 | 1.08 | $1.94 \times 10^{-11}$ | 3742 |
| 289 | Other diseases of blood and blood-forming organs | hematopoietic | MCV | 2.84 | 0.43 | $7.37 \times 10^{-11}$ | 3827 |
| 317.11 | Alcoholic liver damage | mental disorders | MCV | 8.46 | 1.31 | $1.38 \times 10^{-10}$ | 4009 |
| 281 | Other deficiency anemia | hematopoietic | MCV | 4.55 | 0.73 | $4.33 \times 10^{-10}$ | 3759 |
| 317 | Alcohol-related disorders | mental disorders | MCV | 4.15 | 0.67 | $5.96 \times 10^{-10}$ | 4041 |
| 317.1 | Alcoholism | mental disorders | MCV | 5.59 | 0.91 | $9.92 \times 10^{-10}$ | 4021 |
| 571.8 | Liver abscess and sequelae of chronic liver disease | digestive | MCV | 8.01 | 1.50 | $9.85 \times 10^{-8}$ | 3855 |
| 571.51 | Cirrhosis of liver without mention of alcohol | digestive | MCV | 7.22 | 1.42 | $3.61 \times 10^{-7}$ | 3856 |
| 342 | Hemiplegia | neurological | MCV | -14.80 | 3.09 | $1.67 \times 10^{-6}$ | 4060 |
| 573.2 | Liver replaced by transplant | digestive | MCV | 14.01 | 2.99 | $2.97 \times 10^{-6}$ | 3849 |
| 571.81 | Portal hypertension | digestive | MCV | 9.80 | 2.12 | $4.00 \times 10^{-6}$ | 3851 |
| 250.4 | Abnormal glucose | endocrine/metabolic | MCV | -0.88 | 0.19 | $6.30 \times 10^{-6}$ | 3946 |
| 250 | Diabetes mellitus | endocrine/metabolic | MCV | -1.03 | 0.23 | $9.77 \times 10^{-6}$ | 3697 |
| 250.21 | Type 2 diabetes with ketoacidosis | endocrine/metabolic | MCV | 12.99 | 3.04 | $1.94 \times 10^{-5}$ | 3296 |
| 530.2 | Esophageal bleeding (varices/hemorrhage) | digestive | MCV | 8.18 | 1.91 | $1.96 \times 10^{-5}$ | 3176 |
| 70.2 | Viral hepatitis B | infectious diseases | MCV | -17.81 | 4.31 | $3.63 \times 10^{-5}$ | 4266 |
| 539 | Bariatric surgery | digestive | MCV | -1.98 | 0.48 | $3.71 \times 10^{-5}$ | 4550 |
| 287.3 | Thrombocytopenia | hematopoietic | PC | -85.15 | 6.56 | $9.17 \times 10^{-38}$ | 4104 |
| 287 | Purpura and other hemorrhagic conditions | hematopoietic | PC | -71.06 | 5.92 | $1.11 \times 10^{-32}$ | 4124 |
| 278 | Overweight, obesity and other hyperalimentation | endocrine/metabolic | PC | 14.23 | 1.97 | $6.12 \times 10^{-13}$ | 4366 |
| 284 | Aplastic anemia | hematopoietic | PC | -98.52 | 14.12 | $3.54 \times 10^{-12}$ | 3749 |
| 284.1 | Pancytopenia | hematopoietic | PC | -100.90 | 15.03 | $2.16 \times 10^{-11}$ | 3747 |
| 278.1 | Obesity | endocrine/metabolic | PC | 14.80 | 2.35 | $3.18 \times 10^{-10}$ | 3875 |
| 278.11 | Morbid obesity | endocrine/metabolic | PC | 17.70 | 2.92 | $1.56 \times 10^{-9}$ | 3541 |
| 571.51 | Cirrhosis of liver without mention of alcohol | digestive | PC | -120.03 | 19.85 | $1.61 \times 10^{-9}$ | 3857 |
| 571.8 | Liver abscess and sequelae of chronic liver disease | digestive | PC | -121.58 | 21.03 | $7.96 \times 10^{-9}$ | 3856 |
| 288.1 | Decreased white blood cell count | hematopoietic | PC | -32.91 | 5.82 | $1.68 \times 10^{-8}$ | 3832 |
| 287.31 | Primary thrombocytopenia | hematopoietic | PC | -134.74 | 25.98 | $2.24 \times 10^{-7}$ | 4028 |
| 286.3 | Coagulation defects complicating pregnancy or postpartum | hematopoietic | PC | -120.29 | 23.71 | $4.10 \times 10^{-7}$ | 4029 |
| 655 | Known or suspected fetal abnormality affecting mother | pregnancy complications | PC | -35.93 | 7.09 | $4.27 \times 10^{-7}$ | 4455 |
| 571.81 | Portal hypertension | digestive | PC | -143.14 | 29.73 | $1.53 \times 10^{-6}$ | 3852 |
| 288.2 | Elevated white blood cell count | hematopoietic | PC | 22.82 | 4.99 | $4.88 \times 10^{-6}$ | 3875 |
| 395.2 | Nonrheumatic aortic valve disorders | circulatory system | PC | -29.71 | 6.50 | $5.03 \times 10^{-6}$ | 4019 |
| 280.1 | Iron deficiency anemias unspecified or not due to blood loss | hematopoietic | PC | 24.29 | 5.74 | $2.36 \times 10^{-5}$ | 3838 |
| 555 | Inflammatory bowel disease and other gastroenteritis and colitis | digestive | PC | 33.57 | 7.99 | $2.70 \times 10^{-5}$ | 3504 |

## Data Availability Statement

### EHR Data

EHR data for the Healthy Nevada cohort are subject to HIPAA and other privacy and compliance restrictions. Mean standardized RBC component values for each individual are available in Supplementary Table S1.

### GWAS Results

To reduce the possibility of a privacy breach, 23andMe requires that the statistics for only 10,000 SNPs be made publicly available. This is the amount of data considered by 23andMe to be insufficient to enable a re-identification attack. The statistical summary results of the top 10,000 SNPs for the 23andMe data are available here: https://www.dri.edu/grzymski2020. All column definitions are listed in Table 5.

**Table 5.** Column Identifiers for GWAS Results.

| Column name | Definition |
| --- | --- |
| CHR | Chromosome |
| SNP | Individual SNP identifier |
| BP | Location of SNP on relative chromosome |
| A1 | Alternative Allele |
| TEST | Selected statistical test – ADD represents the additive effect |
| NMISS | Indicates the number of observations – non-missing genotypes |
| BETA | The effect size for this variant, defined per copy of the A1 allele |
| SE | The standard error of the effect size |
| LE | Lower end of the 95% confidence interval for the effect size |
| UE | Upper end of the 95% confidence interval for the effect size |
| STAT | The value of the test statistic |
| P | The p-value for the association test |

### PheWAS Results

Summarized counts of each ICD9 classification and phenotype group (phecode) are presented in Supplementary Table 3.

For more information please contact.

## Supporting information

## Acknowledgements

We thank Michele Henderson, Toni Curreri and all the ambassadors of the Healthy Nevada Project. We also thank Iva Neveux for her helpful discussions with phenotypic data. We thank Renown Health and DRI marketing and all the folks at 23andMe who helped launch the project. Research support was provided by the Governor’s Office of Economic Development Knowledge Fund. Support for the Healthy Nevada Project and personal genetics was provided by the Renown Health Foundation.

## Supplemental Figure and Table Legends

**Supplementary Table 1: Mean standardized RBC component values** This table includes mean standardized RBC component values for each individual along with age and gender. Due to the length of this table it can be found online at https://www.dri.edu/grzymski2020

**Supplementary Table 2: General SNP table for MPV, MCV and PC** This table lists the 38 statistically significant SNPs associated to MPV, MCV and PC in our cohort. General information about the SNP such as chromosome location, GWAS *p*-value, power, genotype, cytoband, ANOVA, and references of associations identified in previous studies are listed.

**Supplementary Table 3: Counts of each phecode group** This table presents the mapping between ICD9 codes and phecodes as presented in Carroll and the **R** package PheWAS [86] tested in our study, and the number of incidences from the RBC cohort in each phecode group.

**Supplementary Fig S1 (A, B, C): GWAS results for RBC components MPV, MCV and PC** Genome-wide association study results for the three RBC components. The x-axis represents the genomic position of 498,916 SNPs. The y-axis represents -log_10_-transformed raw *p*-values of each genotypic association. The red horizontal line indicates the threshold of significance *p*=5x10^-8^.

**Supplementary Fig 2: ANOVA results of SNP rs7961894** This figure shows the box and whisker diagram for standardized values of MPV of all members in the cohort based on genotype. Mean and standard deviation values for each genotype are CC: 10.54 ± 0.97; CT: 10.74 ± 1.0; TT: 11.21 ± 0.87. The *p*-value for this ANOVA analysis is *p*=8.7x10^-12^.

**Supplementary Fig S3 (A, B, C): PheWAS results between RBC component-significant SNPs and phecodes** These three figures show the results of individual logistic regressions between incidence of phenotype groups (phecodes) and SNP genotypes, based on the additive model. Models include age, gender and ethnicity as covariates. Each point represents the *p*-value of one SNP and one of 1,488 phecodes with at least 20 cases assigned to it. The horizontal red line in each represents the significance level *p*=1.60x10^-6^ for MPV, *p*=2.40x10^-6^ for MCV, and *p*=1.12x10^-5^ for PC.

**Supplementary Fig S4 (A, B, C): PheWAS results between RBC component and phecodes** These three figures show the results of individual linear regressions between incidence of phenotype groups (phecodes) and continuous RBC component measures. Models include age, gender and ethnicity as covariates. Each point represents the *p*-value of the association between one of 1,488 phecodes with at least 20 cases assigned to it, and the RBC component measure. The horizontal red line in each represents the significance level *p*=1.60x10^-6^ for MPV, *p*=2.40x10^-6^ for MCV, and *p*=1.12x10^-5^ for PC.

**Supplementary Fig S5 (A, B, C): Raw and Standardized RBC Component Lab Measures** Distribution of raw RBC component values are presented in the first row; distribution of component values upon standardization to the most recent lab test are shown in the second row; the QQ-plot of the standardized values is pictured in the third row.

## Author contributions

KAS and RWR conducted genetic and clinical data analysis and wrote the manuscript. GE, ADS and KAS contributed to the clinical discussion. WJM extracted participants and their clinical health data from the Renown EHR. RA and the 23andMe research team provided participant genotype data and edited the manuscript. JJG conceived of and obtained funds to conduct this experiment. All authors reviewed, edited and approved the final version of the manuscript.

