## Supplementary figures and images for "GWAS and PheWAS of Red Blood Cell Components in a Northern Nevadan Cohort"

### Supplementary file 1

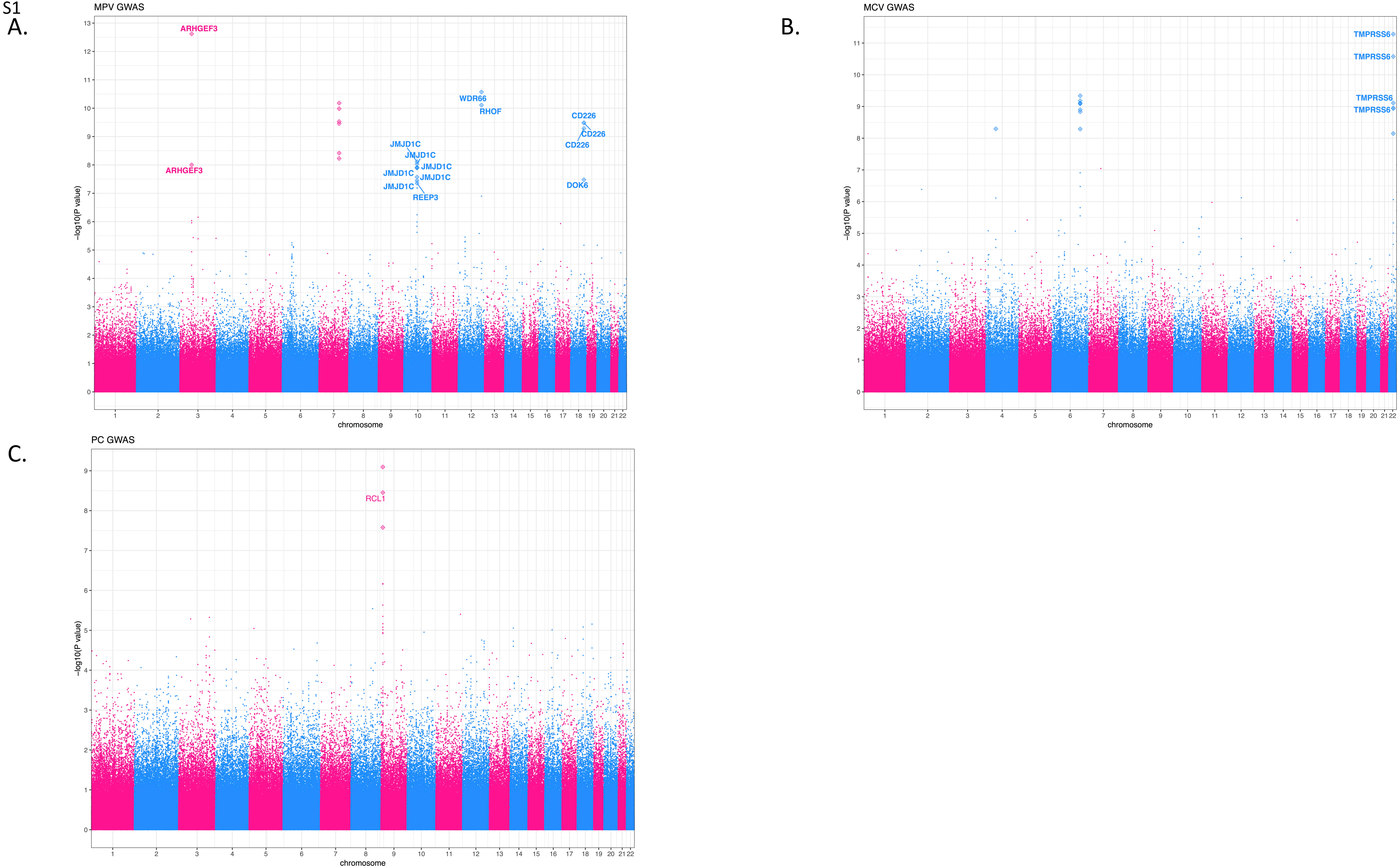

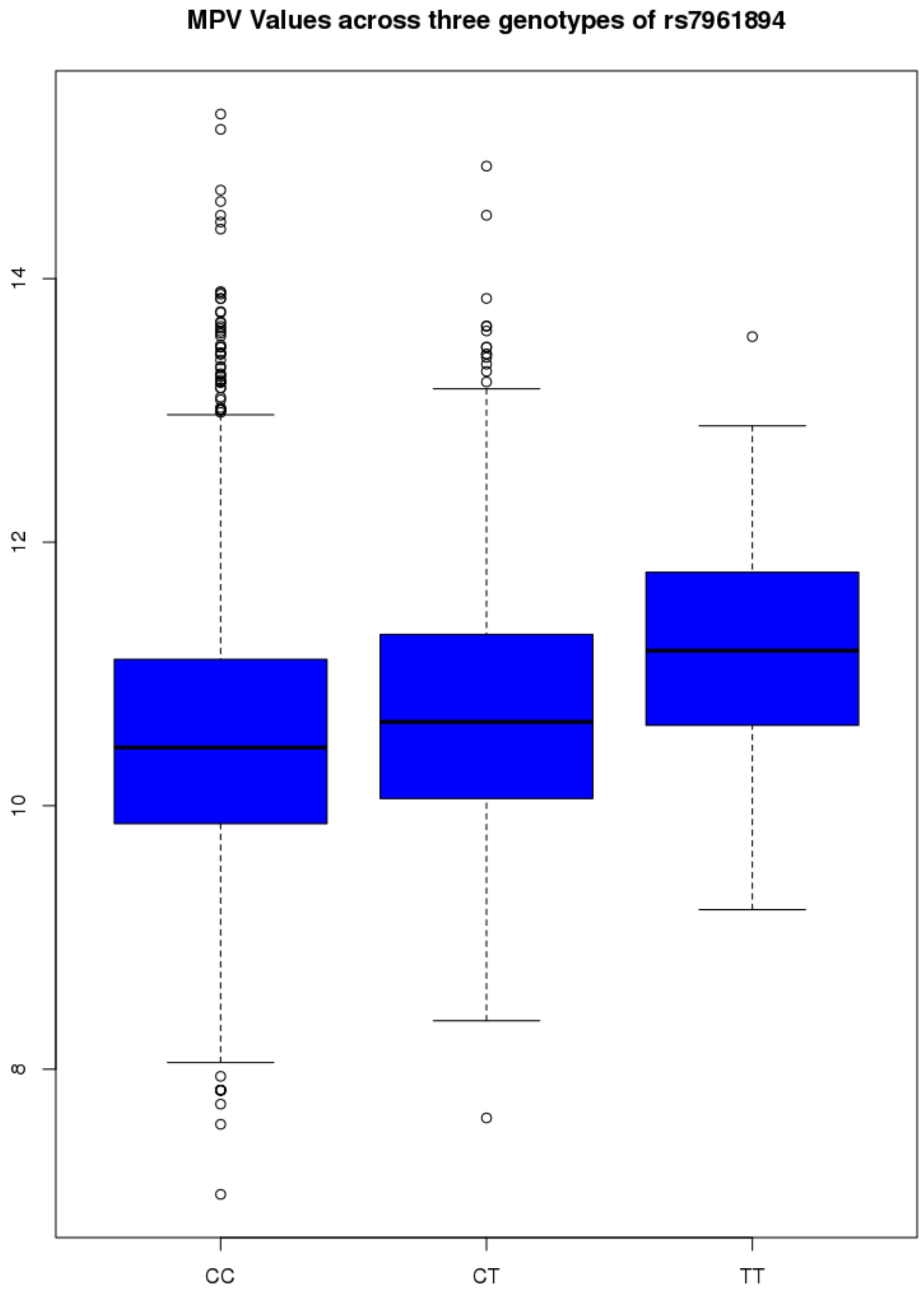

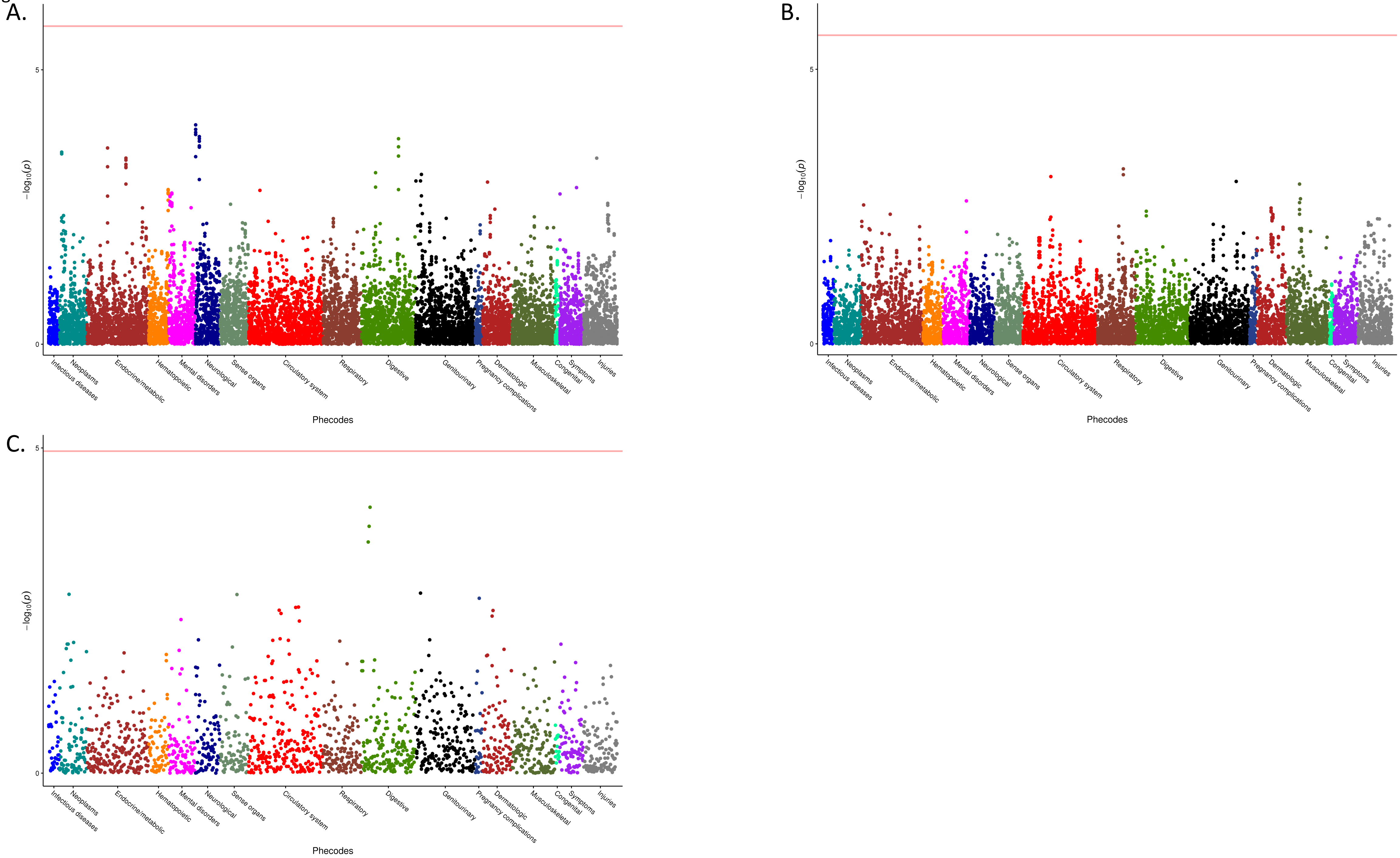

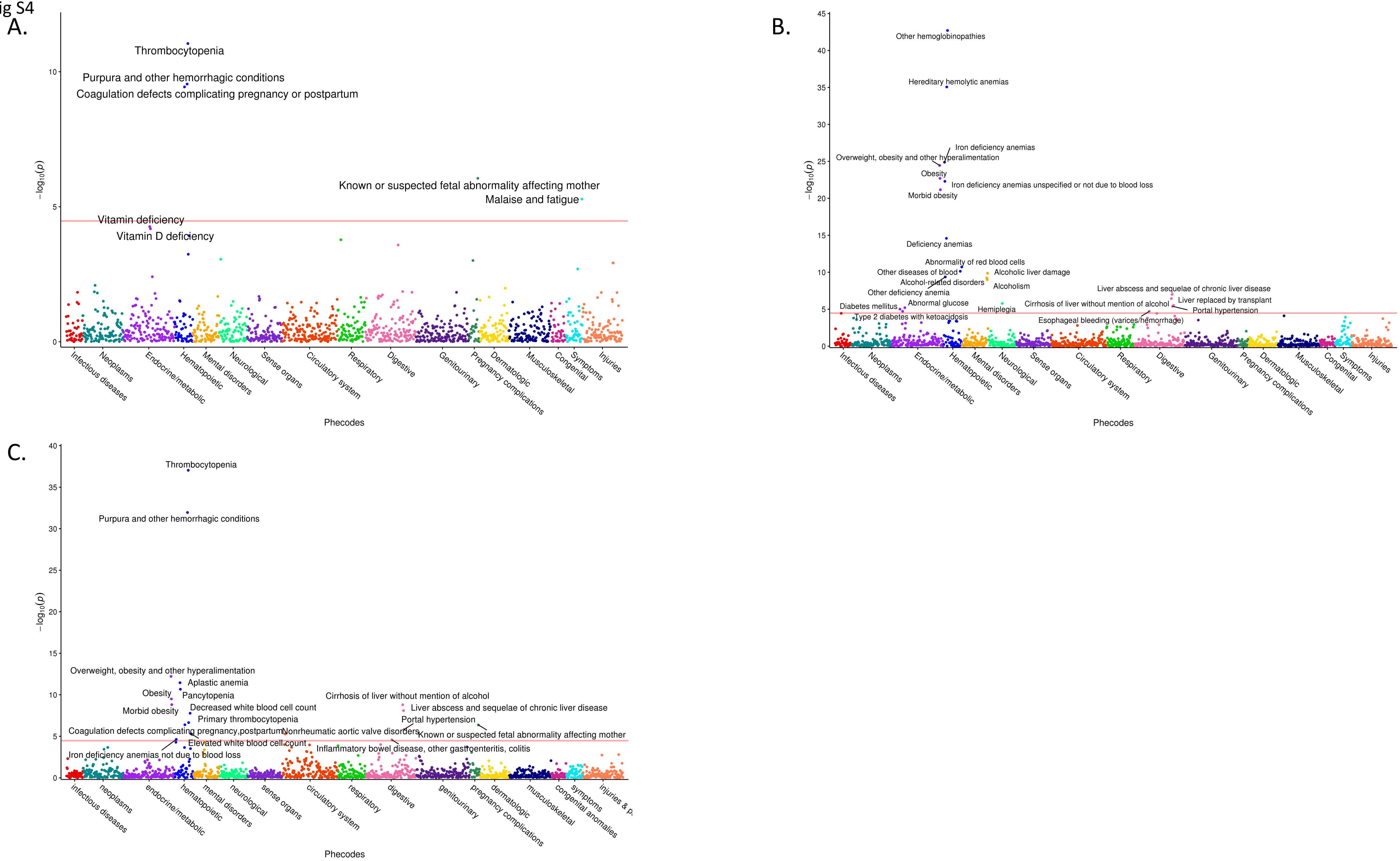

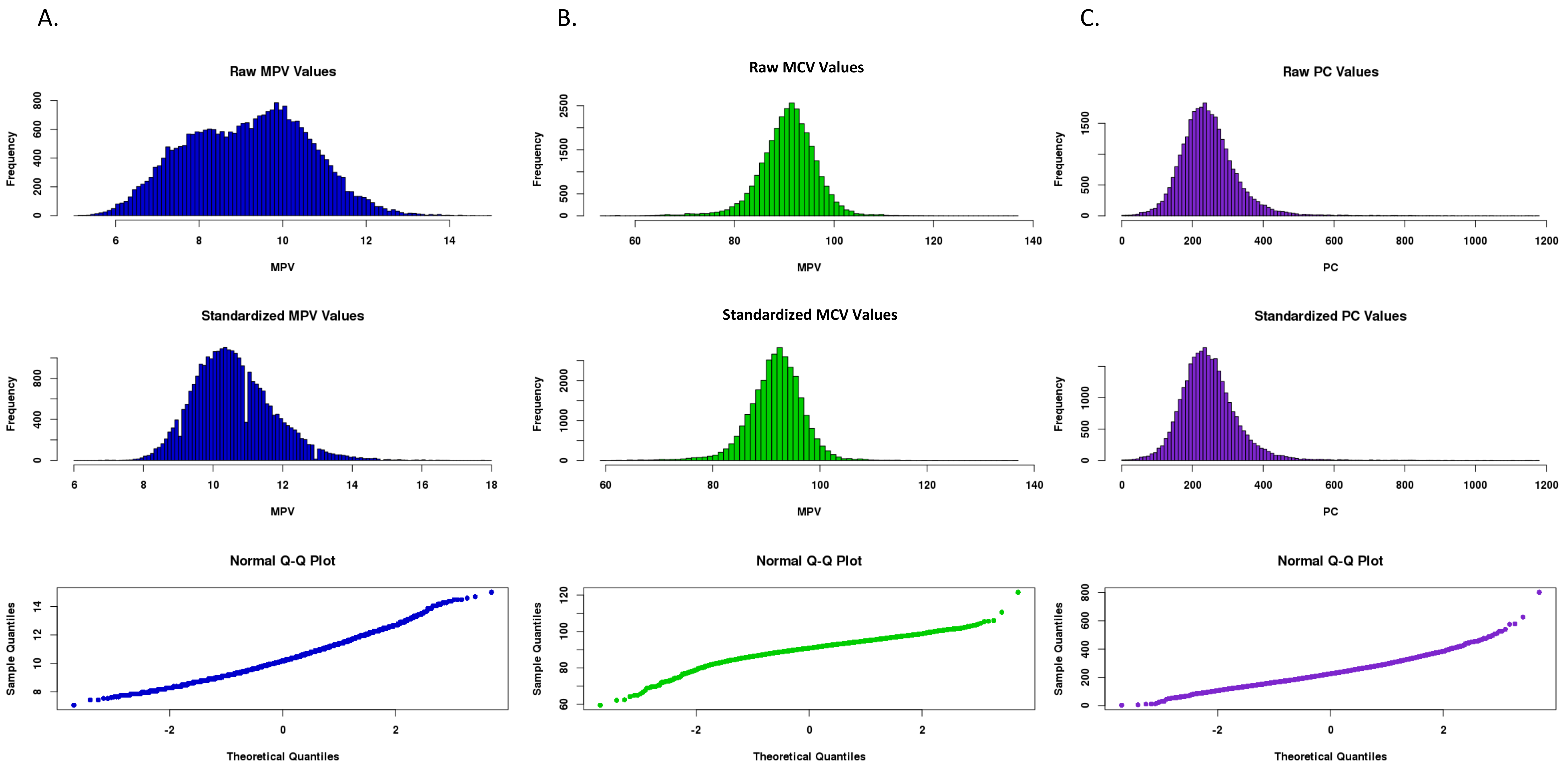
