## Supplementary material for "GWAS and PheWAS of Red Blood Cell Components in a Northern Nevadan Cohort"

| rsID | N | GWAS *p*-value | power of GWAS | Genotype 1 | Genotype 2 | Genotype 3 | Mean +/- SD Genotype 1 | Mean +/- SD Genotype 2 | Mean +/- SD Genotype 3 | ANOVA *p*-value | Chromosome | Cytoband Region | SNP Previously Identified | Cytoband Region Referenced | RBC Component |
|---|---|---|---|---|---|---|---|---|---|---|---|---|---|---|---|
| rs10274553 | 4589 | 3.82x10$^{-9}$ | >90 | T/T | T/C | C/C | 10.72 +/- 0.99 | 10.57 +/- 0.98 | 10.46 +/- 0.94 | 4.50x10$^{-10}$ | chr3 | p14.3 | [13,18,28,33] | [18,28] | MPV |
| rs10509186 | 4589 | 7.75x10$^{-9}$ | >90 | C/C | C/T | T/T | 10.69 +/- 1 | 10.56 +/- 0.97 | 10.47 +/- 0.95 | 2.61x10$^{-7}$ | chr3 | p14.3 | N/A | [18,28] | MPV |
| rs10822186 | 4591 | 4.38x10$^{-8}$ | >90 | A/A | A/G | G/G | 10.69 +/- 0.99 | 10.57 +/- 0.97 | 10.49 +/- 0.96 | 3.13x10$^{-6}$ | chr7 | q22.3 | [13,17,25,28,30,35 | [13,25,59] | MPV |
| rs11130549 | 4589 | 9.94x10$^{-9}$ | >90 | T/T | T/C | C/C | 10.66 +/- 0.99 | 10.55 +/- 0.97 | 10.44 +/- 0.93 | 2.22x10$^{-6}$ | chr7 | q22.3 | [35] | [13,25,59] | MPV |
| rs12355784 | 4578 | 9.32x10$^{-9}$ | >90 | C/C | C/A | A/A | 10.69 +/- 1 | 10.56 +/- 0.97 | 10.48 +/- 0.96 | 7.42x10$^{-7}$ | chr7 | q22.3 | [13,28] | [13,25,59] | MPV |
| rs1354034 | 4583 | 2.39x10$^{-13}$ | >90 | C/C | C/T | T/T | 10.46 +/- 0.94 | 10.6 +/- 0.97 | 10.79 +/- 1.03 | 2.40x10$^{-13}$ | chr7 | q22.3 | [18,30] | [13,25,59] | MPV |
| rs1788103 | 4591 | 5.15x10$^{-10}$ | >90 | A/A | A/G | G/G | 10.69 +/- 1.01 | 10.58 +/- 0.97 | 10.45 +/- 0.95 | 2.59x10$^{-8}$ | chr7 | q22.3 | N/A | [13,25,59] | MPV |
| rs1790588 | 4590 | 3.31x10$^{-10}$ | >90 | T/T | T/C | C/C | 10.69 +/- 1.01 | 10.58 +/- 0.97 | 10.45 +/- 0.94 | 1.70x10$^{-8}$ | chr7 | q22.3 | N/A | [13,25,59] | MPV |
| rs1790974 | 4590 | 3.32x10$^{-8}$ | >90 | C/C | C/T | T/T | 10.67 +/- 1.02 | 10.57 +/- 0.96 | 10.46 +/- 0.93 | 1.14x10$^{-6}$ | chr10 | q21.3 | N/A | [18,30,49] | MPV |
| rs1935 | 4585 | 3.57x10$^{-8}$ | >90 | C/C | C/G | G/G | 10.68 +/- 1 | 10.57 +/- 0.97 | 10.48 +/- 0.95 | 2.53x10$^{-6}$ | chr10 | q21.3 | [49,50] | [18,30,49] | MPV |
| rs201979226 | 4578 | 5.89x10$^{-9}$ | >90 | T/T | T/C | C/C | 10.47 +/- 0.93 | 10.57 +/- 0.99 | 10.73 +/- 0.99 | 4.82x10$^{-10}$ | chr10 | q21.3 | [51,52,53] | [18,30,49] | MPV |
| rs342240 | 4590 | 3.49x10$^{-10}$ | >90 | G/G | G/A | A/A | 10.48 +/- 0.95 | 10.59 +/- 0.98 | 10.76 +/- 1 | 8.26x10$^{-11}$ | chr10 | q21.3 | [18,33] | [18,30,49] | MPV |
| rs342275 | 4591 | 2.96x10$^{-10}$ | >90 | C/C | C/T | T/T | 10.48 +/- 0.94 | 10.59 +/- 0.98 | 10.77 +/- 1 | 6.45x10$^{-11}$ | chr10 | q21.3 | [54,55] | [18,30,49] | MPV |
| rs342293 | 4586 | 6.61x10$^{-11}$ | >90 | C/C | C/G | G/G | 10.48 +/- 0.93 | 10.57 +/- 0.99 | 10.77 +/- 0.99 | 5.44x10$^{-12}$ | chr10 | q21.3 | [35,56] | [18,30,49] | MPV |
| rs342296 | 4588 | 1.04x10$^{-10}$ | >90 | G/G | G/A | A/A | 10.47 +/- 0.93 | 10.58 +/- 0.99 | 10.77 +/- 0.98 | 1.26x10$^{-11}$ | chr10 | q21.3 | [53] | [18,30,49] | MPV |
| rs34818942 | 4582 | 7.77x10$^{-11}$ | >90 | C/C | C/T | T/T | 10.55 +/- 0.97 | 10.79 +/- 1.01 | 11.26 +/- 0.67 | 7.90x10$^{-11}$ | chr12 | q24.31 | [13,18,24,28] | [18,60] | MPV |
| rs386614085 | 4591 | 1.21x10$^{-8}$ | >90 | A/A | A/G | G/G | 10.69 +/- 1 | 10.56 +/- 0.97 | 10.48 +/- 0.96 | 9.58x10$^{-7}$ | chr12 | q24.31 | N/A | [18,60] | MPV |
| rs4379723 | 4582 | 1.29x10$^{-8}$ | >90 | T/T | T/C | C/C | 10.68 +/- 1 | 10.57 +/- 0.97 | 10.47 +/- 0.95 | 1.01x10$^{-6}$ | chr18 | q22.2 | [57] | [17] | MPV |
| rs763361 | 4591 | 3.26x10$^{-10}$ | >90 | C/C | C/T | T/T | 10.7 +/- 1.01 | 10.58 +/- 0.97 | 10.45 +/- 0.94 | 1.48x10$^{-8}$ | chr18 | q22.2 | [58] | [17] | MPV |
| rs7910927 | 4575 | 2.68x10$^{-8}$ | >90 | T/T | T/G | G/G | 10.68 +/- 1 | 10.56 +/- 0.97 | 10.48 +/- 0.95 | 1.74x10$^{-6}$ | chr18 | q22.2 | N/A | [17] | MPV |
| rs7961894 | 4591 | 2.68x10$^{-11}$ | >90 | C/C | C/T | T/T | 10.54 +/- 0.97 | 10.74 +/- 1 | 11.21 +/- 0.87 | 8.70x10$^{-12}$ | chr18 | q22.2 | N/A | [17] | MPV |
| rs218237 | 4699 | 5.07x10$^{-9}$ | >90% | C/C | C/T | T/T | 91.4 +/- 4.47 | 91.84 +/- 4.41 | 91.92 +/- 4.86 | 8.40x10$^{-3}$ | chr4 | q12 | [45] | [20,27,32] | MCV |
| rs9402686 | 4697 | 4.60x10x$^{-10}$ | >90% | G/G | G/A | A/A | 91.13 +/- 4.4 | 91.94 +/- 4.46 | 92.65 +/- 4.76 | 1.88x10$^{-12}$ | chr6 | q23.3 | [17,27,40] | [20,27,32] | MCV |
| rs7776054 | 4699 | 6.65x10$^{-10}$ | >90% | A/A | A/G | G/G | 91.17 +/- 4.41 | 91.88 +/- 4.48 | 92.81 +/- 4.54 | 7.85x10$^{-12}$ | chr6 | q23.3 | [21,23,39,42,44] | [20,27,32] | MCV |
| rs9399137 | 4694 | 7.82x10$^{-10}$ | >90% | T/T | T/C | C/C | 91.15 +/- 4.45 | 91.93 +/- 4.42 | 92.81 +/- 4.53 | 9.76x10$^{-13}$ | chr6 | q23.3 | [20,21,28,42] | [20,27,32] | MCV |
| rs7775698 | 4696 | 8.24x10$^{-10}$ | >90% | C/C | C/T | T/T | 91.17 +/- 4.41 | 91.88 +/- 4.48 | 92.79 +/- 4.55 | 1.12x10$^{-11}$ | chr6 | q23.3 | [26,27,33] | [20,27,32] | MCV |
| rs4895441 | 4699 | 1.27x10$^{-9}$ | >90% | A/A | A/G | G/G | 91.13 +/- 4.4 | 91.94 +/- 4.45 | 92.57 +/- 4.76 | 5.77x10$^{-12}$ | chr6 | q23.3 | [20,23,27,37] | [20,27,32] | MCV |
| rs111194878 | 4697 | 1.47x10$^{-9}$ | >90% | C/C | C/A | A/A | 91.15 +/- 4.4 | 91.89 +/- 4.46 | 92.61 +/- 4.77 | 4.13x10$^{-11}$ | chr6 | q23.3 | [20,27,41] | [20,27,32] | MCV |
| rs9373124 | 4681 | 5.14x10$^{-9}$ | >90% | T/T | T/C | C/C | 91.19 +/- 4.36 | 91.84 +/- 4.51 | 92.49 +/- 4.91 | 8.46x10$^{-9}$ | chr6 | q23.3 | [29,43] | [20,27,32] | MCV |
| rs855791 | 4697 | 5.23x10$^{-12}$ | >90% | G/G | G/A | A/A | 92.09 +/- 4.44 | 91.47 +/- 4.48 | 90.77 +/- 4.36 | 1.60x10$^{-11}$ | chr22 | q12.3 | [27,31,33,36,38] | [27,32] | MCV |
| rs4820268 | 4643 | 2.65x10$^{-11}$ | >90% | A/A | A/G | G/G | 92.08 +/- 4.53 | 91.54 +/- 4.44 | 90.76 +/- 4.4 | 2.03x10$^{-11}$ | chr22 | q12.3 | [22,27,31] | [27,32] | MCV |
| rs5756504 | 4699 | 7.77x10$^{-10}$ | >90% | C/C | C/T | T/T | 91.03 +/- 4.38 | 91.75 +/- 4.47 | 92.19 +/- 4.57 | 5.02x10$^{-10}$ | chr22 | q12.3 | [26,27,45] | [27,32] | MCV |
| rs130624 | 4681 | 1.13x10$^{-9}$ | >90% | T/T | T/G | G/G | 91.01 +/- 4.28 | 91.65 +/- 4.58 | 92.14 +/- 4.42 | 3.27x10$^{-9}$ | chr22 | q12.3 | [32] | [27,32] | MCV |
| rs5756506 | 4618 | 1.15x10$^{-9}$ | >90% | G/G | G/C | C/C | 91.05 +/- 4.38 | 91.74 +/- 4.5 | 92.18 +/- 4.55 | 1.77x10$^{-9}$ | chr22 | q12.3 | [17,27] | [27,32] | MCV |
| rs386563505 | 4698 | 7.12x10$^{-9}$ | >90% | G/G | G/A | A/A | 91.09 +/- 4.35 | 91.62 +/- 4.53 | 92.16 +/- 4.46 | 6.54x10$^{-8}$ | chr22 | q12.3 | N/A | [27,32] | MCV |
| rs385893 | 4699 | 8.04x10$^{-10}$ | >90% | C/C | C/T | T/T | 260.83 +/- 65.81 | 250.23 +/- 61.86 | 244.26 +/- 57.71 | 2.58x10$^{-10}$ | chr9 | p24.1 | [17,25,26,34] | [17,18,25] | PC |
| rs10974808 | 4691 | 3.53x10$^{-9}$ | >90% | A/A | A/G | G/G | 249.15 +/- 61.55 | 259.41 +/- 62.62 | 271.78 +/- 76.72 | 7.39x10$^{-7}$ | chr9 | p24.1 | N/A | [17,18,25] | PC |
| rs423955 | 4699 | 2.64x10$^{-8}$ | >90% | T/T | T/C | C/C | 256.85 +/- 62.51 | 249.14 +/- 63.06 | 241.53 +/- 55.74 | 1.12x10$^{-7}$ | chr9 | p24.1 | [18] | [17,18,25] | PC |
