## Supplementary material for "GWAS and PheWAS of Red Blood Cell Components in a Northern Nevadan Cohort"

| Phecode | Description | Phenotype Group | Phenotype Group Description | PC Counts | MPV Counts | MCV Counts |
|---|---|---|---|---|---|---|
| 8 | Intestinal infection | 1 | infectious diseases | 3 | 3 | 3 |
| 8.5 | Bacterial enteritis | 1 | infectious diseases | 0 | 0 | 0 |
| 8.51 | Intestinal e.coli | 1 | infectious diseases | 0 | 0 | 0 |
| 8.52 | Intestinal infection due to C. difficile | 1 | infectious diseases | 0 | 0 | 0 |
| 8.6 | Viral Enteritis | 1 | infectious diseases | 0 | 0 | 0 |
| 8.7 | Intestinal infection due to protozoa | 1 | infectious diseases | 0 | 0 | 0 |
| 10 | Tuberculosis | 1 | infectious diseases | 1 | 1 | 1 |
| 31 | Diseases due to other mycobacteria | 1 | infectious diseases | 0 | 0 | 0 |
| 31.1 | Leprosy | 1 | infectious diseases | 0 | 0 | 0 |
| 38 | Septicemia | 1 | infectious diseases | 32 | 32 | 32 |
| 38.1 | Gram negative septicemia | 1 | infectious diseases | 0 | 0 | 0 |
| 38.2 | Gram positive septicemia | 1 | infectious diseases | 1 | 1 | 1 |
| 38.3 | Bacteremia | 1 | infectious diseases | 2 | 2 | 2 |
| 41 | Bacterial infection NOS | 1 | infectious diseases | 219 | 219 | 219 |
| 41.1 | Staphylococcus infections | 1 | infectious diseases | 24 | 24 | 24 |
| 41.11 | Methicillin sensitive Staphylococcus aureus | 1 | infectious diseases | 1 | 1 | 1 |
| 41.12 | Methicillin resistant Staphylococcus aureus | 1 | infectious diseases | 14 | 14 | 14 |
| 41.2 | Streptococcus infection | 1 | infectious diseases | 16 | 16 | 16 |
| 41.21 | Rheumatic fever / chorea | 1 | infectious diseases | 3 | 3 | 3 |
| 41.4 | E. coli | 1 | infectious diseases | 22 | 22 | 22 |
| 41.8 | H. pylori | 1 | infectious diseases | 6 | 6 | 6 |
| 41.9 | Infection with drug-resistant microorganisms | 1 | infectious diseases | 4 | 4 | 4 |
| 53 | Herpes zoster | 1 | infectious diseases | 0 | 0 | 0 |
| 53.1 | Herpes zoster with nervous system complications | 1 | infectious diseases | 0 | 0 | 0 |
| 54 | Herpes simplex | 1 | infectious diseases | 7 | 7 | 7 |
| 70 | Viral hepatitis | 1 | infectious diseases | 6 | 6 | 6 |
| 70.1 | Viral hepatitis A | 1 | infectious diseases | 0 | 0 | 0 |
| 70.2 | Viral hepatitis B | 1 | infectious diseases | 1 | 1 | 1 |
| 70.3 | Viral hepatitis C | 1 | infectious diseases | 4 | 4 | 4 |
| 70.4 | Chronic hepatitis | 1 | infectious diseases | 6 | 6 | 6 |
| 70.9 | Hepatitis NOS | 1 | infectious diseases | 7 | 7 | 7 |
| 71 | Human immunodeficiency virus [HIV] disease | 1 | infectious diseases | 8 | 8 | 8 |
| 71.1 | HIV infection, symptomatic | 1 | infectious diseases | 1 | 1 | 1 |
| 78 | Viral warts & HPV | 1 | infectious diseases | 10 | 10 | 10 |
| 79 | Viral infection | 1 | infectious diseases | 22 | 22 | 22 |
| 79.1 | Varicella infection | 1 | infectious diseases | 0 | 0 | 0 |
| 79.2 | Infectious mononucleosis | 1 | infectious diseases | 3 | 3 | 3 |
| 79.9 | Viremia, NOS | 1 | infectious diseases | 1 | 1 | 1 |
| 80 | Postoperative infection | 1 | infectious diseases | 17 | 17 | 17 |
| 81 | Infection/inflammation of internal prosthetic device; implant; and graft | 1 | infectious diseases | 9 | 9 | 9 |
| 81.1 | Graft-versus-host disease | 1 | infectious diseases | 0 | 0 | 0 |
| 81.11 | Acute graft-versus-host disease | 1 | infectious diseases | 0 | 0 | 0 |
| 81.12 | Chronic graft-versus-host disease | 1 | infectious diseases | 0 | 0 | 0 |
| 90 | Sexually transmitted infections (not HIV or hepatitis) | 1 | infectious diseases | 1 | 1 | 1 |
| 90.2 | Gonococcal infections | 1 | infectious diseases | 0 | 0 | 0 |
| 90.3 | Venereal diseases due to Chlamydia trachomatis | 1 | infectious diseases | 0 | 0 | 0 |
| 110 | Dermatophytosis / Dermatomycosis | 1 | infectious diseases | 122 | 122 | 122 |
| 110.1 | Dermatophytosis | 1 | infectious diseases | 104 | 104 | 104 |
| 110.11 | Dermatophytosis of nail | 1 | infectious diseases | 57 | 57 | 57 |
| 110.12 | Althete's foot | 1 | infectious diseases | 13 | 13 | 13 |
| 110.13 | Dermatophytosis of the body | 1 | infectious diseases | 24 | 24 | 24 |
| 110.2 | Dermatomycoses | 1 | infectious diseases | 14 | 14 | 14 |
| 112 | Candidiasis | 1 | infectious diseases | 139 | 139 | 139 |
| 112.3 | Candidiasis of skin and nails | 1 | infectious diseases | 22 | 22 | 22 |
| 117 | Mycoses | 1 | infectious diseases | 9 | 9 | 9 |
| 117.1 | Histoplasmosis | 1 | infectious diseases | 0 | 0 | 0 |
| 117.2 | Coccidioidomycosis | 1 | infectious diseases | 3 | 3 | 3 |
| 117.3 | Blastomycotic infection | 1 | infectious diseases | 0 | 0 | 0 |
| 117.4 | Aspergillosis | 1 | infectious diseases | 0 | 0 | 0 |
| 130 | Spirochetal infection | 1 | infectious diseases | 0 | 0 | 0 |
| 130.1 | Lyme disease | 1 | infectious diseases | 0 | 0 | 0 |
| 131 | Protozoan infection | 1 | infectious diseases | 3 | 3 | 3 |
| 132 | Infestation (lice, mites) | 1 | infectious diseases | 5 | 5 | 5 |
| 132.1 | Pediculosis and phthirus infestation | 1 | infectious diseases | 1 | 1 | 1 |
| 133 | Arthropod-borne diseases | 1 | infectious diseases | 0 | 0 | 0 |

| | | | | | | |
|---|---|---|---|---|---|---|
| 134 | Helminthiases | 1 | infectious diseases | 0 | 0 | 0 |
| 134.1 | Intestinal helminthiases | 1 | infectious diseases | 0 | 0 | 0 |
| 136 | Other infectious and parasitic diseases | 1 | infectious diseases | 33 | 33 | 33 |
| 145 | Cancer of mouth | 2 | neoplasms | 1 | 1 | 1 |
| 145.1 | Cancer of lip | 2 | neoplasms | 0 | 0 | 0 |
| 145.2 | Cancer of tongue | 2 | neoplasms | 1 | 1 | 1 |
| 145.3 | Cancer of major salivary glands | 2 | neoplasms | 0 | 0 | 0 |
| 145.4 | Cancer of the gums | 2 | neoplasms | 0 | 0 | 0 |
| 145.5 | Cancer of the mouth floor | 2 | neoplasms | 0 | 0 | 0 |
| 149 | Cancer of larynx, pharynx, nasal cavities | 2 | neoplasms | 0 | 0 | 0 |
| 149.1 | Cancer of oropharynx | 2 | neoplasms | 0 | 0 | 0 |
| 149.2 | Cancer of nasopharynx | 2 | neoplasms | 0 | 0 | 0 |
| 149.3 | Cancer of hypopharynx | 2 | neoplasms | 0 | 0 | 0 |
| 149.4 | Cancer of larynx | 2 | neoplasms | 0 | 0 | 0 |
| 149.5 | Hx of malignant neoplasm of oral cavity and pharynx | 2 | neoplasms | 0 | 0 | 0 |
| 149.9 | Cancer of of nasal cavities | 2 | neoplasms | 0 | 0 | 0 |
| 150 | Cancer of esophagus | 2 | neoplasms | 2 | 2 | 2 |
| 151 | Cancer of stomach | 2 | neoplasms | 1 | 1 | 1 |
| 153 | Colorectal cancer | 2 | neoplasms | 25 | 25 | 25 |
| 153.2 | Colon cancer | 2 | neoplasms | 22 | 22 | 22 |
| 153.3 | Malignant neoplasm of rectum, rectosigmoid junction, and anus | 2 | neoplasms | 7 | 7 | 7 |
| 155 | Cancer of liver and intrahepatic bile duct | 2 | neoplasms | 5 | 5 | 5 |
| 155.1 | Malignant neoplasm of liver, primary | 2 | neoplasms | 2 | 2 | 2 |
| 157 | Pancreatic cancer | 2 | neoplasms | 4 | 4 | 4 |
| 158 | Neoplasm of unspecified nature of digestive system | 2 | neoplasms | 4 | 4 | 4 |
| 159 | Malignant neoplasm of other and ill-defined sites within the digestive organs and peritoneum | 2 | neoplasms | 6 | 6 | 6 |
| 159.2 | Malignant neoplasm of small intestine, including duodenum | 2 | neoplasms | 1 | 1 | 1 |
| 159.3 | Malignant neoplasm of gallbladder and extrahepatic bile ducts | 2 | neoplasms | 1 | 1 | 1 |
| 159.4 | Malignant neoplasm of retroperitoneum and peritoneum | 2 | neoplasms | 0 | 0 | 0 |
| 164 | Cancer of intrathoracic organs | 2 | neoplasms | 0 | 0 | 0 |
| 165 | Cancer within the respiratory system | 2 | neoplasms | 11 | 11 | 11 |
| 165.1 | Cancer of bronchus; lung | 2 | neoplasms | 10 | 10 | 10 |
| 170 | Cancer of bone and connective tissue | 2 | neoplasms | 4 | 4 | 4 |
| 170.1 | Bone cancer | 2 | neoplasms | 0 | 0 | 0 |
| 170.2 | Cancer of connective tissue | 2 | neoplasms | 3 | 3 | 3 |
| 172 | Skin cancer | 2 | neoplasms | 145 | 145 | 145 |
| 172.1 | Melanomas of skin, dx or hx | 2 | neoplasms | 34 | 34 | 34 |
| 172.11 | Melanomas of skin | 2 | neoplasms | 18 | 18 | 18 |
| 172.2 | Other non-epithelial cancer of skin | 2 | neoplasms | 109 | 109 | 109 |
| 172.21 | Basal cell carcinoma (new) | 2 | neoplasms | 29 | 29 | 29 |
| 172.22 | Squamous cell carcinoma | 2 | neoplasms | 5 | 5 | 5 |
| 172.3 | Carcinoma in situ of skin | 2 | neoplasms | 1 | 1 | 1 |
| 173 | Neoplasm of uncertain behavior of skin | 2 | neoplasms | 35 | 35 | 35 |
| 174 | Breast cancer | 2 | neoplasms | 157 | 157 | 157 |
| 174.1 | Breast cancer [female] | 2 | neoplasms | 155 | 155 | 155 |
| 174.11 | Malignant neoplasm of female breast | 2 | neoplasms | 155 | 155 | 155 |
| 174.2 | Breast cancer [male] | 2 | neoplasms | 0 | 0 | 0 |
| 174.3 | Neoplasm of uncertain behavior of breast | 2 | neoplasms | 1 | 1 | 1 |
| 175 | Acquired absence of breast | 2 | neoplasms | 37 | 37 | 37 |
| 180 | Cervical cancer and dysplasia | 2 | neoplasms | 20 | 20 | 20 |
| 180.1 | Cervical cancer | 2 | neoplasms | 10 | 10 | 10 |
| 180.3 | Cervical intraepithelial neoplasia [CIN] [Cervical dysplasia] | 2 | neoplasms | 10 | 10 | 10 |
| 182 | Malignant neoplasm of uterus | 2 | neoplasms | 18 | 18 | 18 |
| 184 | Cancer of other female genital organs | 2 | neoplasms | 13 | 13 | 13 |
| 184.1 | Malignant neoplasm of ovary and other uterine adnexa | 2 | neoplasms | 10 | 10 | 10 |
| 184.11 | Malignant neoplasm of ovary | 2 | neoplasms | 9 | 9 | 9 |
| 184.2 | Cancer of other female genital organs | 2 | neoplasms | 2 | 2 | 2 |
| 185 | Cancer of prostate | 2 | neoplasms | 52 | 52 | 52 |
| 187 | Cancer of other male genital organs | 2 | neoplasms | 3 | 3 | 3 |
| 187.1 | Malignant neoplasm of unspecified male genital organ | 2 | neoplasms | 1 | 1 | 1 |
| 187.2 | Malignant neoplasm of testis | 2 | neoplasms | 2 | 2 | 2 |
| 187.8 | Neoplasm of uncertain behavior of male genital organs | 2 | neoplasms | 0 | 0 | 0 |
| 189 | Cancer of urinary organs (incl. kidney and bladder) | 2 | neoplasms | 27 | 27 | 27 |
| 189.1 | Cancer of kidney and renal pelvis | 2 | neoplasms | 17 | 17 | 17 |
| 189.11 | Malignant neoplasm of kidney, except pelvis | 2 | neoplasms | 4 | 4 | 4 |
| 189.12 | Malignant neoplasm of renal pelvis | 2 | neoplasms | 1 | 1 | 1 |
| 189.2 | Cancer of bladder | 2 | neoplasms | 9 | 9 | 9 |

| | | | | | | |
|---|---|---|---|---|---|---|
| 189.21 | Malignant neoplasm of bladder | 2 | neoplasms | 7 | 7 | 7 |
| 189.4 | Malignant neoplasm of other urinary organs | 2 | neoplasms | 9 | 9 | 9 |
| 190 | Cancer of eye | 2 | neoplasms | 3 | 3 | 3 |
| 191 | Malignant and unknown neoplasms of brain and nervous system | 2 | neoplasms | 11 | 11 | 11 |
| 191.1 | Cancer of brain and nervous system | 2 | neoplasms | 5 | 5 | 5 |
| 191.11 | Cancer of brain | 2 | neoplasms | 4 | 4 | 4 |
| 193 | Thyroid cancer | 2 | neoplasms | 20 | 20 | 20 |
| 194 | Cancer of other endocrine glands | 2 | neoplasms | 0 | 0 | 0 |
| 195 | Cancer, suspected or other | 2 | neoplasms | 64 | 64 | 64 |
| 195.1 | Malignant neoplasm, other | 2 | neoplasms | 59 | 59 | 59 |
| 196 | Radiotherapy | 2 | neoplasms | 0 | 0 | 0 |
| 197 | Chemotherapy | 2 | neoplasms | 29 | 29 | 29 |
| 198 | Secondary malignant neoplasm | 2 | neoplasms | 26 | 26 | 26 |
| 198.1 | Secondary malignancy of lymph nodes | 2 | neoplasms | 5 | 5 | 5 |
| 198.2 | Secondary malignancy of respiratory organs | 2 | neoplasms | 5 | 5 | 5 |
| 198.3 | Secondary malignant neoplasm of digestive systems | 2 | neoplasms | 0 | 0 | 0 |
| 198.4 | Secondary malignant neoplasm of liver | 2 | neoplasms | 5 | 5 | 5 |
| 198.5 | Secondary malignancy of brain/spine | 2 | neoplasms | 2 | 2 | 2 |
| 198.6 | Secondary malignancy of bone | 2 | neoplasms | 10 | 10 | 10 |
| 198.7 | Secondary malignant neoplasm of skin | 2 | neoplasms | 0 | 0 | 0 |
| 199 | Neoplasm of uncertain behavior | 2 | neoplasms | 94 | 94 | 94 |
| 199.4 | Neurofibromatosis | 2 | neoplasms | 2 | 2 | 2 |
| 200 | Myeloproliferative disease | 2 | neoplasms | 21 | 21 | 21 |
| 200.1 | Polycythemia vera | 2 | neoplasms | 8 | 8 | 8 |
| 201 | Hodgkin's disease | 2 | neoplasms | 6 | 6 | 6 |
| 202 | Cancer of other lymphoid, histiocytic tissue | 2 | neoplasms | 29 | 29 | 29 |
| 202.2 | Non-Hodgkins lymphoma | 2 | neoplasms | 26 | 26 | 26 |
| 202.21 | Nodular lymphoma | 2 | neoplasms | 5 | 5 | 5 |
| 202.22 | Reticulosarcoma | 2 | neoplasms | 0 | 0 | 0 |
| 202.23 | Lymphosarcoma | 2 | neoplasms | 0 | 0 | 0 |
| 202.24 | Large cell lymphoma | 2 | neoplasms | 1 | 1 | 1 |
| 204 | Leukemia | 2 | neoplasms | 14 | 14 | 14 |
| 204.1 | Lymphoid leukemia | 2 | neoplasms | 7 | 7 | 7 |
| 204.11 | Lymphoid leukemia, acute | 2 | neoplasms | 1 | 1 | 1 |
| 204.12 | Lymphoid leukemia, chronic | 2 | neoplasms | 6 | 6 | 6 |
| 204.2 | Myeloid leukemia | 2 | neoplasms | 0 | 0 | 0 |
| 204.21 | Myeloid leukemia, acute | 2 | neoplasms | 0 | 0 | 0 |
| 204.22 | Myeloid leukemia, chronic | 2 | neoplasms | 0 | 0 | 0 |
| 204.3 | Monocytic leukemia | 2 | neoplasms | 0 | 0 | 0 |
| 204.4 | Multiple myeloma | 2 | neoplasms | 5 | 5 | 5 |
| 208 | Benign neoplasm of colon | 2 | neoplasms | 149 | 149 | 149 |
| 209 | Neuroendocrine tumors | 2 | neoplasms | 4 | 4 | 4 |
| 210 | Benign neoplasm of lip, oral cavity, and pharynx | 2 | neoplasms | 1 | 1 | 1 |
| 211 | Benign neoplasm of other parts of digestive system | 2 | neoplasms | 4 | 4 | 4 |
| 212 | Benign neoplasm of respiratory and intrathoracic organs | 2 | neoplasms | 1 | 1 | 1 |
| 213 | Benign neoplasm of bone and articular cartilage | 2 | neoplasms | 1 | 1 | 1 |
| 214 | Lipoma | 2 | neoplasms | 47 | 47 | 47 |
| 214.1 | Lipoma of skin and subcutaneous tissue | 2 | neoplasms | 17 | 17 | 17 |
| 215 | Other benign neoplasm of connective and other soft tissue | 2 | neoplasms | 10 | 10 | 10 |
| 216 | Benign neoplasm of skin | 2 | neoplasms | 65 | 65 | 65 |
| 216.1 | Screening for malignant neoplasms of the skin | 2 | neoplasms | 39 | 39 | 39 |
| 217 | Vascular hamartomas and non-neoplastic nevi | 2 | neoplasms | 11 | 11 | 11 |
| 217.1 | Nevus, non-neoplastic | 2 | neoplasms | 11 | 11 | 11 |
| 218 | Benign neoplasm of uterus | 2 | neoplasms | 64 | 64 | 64 |
| 218.1 | Uterine leiomyoma | 2 | neoplasms | 62 | 62 | 62 |
| 218.2 | Other benign neoplasm of uterus | 2 | neoplasms | 0 | 0 | 0 |
| 220 | Benign neoplasm of ovary | 2 | neoplasms | 4 | 4 | 4 |
| 221 | Benign neoplasm of other female genital organs | 2 | neoplasms | 0 | 0 | 0 |
| 222 | Benign neoplasm of male genital organs | 2 | neoplasms | 0 | 0 | 0 |
| 223 | Benign neoplasm of kidney and other urinary organs | 2 | neoplasms | 3 | 3 | 3 |
| 224 | Benign neoplasm of eye | 2 | neoplasms | 3 | 3 | 3 |
| 224.1 | Benign neoplasm of eye, uveal | 2 | neoplasms | 1 | 1 | 1 |
| 225 | Benign neoplasm of brain and other parts of nervous system | 2 | neoplasms | 22 | 22 | 22 |
| 225.1 | Benign neoplasm of brain, cranial nerves, meninges | 2 | neoplasms | 18 | 18 | 18 |
| 225.2 | Benign neoplasm of spinal cord, meninges | 2 | neoplasms | 0 | 0 | 0 |
| 226 | Benign neoplasm of thyroid glands | 2 | neoplasms | 1 | 1 | 1 |
| 227 | Benign neoplasm of other endocrine glands and related structures | 2 | neoplasms | 31 | 31 | 31 |

| | | | | | | |
|---|---|---|---|---|---|---|
| 227.1 | Benign neoplasm of adrenal gland | 2 | neoplasms | 2 | 2 | 2 |
| 227.2 | Benign neoplasm of parathyroid gland | 2 | neoplasms | 2 | 2 | 2 |
| 227.3 | Benign neoplasm of pituitary gland and craniopharyngeal duct (pouch) | 2 | neoplasms | 20 | 20 | 20 |
| 228 | Hemangioma and lymphangioma, any site | 2 | neoplasms | 18 | 18 | 18 |
| 228.1 | Hemangioma of skin and subcutaneous tissue | 2 | neoplasms | 1 | 1 | 1 |
| 229 | Benign neoplasm of unspecified sites | 2 | neoplasms | 6 | 6 | 6 |
| 229.1 | Benign neoplasm of lymph nodes | 2 | neoplasms | 0 | 0 | 0 |
| 240 | Simple and unspecified goiter | 3 | endocrine/metabolic | 42 | 42 | 42 |
| 241 | Nontoxic nodular goiter | 3 | endocrine/metabolic | 160 | 160 | 160 |
| 241.1 | Nontoxic uninodular goiter | 3 | endocrine/metabolic | 0 | 0 | 0 |
| 241.2 | Nontoxic multinodular goiter | 3 | endocrine/metabolic | 79 | 79 | 79 |
| 242 | Thyrotoxicosis with or without goiter | 3 | endocrine/metabolic | 59 | 59 | 59 |
| 242.1 | Graves' disease | 3 | endocrine/metabolic | 5 | 5 | 5 |
| 242.2 | Toxic multinodular goiter | 3 | endocrine/metabolic | 1 | 1 | 1 |
| 242.3 | Exophthalmos | 3 | endocrine/metabolic | 7 | 7 | 7 |
| 242.31 | Thyrotoxic exophthalmos | 3 | endocrine/metabolic | 4 | 4 | 4 |
| 244 | Hypothyroidism | 3 | endocrine/metabolic | 803 | 803 | 803 |
| 244.1 | Secondary hypothyroidism | 3 | endocrine/metabolic | 8 | 8 | 8 |
| 244.2 | Acquired hypothyroidism | 3 | endocrine/metabolic | 249 | 249 | 249 |
| 244.3 | Iodine hypothyroidism | 3 | endocrine/metabolic | 0 | 0 | 0 |
| 244.4 | Hypothyroidism NOS | 3 | endocrine/metabolic | 732 | 732 | 732 |
| 244.5 | Congenital hypothyroidism | 3 | endocrine/metabolic | 11 | 11 | 11 |
| 245 | Thyroiditis | 3 | endocrine/metabolic | 100 | 100 | 100 |
| 245.1 | Thyroiditis, acute and subacute | 3 | endocrine/metabolic | 1 | 1 | 1 |
| 245.2 | Chronic thyroiditis | 3 | endocrine/metabolic | 89 | 89 | 89 |
| 245.21 | Chronic lymphocytic thyroiditis | 3 | endocrine/metabolic | 88 | 88 | 88 |
| 246 | Other disorders of thyroid | 3 | endocrine/metabolic | 241 | 241 | 241 |
| 246.2 | Thyroid cyst | 3 | endocrine/metabolic | 4 | 4 | 4 |
| 246.7 | Abnormal results of function study of thyroid | 3 | endocrine/metabolic | 90 | 90 | 90 |
| 249 | Secondary diabetes mellitus | 3 | endocrine/metabolic | 13 | 13 | 13 |
| 250 | Diabetes mellitus | 3 | endocrine/metabolic | 411 | 411 | 411 |
| 250.1 | Type 1 diabetes | 3 | endocrine/metabolic | 36 | 36 | 36 |
| 250.11 | Type 1 diabetes with ketoacidosis | 3 | endocrine/metabolic | 3 | 3 | 3 |
| 250.12 | Type 1 diabetes with renal manifestations | 3 | endocrine/metabolic | 6 | 6 | 6 |
| 250.13 | Type 1 diabetes with ophthalmic manifestations | 3 | endocrine/metabolic | 3 | 3 | 3 |
| 250.14 | Type 1 diabetes with neurological manifestations | 3 | endocrine/metabolic | 5 | 5 | 5 |
| 250.15 | Diabetes type 1 with peripheral circulatory disorders | 3 | endocrine/metabolic | 0 | 0 | 0 |
| 250.2 | Type 2 diabetes | 3 | endocrine/metabolic | 218 | 218 | 218 |
| 250.21 | Type 2 diabetes with ketoacidosis | 3 | endocrine/metabolic | 2 | 2 | 2 |
| 250.22 | Type 2 diabetes with renal manifestations | 3 | endocrine/metabolic | 31 | 31 | 31 |
| 250.23 | Type 2 diabetes with ophthalmic manifestations | 3 | endocrine/metabolic | 12 | 12 | 12 |
| 250.24 | Type 2 diabetes with neurological manifestations | 3 | endocrine/metabolic | 21 | 21 | 21 |
| 250.25 | Diabetes type 2 with peripheral circulatory disorders | 3 | endocrine/metabolic | 7 | 7 | 7 |
| 250.3 | Insulin pump user | 3 | endocrine/metabolic | 73 | 73 | 73 |
| 250.4 | Abnormal glucose | 3 | endocrine/metabolic | 661 | 661 | 661 |
| 250.41 | Impaired fasting glucose | 3 | endocrine/metabolic | 402 | 402 | 402 |
| 250.42 | Other abnormal glucose | 3 | endocrine/metabolic | 337 | 337 | 337 |
| 250.5 | Glycosuria or Acetonuria | 3 | endocrine/metabolic | 6 | 6 | 6 |
| 250.6 | Polyneuropathy in diabetes | 3 | endocrine/metabolic | 26 | 26 | 26 |
| 250.7 | Diabetic retinopathy | 3 | endocrine/metabolic | 11 | 11 | 11 |
| 251 | Other disorders of pancreatic internal secretion | 3 | endocrine/metabolic | 1 | 1 | 1 |
| 251.1 | Hypoglycemia | 3 | endocrine/metabolic | 21 | 21 | 21 |
| 251.8 | Abnormality of secretion of glucagon or gastrin | 3 | endocrine/metabolic | 0 | 0 | 0 |
| 252 | Disorders of parathyroid gland | 3 | endocrine/metabolic | 34 | 34 | 34 |
| 252.1 | Hyperparathyroidism | 3 | endocrine/metabolic | 19 | 19 | 19 |
| 252.2 | Hypoparathyroidism | 3 | endocrine/metabolic | 2 | 2 | 2 |
| 253 | Disorders of the pituitary gland and its hypothalamic control | 3 | endocrine/metabolic | 34 | 34 | 34 |
| 253.1 | Pituitary hyperfunction | 3 | endocrine/metabolic | 13 | 13 | 13 |
| 253.11 | Acromegaly and gigantism | 3 | endocrine/metabolic | 0 | 0 | 0 |
| 253.2 | Pituitary hypofunction | 3 | endocrine/metabolic | 4 | 4 | 4 |
| 253.3 | Diabetes insipidus | 3 | endocrine/metabolic | 1 | 1 | 1 |
| 253.4 | Anterior pituitary disorders | 3 | endocrine/metabolic | 3 | 3 | 3 |
| 253.5 | Pituitary dwarfism | 3 | endocrine/metabolic | 2 | 2 | 2 |
| 253.7 | Other disorders of neurohypophysis | 3 | endocrine/metabolic | 3 | 3 | 3 |
| 254 | Diseases of thymus gland | 3 | endocrine/metabolic | 0 | 0 | 0 |
| 255 | Disorders of adrenal glands | 3 | endocrine/metabolic | 44 | 44 | 44 |
| 255.1 | Adrenal hyperfunction | 3 | endocrine/metabolic | 5 | 5 | 5 |

| | | | | | | |
|---|---|---|---|---|---|---|
| 255.11 | Cushing's syndrome | 3 | endocrine/metabolic | 0 | 0 | 0 |
| 255.12 | Hyperaldosteronism | 3 | endocrine/metabolic | 1 | 1 | 1 |
| 255.13 | Medulloadrenal hyperfunction | 3 | endocrine/metabolic | 0 | 0 | 0 |
| 255.2 | Adrenal hypofunction | 3 | endocrine/metabolic | 21 | 21 | 21 |
| 255.21 | Glucocorticoid deficiency | 3 | endocrine/metabolic | 21 | 21 | 21 |
| 255.22 | Mineralocorticoid deficiency | 3 | endocrine/metabolic | 0 | 0 | 0 |
| 255.3 | Adrenogenital disorders | 3 | endocrine/metabolic | 0 | 0 | 0 |
| 256 | Ovarian dysfunction | 3 | endocrine/metabolic | 80 | 80 | 80 |
| 256.1 | Hyperestrogenism | 3 | endocrine/metabolic | 0 | 0 | 0 |
| 256.4 | Polycystic ovaries | 3 | endocrine/metabolic | 70 | 70 | 70 |
| 257 | Testicular dysfunction | 3 | endocrine/metabolic | 86 | 86 | 86 |
| 257.1 | Testicular hypofunction | 3 | endocrine/metabolic | 81 | 81 | 81 |
| 258 | Iatrogenic endocrine disorders | 3 | endocrine/metabolic | 1 | 1 | 1 |
| 258.1 | Postablative ovarian failure | 3 | endocrine/metabolic | 5 | 5 | 5 |
| 259 | Other endocrine disorders | 3 | endocrine/metabolic | 48 | 48 | 48 |
| 259.1 | Nonspecific abnormal results of other endocrine function study | 3 | endocrine/metabolic | 1 | 1 | 1 |
| 259.2 | Carcinoid syndrome | 3 | endocrine/metabolic | 0 | 0 | 0 |
| 259.3 | Delay in sexual development and puberty NEC | 3 | endocrine/metabolic | 0 | 0 | 0 |
| 259.4 | Precocious sexual development and puberty NEC | 3 | endocrine/metabolic | 1 | 1 | 1 |
| 259.8 | Polyglandular activity in multiple endocrine adenomatosis | 3 | endocrine/metabolic | 1 | 1 | 1 |
| 260 | Protein-calorie malnutrition | 3 | endocrine/metabolic | 27 | 27 | 27 |
| 260.1 | Cachexia | 3 | endocrine/metabolic | 0 | 0 | 0 |
| 260.2 | severe protein-calorie malnutrition | 3 | endocrine/metabolic | 10 | 10 | 10 |
| 260.21 | Kwashiorkor | 3 | endocrine/metabolic | 0 | 0 | 0 |
| 260.22 | Nutritional marasmus | 3 | endocrine/metabolic | 0 | 0 | 0 |
| 260.3 | Adult failure to thrive | 3 | endocrine/metabolic | 4 | 4 | 4 |
| 260.6 | Anorexia | 3 | endocrine/metabolic | 0 | 0 | 0 |
| 260.7 | Polyphagia | 3 | endocrine/metabolic | 7 | 7 | 7 |
| 261 | Vitamin deficiency | 3 | endocrine/metabolic | 1192 | 1192 | 1192 |
| 261.1 | Vitamin A deficiency | 3 | endocrine/metabolic | 0 | 0 | 0 |
| 261.2 | Vitamin B-complex deficiencies | 3 | endocrine/metabolic | 112 | 112 | 112 |
| 261.3 | Vitamin C deficiencies | 3 | endocrine/metabolic | 0 | 0 | 0 |
| 261.4 | Vitamin D deficiency | 3 | endocrine/metabolic | 1135 | 1135 | 1135 |
| 261.41 | Rickets or osteomalacia | 3 | endocrine/metabolic | 0 | 0 | 0 |
| 262 | Mineral deficiency NEC | 3 | endocrine/metabolic | 2 | 2 | 2 |
| 263 | Other nutritional deficiency | 3 | endocrine/metabolic | 1 | 1 | 1 |
| 264 | Lack of normal physiological development | 3 | endocrine/metabolic | 2 | 2 | 2 |
| 264.1 | Short stature | 3 | endocrine/metabolic | 1 | 1 | 1 |
| 264.2 | Failure to thrive (childhood) | 3 | endocrine/metabolic | 1 | 1 | 1 |
| 264.3 | Delayed milestones | 3 | endocrine/metabolic | 0 | 0 | 0 |
| 264.9 | Lack of normal physiological development, unspecified | 3 | endocrine/metabolic | 0 | 0 | 0 |
| 269 | Proteinuria | 3 | endocrine/metabolic | 12 | 12 | 12 |
| 270 | Disorders of protein plasma/amino-acid transport and metabolism | 3 | endocrine/metabolic | 40 | 40 | 40 |
| 270.1 | Disturbances of amino-acid transport | 3 | endocrine/metabolic | 15 | 15 | 15 |
| 270.11 | Disturbances of sulphur-bearing amino-acid metabolism | 3 | endocrine/metabolic | 15 | 15 | 15 |
| 270.12 | Phenylketonuria [PKU] | 3 | endocrine/metabolic | 0 | 0 | 0 |
| 270.2 | Disorders of amino-acid metabolism | 3 | endocrine/metabolic | 2 | 2 | 2 |
| 270.21 | Disorders of urea cycle metabolism | 3 | endocrine/metabolic | 1 | 1 | 1 |
| 270.3 | Disorders of plasma protein metabolism | 3 | endocrine/metabolic | 21 | 21 | 21 |
| 270.31 | Polyclonal hypergammaglobulinemia | 3 | endocrine/metabolic | 0 | 0 | 0 |
| 270.32 | Paraproteinemia | 3 | endocrine/metabolic | 5 | 5 | 5 |
| 270.33 | Amyloidosis | 3 | endocrine/metabolic | 1 | 1 | 1 |
| 270.34 | Alpha-1-antitrypsin deficiency | 3 | endocrine/metabolic | 5 | 5 | 5 |
| 270.35 | Macroglobulinemia | 3 | endocrine/metabolic | 0 | 0 | 0 |
| 270.38 | Other specified disorders of plasma protein metabolism | 3 | endocrine/metabolic | 11 | 11 | 11 |
| 271 | Disorders of carbohydrate transport and metabolism | 3 | endocrine/metabolic | 8 | 8 | 8 |
| 271.3 | Intestinal disaccharidase deficiencies and disaccharide malabsorption | 3 | endocrine/metabolic | 6 | 6 | 6 |
| 271.9 | Other disorders of carbohydrate transport and metabolism | 3 | endocrine/metabolic | 2 | 2 | 2 |
| 272 | Disorders of lipoid metabolism | 3 | endocrine/metabolic | 1922 | 1922 | 1922 |
| 272.1 | Hyperlipidemia | 3 | endocrine/metabolic | 1793 | 1793 | 1793 |
| 272.11 | Hypercholesterolemia | 3 | endocrine/metabolic | 142 | 142 | 142 |
| 272.12 | Hyperglyceridemia | 3 | endocrine/metabolic | 117 | 117 | 117 |
| 272.13 | Mixed hyperlipidemia | 3 | endocrine/metabolic | 345 | 345 | 345 |
| 272.14 | Hyperchylomicronemia | 3 | endocrine/metabolic | 0 | 0 | 0 |
| 272.9 | Unspecified disorder of lipoid metabolism | 3 | endocrine/metabolic | 1 | 1 | 1 |
| 274 | Gout and other crystal arthropathies | 3 | endocrine/metabolic | 121 | 121 | 121 |
| 274.1 | Gout | 3 | endocrine/metabolic | 116 | 116 | 116 |

| | | | | | | |
|---|---|---|---|---|---|---|
| 274.11 | Gouty arthropathy | 3 | endocrine/metabolic | 41 | 41 | 41 |
| 274.2 | Crystal arthropathies | 3 | endocrine/metabolic | 6 | 6 | 6 |
| 274.21 | Chondrocalcinosis | 3 | endocrine/metabolic | 5 | 5 | 5 |
| 275 | Disorders of mineral metabolism | 3 | endocrine/metabolic | 144 | 144 | 144 |
| 275.1 | Disorders of iron metabolism | 4 | hematopoietic | 35 | 35 | 35 |
| 275.11 | Hereditary hemochromatosis | 4 | hematopoietic | 7 | 7 | 7 |
| 275.2 | Disorders of copper metabolism | 3 | endocrine/metabolic | 1 | 1 | 1 |
| 275.3 | Disorders of magnesium metabolism | 3 | endocrine/metabolic | 19 | 19 | 19 |
| 275.5 | Disorders of calcium/phosphorus metabolism | 3 | endocrine/metabolic | 46 | 46 | 46 |
| 275.51 | Hypocalcemia | 3 | endocrine/metabolic | 19 | 19 | 19 |
| 275.53 | Disorders of phosphorus metabolism | 3 | endocrine/metabolic | 18 | 18 | 18 |
| 275.6 | Hypercalcemia | 3 | endocrine/metabolic | 38 | 38 | 38 |
| 276 | Disorders of fluid, electrolyte, and acid-base balance | 3 | endocrine/metabolic | 429 | 429 | 429 |
| 276.1 | Electrolyte imbalance | 3 | endocrine/metabolic | 388 | 388 | 388 |
| 276.11 | Hyperosmolality and/or hypernatremia | 3 | endocrine/metabolic | 0 | 0 | 0 |
| 276.12 | Hyposmolality and/or hyponatremia | 3 | endocrine/metabolic | 137 | 137 | 137 |
| 276.13 | Hyperpotassemia | 3 | endocrine/metabolic | 16 | 16 | 16 |
| 276.14 | Hypopotassemia | 3 | endocrine/metabolic | 254 | 254 | 254 |
| 276.4 | Acid-base balance disorder | 3 | endocrine/metabolic | 33 | 33 | 33 |
| 276.41 | Acidosis | 3 | endocrine/metabolic | 32 | 32 | 32 |
| 276.42 | Alkalosis | 3 | endocrine/metabolic | 0 | 0 | 0 |
| 276.5 | Hypovolemia | 3 | endocrine/metabolic | 64 | 64 | 64 |
| 276.6 | Fluid overload | 3 | endocrine/metabolic | 10 | 10 | 10 |
| 276.8 | Polydipsia | 3 | endocrine/metabolic | 3 | 3 | 3 |
| 277 | Other disorders of metabolism | 3 | endocrine/metabolic | 59 | 59 | 59 |
| 277.1 | Disorders of porphyrin metabolism | 3 | endocrine/metabolic | 1 | 1 | 1 |
| 277.2 | Other disorders of purine and pyrimidine metabolism | 3 | endocrine/metabolic | 0 | 0 | 0 |
| 277.4 | Disorders of bilirubin excretion | 3 | endocrine/metabolic | 18 | 18 | 18 |
| 277.5 | Other disorders of lipoid metabolism | 3 | endocrine/metabolic | 39 | 39 | 39 |
| 277.51 | Lipoprotein disorders | 3 | endocrine/metabolic | 34 | 34 | 34 |
| 277.6 | Other deficiencies of circulating enzymes | 3 | endocrine/metabolic | 0 | 0 | 0 |
| 277.7 | Dysmetabolic syndrome X | 3 | endocrine/metabolic | 30 | 30 | 30 |
| 277.8 | Carnitine deficiencies | 3 | endocrine/metabolic | 0 | 0 | 0 |
| 278 | Overweight, obesity and other hyperalimentation | 3 | endocrine/metabolic | 1325 | 1325 | 1325 |
| 278.1 | Obesity | 3 | endocrine/metabolic | 826 | 826 | 826 |
| 278.11 | Morbid obesity | 3 | endocrine/metabolic | 490 | 490 | 490 |
| 278.3 | Localized adiposity | 3 | endocrine/metabolic | 4 | 4 | 4 |
| 278.4 | Abnormal weight gain | 3 | endocrine/metabolic | 122 | 122 | 122 |
| 279 | Disorders involving the immune mechanism | 3 | endocrine/metabolic | 31 | 31 | 31 |
| 279.1 | Immunity deficiency | 3 | endocrine/metabolic | 15 | 15 | 15 |
| 279.11 | Deficiency of humoral immunity | 3 | endocrine/metabolic | 9 | 9 | 9 |
| 279.2 | Autoimmune disease NEC | 3 | endocrine/metabolic | 4 | 4 | 4 |
| 279.7 | Other immunological findings | 3 | endocrine/metabolic | 24 | 24 | 24 |
| 279.8 | Other specified disorders involving the immune mechanism | 3 | endocrine/metabolic | 2 | 2 | 2 |
| 280 | Iron deficiency anemias | 4 | hematopoietic | 124 | 124 | 124 |
| 280.1 | Iron deficiency anemias unspecified or not due to blood loss | 4 | hematopoietic | 107 | 107 | 107 |
| 280.2 | Iron deficiency anemia secondary to blood loss (chronic) | 4 | hematopoietic | 0 | 0 | 0 |
| 281 | Other deficiency anemia | 4 | hematopoietic | 28 | 28 | 28 |
| 281.1 | Megaloblastic anemia | 4 | hematopoietic | 9 | 9 | 9 |
| 281.11 | Pernicious anemia | 4 | hematopoietic | 0 | 0 | 0 |
| 281.12 | Other vitamin B12 deficiency anemia | 4 | hematopoietic | 6 | 6 | 6 |
| 281.13 | Folate-deficiency anemia | 4 | hematopoietic | 0 | 0 | 0 |
| 281.9 | Deficiency anemias | 4 | hematopoietic | 12 | 12 | 12 |
| 282 | Hereditary hemolytic anemias | 4 | hematopoietic | 24 | 24 | 24 |
| 282.5 | Sickle cell anemia | 4 | hematopoietic | 2 | 2 | 2 |
| 282.8 | Other hemoglobinopathies | 4 | hematopoietic | 21 | 21 | 21 |
| 282.9 | Other hereditary hemolytic anemias | 4 | hematopoietic | 1 | 1 | 1 |
| 283 | Acquired hemolytic anemias | 4 | hematopoietic | 2 | 2 | 2 |
| 283.1 | Autoimmune hemolytic anemias | 4 | hematopoietic | 0 | 0 | 0 |
| 283.2 | Non-autoimmune hemolytic anemias | 4 | hematopoietic | 0 | 0 | 0 |
| 283.21 | Hemolytic-uremic syndrome | 4 | hematopoietic | 0 | 0 | 0 |
| 284 | Aplastic anemia | 4 | hematopoietic | 17 | 17 | 17 |
| 284.1 | Pancytopenia | 4 | hematopoietic | 15 | 15 | 15 |
| 284.2 | Constitutional aplastic anemia | 4 | hematopoietic | 1 | 1 | 1 |
| 285 | Other anemias | 4 | hematopoietic | 311 | 311 | 311 |
| 285.1 | Acute posthemorrhagic anemia | 4 | hematopoietic | 22 | 22 | 22 |
| 285.2 | Anemia of chronic disease | 4 | hematopoietic | 9 | 9 | 9 |

| | | | | | | |
|---|---|---|---|---|---|---|
| 285.21 | Anemia in chronic kidney disease | 4 | hematopoietic | 3 | 3 | 3 |
| 285.22 | Anemia in neoplastic disease | 4 | hematopoietic | 2 | 2 | 2 |
| 285.3 | Sideroblastic anemia | 4 | hematopoietic | 0 | 0 | 0 |
| 285.8 | Hemoglobinuria | 4 | hematopoietic | 0 | 0 | 0 |
| 286 | Coagulation defects | 4 | hematopoietic | 78 | 78 | 78 |
| 286.1 | Congenital coagulation defects | 4 | hematopoietic | 6 | 6 | 6 |
| 286.11 | Von willebrand's disease | 4 | hematopoietic | 3 | 3 | 3 |
| 286.12 | Congenital deficiency of other clotting factors (including factor VII) | 4 | hematopoietic | 3 | 3 | 3 |
| 286.13 | Congenital factor VIII disorder | 4 | hematopoietic | 0 | 0 | 0 |
| 286.2 | Encounter for long-term (current) use of anticoagulants | 4 | hematopoietic | 127 | 127 | 127 |
| 286.3 | Coagulation defects complicating pregnancy or postpartum | 4 | hematopoietic | 6 | 6 | 6 |
| 286.4 | Acquired coagulation factor deficiency | 4 | hematopoietic | 2 | 2 | 2 |
| 286.5 | Hemorrhagic disorder due to intrinsic circulating anticoagulants | 4 | hematopoietic | 3 | 3 | 3 |
| 286.6 | Defibrination syndrome | 4 | hematopoietic | 0 | 0 | 0 |
| 286.7 | Other and unspecified coagulation defects | 4 | hematopoietic | 20 | 20 | 20 |
| 286.8 | Hypercoagulable state | 4 | hematopoietic | 30 | 30 | 30 |
| 286.81 | Primary hypercoagulable state | 4 | hematopoietic | 29 | 29 | 29 |
| 286.9 | Abnormal coagulation profile | 4 | hematopoietic | 15 | 15 | 15 |
| 287 | Purpura and other hemorrhagic conditions | 4 | hematopoietic | 102 | 102 | 102 |
| 287.1 | Spontaneous ecchymoses | 4 | hematopoietic | 8 | 8 | 8 |
| 287.2 | Allergic purpura | 4 | hematopoietic | 0 | 0 | 0 |
| 287.3 | Thrombocytopenia | 4 | hematopoietic | 81 | 81 | 81 |
| 287.31 | Primary thrombocytopenia | 4 | hematopoietic | 5 | 5 | 5 |
| 287.32 | Secondary thrombocytopenia | 4 | hematopoietic | 3 | 3 | 3 |
| 287.4 | Qualitative platelet defects | 4 | hematopoietic | 4 | 4 | 4 |
| 288 | Diseases of white blood cells | 4 | hematopoietic | 181 | 181 | 181 |
| 288.1 | Decreased white blood cell count | 4 | hematopoietic | 104 | 104 | 104 |
| 288.11 | Neutropenia | 4 | hematopoietic | 22 | 22 | 22 |
| 288.2 | Elevated white blood cell count | 4 | hematopoietic | 148 | 148 | 148 |
| 288.3 | Eosinophilia | 4 | hematopoietic | 3 | 3 | 3 |
| 289 | Other diseases of blood and blood-forming organs | 4 | hematopoietic | 101 | 101 | 101 |
| 289.1 | Myelofibrosis | 4 | hematopoietic | 0 | 0 | 0 |
| 289.3 | Personal history of diseases of blood and blood-forming organs | 4 | hematopoietic | 24 | 24 | 24 |
| 289.4 | Lymphadenitis | 4 | hematopoietic | 118 | 118 | 118 |
| 289.5 | Diseases of spleen | 4 | hematopoietic | 3 | 3 | 3 |
| 289.8 | Polycythemia vera, secondary | 4 | hematopoietic | 0 | 0 | 0 |
| 289.9 | Abnormality of red blood cells | 4 | hematopoietic | 16 | 16 | 16 |
| 290 | Delirium dementia and amnestic and other cognitive disorders | 5 | mental disorders | 19 | 19 | 19 |
| 290.1 | Dementias | 5 | mental disorders | 15 | 15 | 15 |
| 290.11 | Alzheimer's disease | 5 | mental disorders | 5 | 5 | 5 |
| 290.12 | Dementia with cerebral degenerations | 5 | mental disorders | 1 | 1 | 1 |
| 290.13 | Senile dementia | 5 | mental disorders | 0 | 0 | 0 |
| 290.16 | Vascular dementia | 5 | mental disorders | 3 | 3 | 3 |
| 290.2 | Delirium due to conditions classified elsewhere | 5 | mental disorders | 0 | 0 | 0 |
| 290.3 | Other persistent mental disorders due to conditions classified elsewhere | 5 | mental disorders | 6 | 6 | 6 |
| 291 | Other specified nonpsychotic and/or transient mental disorders | 5 | mental disorders | 25 | 25 | 25 |
| 291.1 | Transient mental disorders due to conditions classified elsewhere | 5 | mental disorders | 1 | 1 | 1 |
| 291.4 | Specific nonpsychotic mental disorders due to brain damage | 5 | mental disorders | 16 | 16 | 16 |
| 291.8 | Alteration of consciousness | 5 | mental disorders | 6 | 6 | 6 |
| 292 | Neurological disorders | 5 | mental disorders | 166 | 166 | 166 |
| 292.1 | Aphasia/speech disturbance | 5 | mental disorders | 30 | 30 | 30 |
| 292.11 | Aphasia | 5 | mental disorders | 6 | 6 | 6 |
| 292.12 | Symbolic dysfunction | 5 | mental disorders | 1 | 1 | 1 |
| 292.2 | Mild cognitive impairment | 5 | mental disorders | 11 | 11 | 11 |
| 292.3 | Memory loss | 5 | mental disorders | 76 | 76 | 76 |
| 292.4 | Altered mental status | 5 | mental disorders | 38 | 38 | 38 |
| 292.5 | Transient alteration of awareness | 5 | mental disorders | 2 | 2 | 2 |
| 292.6 | Hallucinations | 5 | mental disorders | 3 | 3 | 3 |
| 293 | Symptoms involving head and neck | 5 | mental disorders | 303 | 303 | 303 |
| 293.1 | Swelling, mass, or lump in head and neck [Space-occupying lesion, intracranial NOS] | 5 | mental disorders | 75 | 75 | 75 |
| 295 | Schizophrenia and other psychotic disorders | 5 | mental disorders | 15 | 15 | 15 |
| 295.1 | Schizophrenia | 5 | mental disorders | 4 | 4 | 4 |
| 295.2 | Paranoid disorders | 5 | mental disorders | 1 | 1 | 1 |
| 295.3 | Psychosis | 5 | mental disorders | 11 | 11 | 11 |
| 296 | Mood disorders | 5 | mental disorders | 643 | 643 | 643 |
| 296.1 | Bipolar | 5 | mental disorders | 57 | 57 | 57 |
| 296.2 | Depression | 5 | mental disorders | 567 | 567 | 567 |

| | | | | | | |
|---|---|---|---|---|---|---|
| 296.22 | Major depressive disorder | 5 | mental disorders | 252 | 252 | 252 |
| 297 | Suicidal ideation or attempt | 5 | mental disorders | 18 | 18 | 18 |
| 297.1 | Suicidal ideation | 5 | mental disorders | 7 | 7 | 7 |
| 297.2 | Suicide or self-inflicted injury | 5 | mental disorders | 8 | 8 | 8 |
| 300 | Anxiety, phobic and dissociative disorders | 5 | mental disorders | 951 | 951 | 951 |
| 300.1 | Anxiety disorder | 5 | mental disorders | 380 | 380 | 380 |
| 300.11 | Generalized anxiety disorder | 5 | mental disorders | 109 | 109 | 109 |
| 300.12 | Agorophobia, social phobia, and panic disorder | 5 | mental disorders | 70 | 70 | 70 |
| 300.13 | Phobia | 5 | mental disorders | 29 | 29 | 29 |
| 300.2 | Generalized anxiety & phobic disorders | 5 | mental disorders | 0 | 0 | 0 |
| 300.3 | Obsessive-compulsive disorders | 5 | mental disorders | 12 | 12 | 12 |
| 300.4 | Dysthymic disorder | 5 | mental disorders | 32 | 32 | 32 |
| 300.8 | Acute reaction to stress | 5 | mental disorders | 21 | 21 | 21 |
| 300.9 | Posttraumatic stress disorder | 5 | mental disorders | 55 | 55 | 55 |
| 301 | Personality disorders | 5 | mental disorders | 3 | 3 | 3 |
| 301.1 | Schizoid personality disorder | 5 | mental disorders | 0 | 0 | 0 |
| 301.2 | Antisocial/borderline personality disorder | 5 | mental disorders | 2 | 2 | 2 |
| 302 | Sexual and gender identity disorders | 5 | mental disorders | 31 | 31 | 31 |
| 302.1 | Decreased libido | 5 | mental disorders | 26 | 26 | 26 |
| 303 | Psychogenic and somatoform disorders | 5 | mental disorders | 4 | 4 | 4 |
| 303.1 | Dissociative disorder | 5 | mental disorders | 1 | 1 | 1 |
| 303.3 | Psychogenic disorder | 5 | mental disorders | 2 | 2 | 2 |
| 303.31 | Gastrointestinal malfunction arising from mental factors | 5 | mental disorders | 0 | 0 | 0 |
| 303.4 | Somatoform disorder | 5 | mental disorders | 2 | 2 | 2 |
| 304 | Adjustment reaction | 5 | mental disorders | 119 | 119 | 119 |
| 305.2 | Eating disorder | 5 | mental disorders | 10 | 10 | 10 |
| 305.21 | Anorexia nervosa | 5 | mental disorders | 1 | 1 | 1 |
| 306 | Other mental disorder | 5 | mental disorders | 22 | 22 | 22 |
| 306.1 | Mental disorders durring/after pregnancy | 5 | mental disorders | 10 | 10 | 10 |
| 306.9 | Tension headache | 5 | mental disorders | 3 | 3 | 3 |
| 312 | Conduct disorders | 5 | mental disorders | 4 | 4 | 4 |
| 312.3 | Impulse control disorder | 5 | mental disorders | 2 | 2 | 2 |
| 313 | Pervasive developmental disorders | 5 | mental disorders | 68 | 68 | 68 |
| 313.1 | Attention deficit hyperactivity disorder | 5 | mental disorders | 63 | 63 | 63 |
| 313.2 | Tics and stuttering | 5 | mental disorders | 2 | 2 | 2 |
| 313.3 | Autism | 5 | mental disorders | 1 | 1 | 1 |
| 315 | Develomental delays and disorders | 5 | mental disorders | 9 | 9 | 9 |
| 315.1 | Learning disorder | 5 | mental disorders | 0 | 0 | 0 |
| 315.2 | Speech and language disorder | 5 | mental disorders | 4 | 4 | 4 |
| 315.3 | Mental retardation | 5 | mental disorders | 1 | 1 | 1 |
| 316 | Substance addiction and disorders | 5 | mental disorders | 118 | 118 | 118 |
| 316.1 | Polyneuropathy due to drugs | 5 | mental disorders | 4 | 4 | 4 |
| 317 | Alcohol-related disorders | 5 | mental disorders | 43 | 43 | 43 |
| 317.1 | Alcoholism | 5 | mental disorders | 23 | 23 | 23 |
| 317.11 | Alcoholic liver damage | 5 | mental disorders | 11 | 11 | 11 |
| 318 | Tobacco use disorder | 5 | mental disorders | 197 | 197 | 197 |
| 320 | Meningitis | 6 | neurological | 4 | 4 | 4 |
| 323 | Encephalitis | 6 | neurological | 4 | 4 | 4 |
| 323.2 | Acute (transverse) myelitis | 6 | neurological | 0 | 0 | 0 |
| 323.8 | Encephalitis, non-infectious | 6 | neurological | 4 | 4 | 4 |
| 324 | Other CNS infection and poliomyelitis | 6 | neurological | 9 | 9 | 9 |
| 324.1 | Jakob-Creutzfeldt disease | 6 | neurological | 0 | 0 | 0 |
| 325 | Phlebitis and thrombophlebitis of intracranial venous sinuses | 6 | neurological | 1 | 1 | 1 |
| 327 | Sleep disorders | 6 | neurological | 626 | 626 | 626 |
| 327.1 | Hypersomnia | 6 | neurological | 17 | 17 | 17 |
| 327.3 | Sleep apnea | 6 | neurological | 467 | 467 | 467 |
| 327.31 | Central/nonobstroctive sleep apnea | 6 | neurological | 59 | 59 | 59 |
| 327.32 | Obstructive sleep apnea | 6 | neurological | 346 | 346 | 346 |
| 327.4 | Insomnia | 6 | neurological | 496 | 496 | 496 |
| 327.41 | Organic or persistent insomnia | 6 | neurological | 150 | 150 | 150 |
| 327.5 | Parasomnia | 6 | neurological | 4 | 4 | 4 |
| 327.6 | Circadian rhythm sleep disorder | 6 | neurological | 15 | 15 | 15 |
| 327.7 | Sleep related movement disorders | 6 | neurological | 70 | 70 | 70 |
| 327.71 | Restless legs syndrome | 6 | neurological | 58 | 58 | 58 |
| 327.72 | Sleep related leg cramps | 6 | neurological | 12 | 12 | 12 |
| 331 | Other cerebral degenerations | 6 | neurological | 12 | 12 | 12 |
| 331.1 | Hydrocephalus | 6 | neurological | 1 | 1 | 1 |

| | | | | | | |
|---|---|---|---|---|---|---|
| 331.9 | Cerebral degeneration, unspecified | 6 | neurological | 7 | 7 | 7 |
| 332 | Parkinson's disease | 6 | neurological | 8 | 8 | 8 |
| 333 | Extrapyramidal disease and abnormal movement disorders | 6 | neurological | 42 | 42 | 42 |
| 333.1 | Essential tremor | 6 | neurological | 36 | 36 | 36 |
| 333.2 | Myoclonus | 6 | neurological | 1 | 1 | 1 |
| 333.3 | Tics and choreas | 6 | neurological | 0 | 0 | 0 |
| 333.4 | Torsion dystonia | 6 | neurological | 2 | 2 | 2 |
| 333.8 | Other degenerative diseases of the basal ganglia | 6 | neurological | 0 | 0 | 0 |
| 334 | Degenerative disease of the spinal cord | 6 | neurological | 11 | 11 | 11 |
| 334.1 | Spinocerebellar disease | 6 | neurological | 0 | 0 | 0 |
| 334.2 | Anterior horn cell disease | 6 | neurological | 1 | 1 | 1 |
| 334.21 | Amyotrophic Lateral Sclerosis | 6 | neurological | 1 | 1 | 1 |
| 335 | Multiple sclerosis | 6 | neurological | 38 | 38 | 38 |
| 337 | Disorders of the autonomic nervous system | 6 | neurological | 7 | 7 | 7 |
| 337.1 | Peripheral autonomic neuropathy | 6 | neurological | 3 | 3 | 3 |
| 338 | Pain | 6 | neurological | 812 | 812 | 812 |
| 338.1 | Acute pain | 6 | neurological | 89 | 89 | 89 |
| 338.2 | Chronic pain | 6 | neurological | 745 | 745 | 745 |
| 339 | Other headache syndromes | 6 | neurological | 86 | 86 | 86 |
| 340 | Migraine | 6 | neurological | 457 | 457 | 457 |
| 340.1 | Migrain with aura | 6 | neurological | 46 | 46 | 46 |
| 341 | Other demyelinating diseases of central nervous system | 6 | neurological | 4 | 4 | 4 |
| 342 | Hemiplegia | 6 | neurological | 2 | 2 | 2 |
| 343 | Infantile cerebral palsy | 6 | neurological | 2 | 2 | 2 |
| 344 | Other paralytic syndromes | 6 | neurological | 7 | 7 | 7 |
| 345 | Epilepsy, recurrent seizures, convulsions | 6 | neurological | 90 | 90 | 90 |
| 345.1 | Epilepsy | 6 | neurological | 21 | 21 | 21 |
| 345.11 | Generalized convulsive epilepsy | 6 | neurological | 1 | 1 | 1 |
| 345.12 | Partial epilepsy | 6 | neurological | 16 | 16 | 16 |
| 345.3 | Convulsions | 6 | neurological | 58 | 58 | 58 |
| 346 | Abnormal findings on study of brain and/or nervous system | 6 | neurological | 4 | 4 | 4 |
| 346.1 | Nonspecific abnormal findings on radiological and other examination of skull and head | 6 | neurological | 3 | 3 | 3 |
| 346.2 | Nonspecific abnormal results of function study of brain and central nervous system | 6 | neurological | 3 | 3 | 3 |
| 346.3 | Nonspecific abnormal findings in cerebrospinal fluid | 6 | neurological | 0 | 0 | 0 |
| 347 | Cataplexy and narcolepsy | 6 | neurological | 5 | 5 | 5 |
| 348 | Other conditions of brain | 6 | neurological | 34 | 34 | 34 |
| 348.2 | Cerebral edema and compression of brain | 6 | neurological | 14 | 14 | 14 |
| 348.4 | Cerebral cysts | 6 | neurological | 0 | 0 | 0 |
| 348.7 | Coma | 6 | neurological | 1 | 1 | 1 |
| 348.8 | Encephalopathy, not elsewhere classified | 6 | neurological | 16 | 16 | 16 |
| 348.9 | Other conditions of brain, NOS | 6 | neurological | 16 | 16 | 16 |
| 349 | Other and unspecified disorders of the nervous system | 6 | neurological | 3 | 3 | 3 |
| 350 | Abnormal movement | 6 | neurological | 65 | 65 | 65 |
| 350.1 | Abnormal involuntary movements | 6 | neurological | 1 | 1 | 1 |
| 350.2 | Abnormality of gait | 6 | neurological | 39 | 39 | 39 |
| 350.3 | Lack of coordination | 6 | neurological | 16 | 16 | 16 |
| 350.5 | Abnormal reflex | 6 | neurological | 4 | 4 | 4 |
| 350.6 | Disturbances of sensation of smell and taste | 6 | neurological | 13 | 13 | 13 |
| 351 | Other peripheral nerve disorders | 6 | neurological | 198 | 198 | 198 |
| 352 | Disorders of other cranial nerves | 6 | neurological | 32 | 32 | 32 |
| 352.1 | Trigeminal nerve disorders [CN5] | 6 | neurological | 12 | 12 | 12 |
| 352.2 | Facial nerve disorders [CN7] | 6 | neurological | 19 | 19 | 19 |
| 353 | Nerve root and plexus disorders | 6 | neurological | 3 | 3 | 3 |
| 353.1 | Nerve plexus lesions | 6 | neurological | 1 | 1 | 1 |
| 353.2 | Nerve root lesions | 6 | neurological | 0 | 0 | 0 |
| 355 | Complex regional/central pain syndrome | 6 | neurological | 10 | 10 | 10 |
| 355.1 | Chronic pain syndrome | 6 | neurological | 42 | 42 | 42 |
| 356 | Hereditary and idiopathic peripheral neuropathy | 6 | neurological | 54 | 54 | 54 |
| 357 | Inflammatory and toxic neuropathy | 6 | neurological | 31 | 31 | 31 |
| 358 | Myoneural disorders | 6 | neurological | 3 | 3 | 3 |
| 358.1 | Myasthenia gravis | 6 | neurological | 0 | 0 | 0 |
| 359 | Muscular dystrophies and other myopathies | 6 | neurological | 7 | 7 | 7 |
| 359.1 | Muscular dystrophies | 6 | neurological | 1 | 1 | 1 |
| 359.2 | Myopathy | 6 | neurological | 6 | 6 | 6 |
| 360 | Disorders of the globe | 7 | sense organs | 0 | 0 | 0 |
| 360.2 | Progressive myopia | 7 | sense organs | 0 | 0 | 0 |
| 360.3 | Hypotony of eye | 7 | sense organs | 0 | 0 | 0 |

| | | | | | | |
|---|---|---|---|---|---|---|
| 361 | Retinal detachments and defects | 7 | sense organs | 13 | 13 | 13 |
| 361.1 | Retinal detachment with retinal defect | 7 | sense organs | 6 | 6 | 6 |
| 361.2 | Retinoschisis and retinal cysts | 7 | sense organs | 0 | 0 | 0 |
| 362 | Other retinal disorders | 7 | sense organs | 47 | 47 | 47 |
| 362.1 | Retinopathy of prematurity | 7 | sense organs | 0 | 0 | 0 |
| 362.2 | Degeneration of macula and posterior pole of retina | 7 | sense organs | 25 | 25 | 25 |
| 362.21 | Macular degeneration, dry | 7 | sense organs | 0 | 0 | 0 |
| 362.22 | Macular degeneration, wet | 7 | sense organs | 1 | 1 | 1 |
| 362.23 | Cystoid macular degeneration of retina | 7 | sense organs | 2 | 2 | 2 |
| 362.26 | Macular puckering of retina | 7 | sense organs | 1 | 1 | 1 |
| 362.27 | Drusen (degenerative) of retina | 7 | sense organs | 1 | 1 | 1 |
| 362.29 | Macular degeneration (senile) of retina NOS | 7 | sense organs | 0 | 0 | 0 |
| 362.3 | Other nondiabetic retinopathy | 7 | sense organs | 0 | 0 | 0 |
| 362.31 | Separation of retinal layers | 7 | sense organs | 0 | 0 | 0 |
| 362.4 | Retinal vascular changes and abnomalities | 7 | sense organs | 13 | 13 | 13 |
| 362.5 | Toxic maculopathy of retina | 7 | sense organs | 0 | 0 | 0 |
| 362.6 | Peripheral retinal degenerations | 7 | sense organs | 0 | 0 | 0 |
| 362.7 | Hereditary retinal dystrophies | 7 | sense organs | 3 | 3 | 3 |
| 362.8 | Retinal hemorrhage/ischemia | 7 | sense organs | 3 | 3 | 3 |
| 362.9 | Retinal edema | 7 | sense organs | 0 | 0 | 0 |
| 363 | Chorioretinal inflammations, scars, and other disorders of choroid | 7 | sense organs | 0 | 0 | 0 |
| 363.3 | Chorioretinal scars | 7 | sense organs | 0 | 0 | 0 |
| 363.4 | Choroidal degenerations | 7 | sense organs | 0 | 0 | 0 |
| 364 | Corneal opacity and other disorders of cornea | 7 | sense organs | 10 | 10 | 10 |
| 364.1 | Corneal opacity | 7 | sense organs | 0 | 0 | 0 |
| 364.2 | Corneal edema | 7 | sense organs | 0 | 0 | 0 |
| 364.4 | Corneal degenerations | 7 | sense organs | 3 | 3 | 3 |
| 364.41 | Keratoconus | 7 | sense organs | 2 | 2 | 2 |
| 364.5 | Corneal dystrophy | 7 | sense organs | 1 | 1 | 1 |
| 364.51 | Fuchs' dystrophy | 7 | sense organs | 0 | 0 | 0 |
| 364.9 | Cornea replaced by transplant | 7 | sense organs | 3 | 3 | 3 |
| 365 | Glaucoma | 7 | sense organs | 62 | 62 | 62 |
| 365.1 | Open-angle glaucoma | 7 | sense organs | 8 | 8 | 8 |
| 365.11 | Primary open angle glaucoma | 7 | sense organs | 3 | 3 | 3 |
| 365.2 | Primary angle-closure glaucoma | 7 | sense organs | 4 | 4 | 4 |
| 365.5 | Pseudoexfoliation glaucoma | 7 | sense organs | 0 | 0 | 0 |
| 366 | Cataract | 7 | sense organs | 154 | 154 | 154 |
| 366.1 | Nonsenile Cataract | 7 | sense organs | 0 | 0 | 0 |
| 366.2 | Senile cataract | 7 | sense organs | 58 | 58 | 58 |
| 366.3 | Traumatic cataract | 7 | sense organs | 0 | 0 | 0 |
| 367 | Disorders of refraction and accommodation; blindness and low vision | 7 | sense organs | 26 | 26 | 26 |
| 367.1 | Myopia | 7 | sense organs | 4 | 4 | 4 |
| 367.2 | Astigmatism | 7 | sense organs | 5 | 5 | 5 |
| 367.8 | Hypermetropia | 7 | sense organs | 0 | 0 | 0 |
| 367.9 | Blindness and low vision | 7 | sense organs | 13 | 13 | 13 |
| 368 | Visual disturbances | 7 | sense organs | 98 | 98 | 98 |
| 368.1 | Amblyopia | 7 | sense organs | 0 | 0 | 0 |
| 368.2 | Diplopia and disorders of binocular vision | 7 | sense organs | 10 | 10 | 10 |
| 368.3 | Anisometropia | 7 | sense organs | 0 | 0 | 0 |
| 368.4 | Visual field defects | 7 | sense organs | 18 | 18 | 18 |
| 368.5 | Color vision deficiencies | 7 | sense organs | 0 | 0 | 0 |
| 368.7 | Disorders of accommodation | 7 | sense organs | 0 | 0 | 0 |
| 368.9 | Subjective visual disturbances | 7 | sense organs | 13 | 13 | 13 |
| 368.91 | Psychophysical visual disturbances | 7 | sense organs | 1 | 1 | 1 |
| 369 | Infection of the eye | 7 | sense organs | 167 | 167 | 167 |
| 369.2 | Eye infection, viral | 7 | sense organs | 1 | 1 | 1 |
| 369.5 | Conjunctivitis, infectious | 7 | sense organs | 163 | 163 | 163 |
| 370 | Keratitis | 7 | sense organs | 3 | 3 | 3 |
| 370.1 | Corneal ulcer | 7 | sense organs | 1 | 1 | 1 |
| 370.2 | Superficial keratitis | 7 | sense organs | 2 | 2 | 2 |
| 370.3 | Keratoconjunctivitis | 7 | sense organs | 0 | 0 | 0 |
| 370.31 | Keratoconjunctivitis sicca | 7 | sense organs | 0 | 0 | 0 |
| 371 | Inflammation of the eye | 7 | sense organs | 96 | 96 | 96 |
| 371.1 | Uveitis, noninfectious or NOS | 7 | sense organs | 7 | 7 | 7 |
| 371.2 | Conjunctivitis, noninfectious | 7 | sense organs | 12 | 12 | 12 |
| 371.21 | Allergic conjunctivitis | 7 | sense organs | 11 | 11 | 11 |
| 371.3 | Inflammation of eyelids | 7 | sense organs | 53 | 53 | 53 |

| | | | | | | |
|---|---|---|---|---|---|---|
| 371.33 | Noninfectious dermatoses of eyelid | 7 | sense organs | 3 | 3 | 3 |
| 371.9 | Chronic inflammatory disorders of orbit | 7 | sense organs | 0 | 0 | 0 |
| 372 | Disorders of conjunctiva | 7 | sense organs | 22 | 22 | 22 |
| 374 | Other disorders of eyelids | 7 | sense organs | 48 | 48 | 48 |
| 374.1 | Ectropion or entropion | 7 | sense organs | 3 | 3 | 3 |
| 374.2 | Lagophthalmos | 7 | sense organs | 0 | 0 | 0 |
| 374.3 | Ptosis of eyelid | 7 | sense organs | 19 | 19 | 19 |
| 374.6 | Dermatochalasis | 7 | sense organs | 10 | 10 | 10 |
| 375 | Disorders of lacrimal system | 7 | sense organs | 4 | 4 | 4 |
| 375.1 | Dry eyes | 7 | sense organs | 25 | 25 | 25 |
| 375.2 | Epiphora | 7 | sense organs | 1 | 1 | 1 |
| 376 | Disorders of the orbit | 7 | sense organs | 2 | 2 | 2 |
| 377 | Disorders of optic nerve and visual pathways | 7 | sense organs | 28 | 28 | 28 |
| 377.1 | Optic atrophy | 7 | sense organs | 2 | 2 | 2 |
| 377.3 | Optic neuritis/neuropathy | 7 | sense organs | 20 | 20 | 20 |
| 378 | Strabismus and other disorders of binocular eye movements | 7 | sense organs | 14 | 14 | 14 |
| 378.1 | Strabismus (not specified as paralytic) | 7 | sense organs | 3 | 3 | 3 |
| 378.2 | Nystagmus and other irregular eye movements | 7 | sense organs | 1 | 1 | 1 |
| 378.5 | Paralytic strabismus | 7 | sense organs | 4 | 4 | 4 |
| 379 | Other disorders of eye | 7 | sense organs | 35 | 35 | 35 |
| 379.1 | Scleritis and episcleritis | 7 | sense organs | 0 | 0 | 0 |
| 379.2 | Disorders of vitreous body | 7 | sense organs | 17 | 17 | 17 |
| 379.3 | Aphakia and other disorders of lens | 7 | sense organs | 1 | 1 | 1 |
| 379.4 | Anomalies of pupillary function | 7 | sense organs | 5 | 5 | 5 |
| 379.5 | Disorders of iris and ciliary body | 7 | sense organs | 1 | 1 | 1 |
| 379.51 | Pigmentary iris degeneration | 7 | sense organs | 0 | 0 | 0 |
| 379.9 | Pain, swelling or discharge of eye | 7 | sense organs | 29 | 29 | 29 |
| 380 | Disorders of external ear | 7 | sense organs | 3 | 3 | 3 |
| 380.1 | Otitis externa | 7 | sense organs | 60 | 60 | 60 |
| 380.4 | Impacted cerumen | 7 | sense organs | 135 | 135 | 135 |
| 381 | Otitis media and Eustachian tube disorders | 7 | sense organs | 345 | 345 | 345 |
| 381.1 | Otitis media | 7 | sense organs | 238 | 238 | 238 |
| 381.11 | Suppurative and unspecified otitis media | 7 | sense organs | 168 | 168 | 168 |
| 381.2 | Eustachian tube disorders | 7 | sense organs | 75 | 75 | 75 |
| 381.3 | Mastoiditis & related conditions | 7 | sense organs | 9 | 9 | 9 |
| 381.9 | Otorrhea | 7 | sense organs | 3 | 3 | 3 |
| 382 | Otalgia | 7 | sense organs | 173 | 173 | 173 |
| 383 | Otosclerosis | 7 | sense organs | 1 | 1 | 1 |
| 384 | Other disorders of tympanic membrane | 7 | sense organs | 12 | 12 | 12 |
| 384.1 | Myringitis | 7 | sense organs | 4 | 4 | 4 |
| 384.4 | Perforation of tympanic membrane | 7 | sense organs | 7 | 7 | 7 |
| 385 | Other disorders of middle ear and mastoid | 7 | sense organs | 6 | 6 | 6 |
| 385.3 | Cholesteatoma | 7 | sense organs | 4 | 4 | 4 |
| 385.5 | Tympanosclerosis and middle ear disease related to otitis media | 7 | sense organs | 1 | 1 | 1 |
| 386 | Vertiginous syndromes and other disorders of vestibular system | 7 | sense organs | 76 | 76 | 76 |
| 386.1 | Meniere's disease | 7 | sense organs | 0 | 0 | 0 |
| 386.2 | Peripheral or central vertigo | 7 | sense organs | 60 | 60 | 60 |
| 386.21 | Central origin vertigo | 7 | sense organs | 0 | 0 | 0 |
| 386.3 | Labyrinthitis | 7 | sense organs | 2 | 2 | 2 |
| 386.9 | Dizziness and giddiness (Light-headedness and vertigo) | 7 | sense organs | 401 | 401 | 401 |
| 388 | Other disorders of ear | 7 | sense organs | 35 | 35 | 35 |
| 389 | Hearing loss | 7 | sense organs | 168 | 168 | 168 |
| 389.1 | Sensorineural hearing loss | 7 | sense organs | 1 | 1 | 1 |
| 389.2 | Conductive hearing loss | 7 | sense organs | 1 | 1 | 1 |
| 389.3 | Degenerative and vascular disorders of ear | 7 | sense organs | 5 | 5 | 5 |
| 389.4 | Tinnitus | 7 | sense organs | 36 | 36 | 36 |
| 389.5 | Disorders of acoustic nerve | 7 | sense organs | 1 | 1 | 1 |
| 394 | Rheumatic disease of the heart valves | 8 | circulatory system | 66 | 66 | 66 |
| 394.1 | Mitral valve stenosis and aortic valve stenosis | 8 | circulatory system | 5 | 5 | 5 |
| 394.2 | Mitral valve disease | 8 | circulatory system | 12 | 12 | 12 |
| 394.3 | Aortic valve disease | 8 | circulatory system | 1 | 1 | 1 |
| 394.4 | Acute rheumatic heart disease | 8 | circulatory system | 1 | 1 | 1 |
| 394.7 | Disease of tricuspid valve | 8 | circulatory system | 0 | 0 | 0 |
| 395 | Heart valve disorders | 8 | circulatory system | 331 | 331 | 331 |
| 395.1 | Nonrheumatic mitral valve disorders | 8 | circulatory system | 47 | 47 | 47 |
| 395.2 | Nonrheumatic aortic valve disorders | 8 | circulatory system | 93 | 93 | 93 |
| 395.3 | Nonrheumatic tricuspid valve disorders | 8 | circulatory system | 43 | 43 | 43 |

| | | | | | | |
|---|---|---|---|---|---|---|
| 395.4 | Nonrheumatic pulmonary valve disorders | 8 | circulatory system | 12 | 12 | 12 |
| 395.6 | Heart valve replaced | 8 | circulatory system | 22 | 22 | 22 |
| 396 | Abnormal heart sounds | 8 | circulatory system | 101 | 101 | 101 |
| 401 | Hypertension | 8 | circulatory system | 1242 | 1242 | 1242 |
| 401.1 | Essential hypertension | 8 | circulatory system | 1229 | 1229 | 1229 |
| 401.2 | Hypertensive heart and/or renal disease | 8 | circulatory system | 9 | 9 | 9 |
| 401.21 | Hypertensive heart disease | 8 | circulatory system | 2 | 2 | 2 |
| 401.22 | Hypertensive chronic kidney disease | 8 | circulatory system | 5 | 5 | 5 |
| 401.3 | Other hypertensive complications | 8 | circulatory system | 43 | 43 | 43 |
| 411 | Ischemic Heart Disease | 8 | circulatory system | 254 | 254 | 254 |
| 411.1 | Unstable angina (intermediate coronary syndrome) | 8 | circulatory system | 14 | 14 | 14 |
| 411.2 | Myocardial infarction | 8 | circulatory system | 97 | 97 | 97 |
| 411.3 | Angina pectoris | 8 | circulatory system | 36 | 36 | 36 |
| 411.4 | Coronary atherosclerosis | 8 | circulatory system | 158 | 158 | 158 |
| 411.41 | Aneurysm and dissection of heart | 8 | circulatory system | 1 | 1 | 1 |
| 411.8 | Other chronic ischemic heart disease, unspecified | 8 | circulatory system | 115 | 115 | 115 |
| 411.9 | Other acute and subacute forms of ischemic heart disease | 8 | circulatory system | 0 | 0 | 0 |
| 414 | Other forms of chronic heart disease | 8 | circulatory system | 52 | 52 | 52 |
| 414.2 | ASCVD | 8 | circulatory system | 2 | 2 | 2 |
| 415 | Pulmonary heart disease | 8 | circulatory system | 97 | 97 | 97 |
| 415.1 | Acute pulmonary heart disease | 8 | circulatory system | 37 | 37 | 37 |
| 415.11 | Pulmonary embolism and infarction, acute | 8 | circulatory system | 37 | 37 | 37 |
| 415.2 | Chronic pulmonary heart disease | 8 | circulatory system | 48 | 48 | 48 |
| 415.21 | Primary pulmonary hypertension | 8 | circulatory system | 0 | 0 | 0 |
| 416 | Cardiomegaly | 8 | circulatory system | 54 | 54 | 54 |
| 418 | Nonspecific chest pain | 8 | circulatory system | 764 | 764 | 764 |
| 418.1 | Precordial pain | 8 | circulatory system | 26 | 26 | 26 |
| 420 | Carditis | 8 | circulatory system | 33 | 33 | 33 |
| 420.1 | Myocarditis | 8 | circulatory system | 1 | 1 | 1 |
| 420.2 | Pericarditis | 8 | circulatory system | 23 | 23 | 23 |
| 420.21 | Acute pericarditis | 8 | circulatory system | 2 | 2 | 2 |
| 420.22 | Chronic pericarditis | 8 | circulatory system | 3 | 3 | 3 |
| 420.3 | Endocarditis | 8 | circulatory system | 8 | 8 | 8 |
| 425 | Cardiomyopathy | 8 | circulatory system | 27 | 27 | 27 |
| 425.1 | Primary/intrinsic cardiomyopathies | 8 | circulatory system | 26 | 26 | 26 |
| 425.11 | Hypertrophic obstructive cardiomyopathy | 8 | circulatory system | 1 | 1 | 1 |
| 425.12 | Other hypertrophic cardiomyopathy | 8 | circulatory system | 4 | 4 | 4 |
| 425.2 | Secondary/extrinsic cardiomyopathies | 8 | circulatory system | 3 | 3 | 3 |
| 425.8 | Other cardiomyopathy | 8 | circulatory system | 2 | 2 | 2 |
| 426 | Cardiac conduction disorders | 8 | circulatory system | 241 | 241 | 241 |
| 426.2 | Atrioventricular [AV] block | 8 | circulatory system | 21 | 21 | 21 |
| 426.21 | First degree AV block | 8 | circulatory system | 9 | 9 | 9 |
| 426.22 | Mobitz II AV block | 8 | circulatory system | 1 | 1 | 1 |
| 426.23 | Second degree AV block | 8 | circulatory system | 2 | 2 | 2 |
| 426.24 | Atrioventricular block, complete | 8 | circulatory system | 0 | 0 | 0 |
| 426.25 | Other heart block | 8 | circulatory system | 3 | 3 | 3 |
| 426.3 | Bundle branch block | 8 | circulatory system | 53 | 53 | 53 |
| 426.31 | Right bundle branch block | 8 | circulatory system | 24 | 24 | 24 |
| 426.32 | Left bundle branch block | 8 | circulatory system | 23 | 23 | 23 |
| 426.4 | Anomalous atrioventricular excitation | 8 | circulatory system | 5 | 5 | 5 |
| 426.7 | Abnormal electrocardiogram [ECG] [EKG] | 8 | circulatory system | 117 | 117 | 117 |
| 426.8 | Other cardiac conduction disorders | 8 | circulatory system | 10 | 10 | 10 |
| 426.9 | Cardiac pacemaker/device in situ | 8 | circulatory system | 45 | 45 | 45 |
| 426.91 | Cardiac pacemaker in situ | 8 | circulatory system | 41 | 41 | 41 |
| 426.92 | Cardiac defibrillator in situ | 8 | circulatory system | 7 | 7 | 7 |
| 427 | Cardiac dysrhythmias | 8 | circulatory system | 655 | 655 | 655 |
| 427.1 | Paroxysmal tachycardia, unspecified | 8 | circulatory system | 31 | 31 | 31 |
| 427.11 | Paroxysmal supraventricular tachycardia | 8 | circulatory system | 0 | 0 | 0 |
| 427.12 | Paroxysmal ventricular tachycardia | 8 | circulatory system | 24 | 24 | 24 |
| 427.2 | Atrial fibrillation and flutter | 8 | circulatory system | 153 | 153 | 153 |
| 427.21 | Atrial fibrillation | 8 | circulatory system | 146 | 146 | 146 |
| 427.22 | Atrial flutter | 8 | circulatory system | 24 | 24 | 24 |
| 427.3 | Other specified cardiac dysrhythmias | 8 | circulatory system | 125 | 125 | 125 |
| 427.4 | Cardiac arrest and ventricular fibrillation | 8 | circulatory system | 9 | 9 | 9 |
| 427.41 | Ventricular fibrillation and flutter | 8 | circulatory system | 3 | 3 | 3 |
| 427.42 | Cardiac arrest | 8 | circulatory system | 6 | 6 | 6 |
| 427.5 | Arrhythmia (cardiac) NOS | 8 | circulatory system | 76 | 76 | 76 |

| | | | | | | |
|---|---|---|---|---|---|---|
| 427.6 | Premature beats | 8 | circulatory system | 176 | 176 | 176 |
| 427.61 | Supraventricular premature beats | 8 | circulatory system | 76 | 76 | 76 |
| 427.7 | Tachycardia NOS | 8 | circulatory system | 0 | 0 | 0 |
| 427.8 | Sinoatrial node dysfunction (Bradycardia) | 8 | circulatory system | 27 | 27 | 27 |
| 427.9 | Palpitations | 8 | circulatory system | 279 | 279 | 279 |
| 428 | Congestive heart failure; nonhypertensive | 8 | circulatory system | 58 | 58 | 58 |
| 428.1 | Congestive heart failure (CHF) NOS | 8 | circulatory system | 11 | 11 | 11 |
| 428.2 | Heart failure NOS | 8 | circulatory system | 4 | 4 | 4 |
| 428.3 | Heart failure with reduced EF [Systolic or combined heart failure] | 8 | circulatory system | 13 | 13 | 13 |
| 428.4 | Heart failure with preserved EF [Diastolic heart failure] | 8 | circulatory system | 10 | 10 | 10 |
| 429 | Ill-defined descriptions and complications of heart disease | 8 | circulatory system | 99 | 99 | 99 |
| 429.1 | Heart transplant/surgery | 8 | circulatory system | 6 | 6 | 6 |
| 429.2 | Abnormal function study of cardiovascular system | 8 | circulatory system | 48 | 48 | 48 |
| 429.3 | Symptoms involving cardiovascular system | 8 | circulatory system | 48 | 48 | 48 |
| 429.9 | Cardiac complications, not elsewhere classified | 8 | circulatory system | 1 | 1 | 1 |
| 430 | Intracranial hemorrhage | 8 | circulatory system | 24 | 24 | 24 |
| 430.1 | Subarachnoid hemorrhage | 8 | circulatory system | 6 | 6 | 6 |
| 430.2 | Intracerebral hemorrhage | 8 | circulatory system | 8 | 8 | 8 |
| 430.3 | Subdural hemorrhage | 8 | circulatory system | 8 | 8 | 8 |
| 433 | Cerebrovascular disease | 8 | circulatory system | 218 | 218 | 218 |
| 433.1 | Occlusion and stenosis of precerebral arteries | 8 | circulatory system | 86 | 86 | 86 |
| 433.11 | Occlusion of cerebral arteries, with cerebral infarction | 8 | circulatory system | 6 | 6 | 6 |
| 433.12 | Cerebral atherosclerosis | 8 | circulatory system | 0 | 0 | 0 |
| 433.2 | Occlusion of cerebral arteries | 8 | circulatory system | 56 | 56 | 56 |
| 433.21 | Cerebral artery occlusion, with cerebral infarction | 8 | circulatory system | 55 | 55 | 55 |
| 433.3 | Cerebral ischemia | 8 | circulatory system | 97 | 97 | 97 |
| 433.31 | Transient cerebral ischemia | 8 | circulatory system | 95 | 95 | 95 |
| 433.32 | Moyamoya disease | 8 | circulatory system | 0 | 0 | 0 |
| 433.5 | Cerebral aneurysm | 8 | circulatory system | 11 | 11 | 11 |
| 433.6 | Acute, but ill-defined cerebrovascular disease | 8 | circulatory system | 3 | 3 | 3 |
| 433.8 | Late effects of cerebrovascular disease | 8 | circulatory system | 25 | 25 | 25 |
| 440 | Atherosclerosis | 8 | circulatory system | 76 | 76 | 76 |
| 440.1 | Atherosclerosis of renal artery | 8 | circulatory system | 2 | 2 | 2 |
| 440.2 | Atherosclerosis of the extremities | 8 | circulatory system | 13 | 13 | 13 |
| 440.21 | Atherosclerosis of native arteries of the extremities with ulceration or gangrene | 8 | circulatory system | 0 | 0 | 0 |
| 440.22 | Atherosclerosis of native arteries of the extremities with intermittent claudication | 8 | circulatory system | 4 | 4 | 4 |
| 440.9 | Atherosclerosis of aorta | 8 | circulatory system | 1 | 1 | 1 |
| 441 | Vascular insufficiency of intestine | 8 | circulatory system | 6 | 6 | 6 |
| 441.1 | Acute vascular insufficiency of intestine | 8 | circulatory system | 1 | 1 | 1 |
| 441.2 | Chronic vascular insufficiency of intestine | 8 | circulatory system | 0 | 0 | 0 |
| 442 | Other aneurysm | 8 | circulatory system | 47 | 47 | 47 |
| 442.1 | Aortic aneurysm | 8 | circulatory system | 41 | 41 | 41 |
| 442.11 | Abdominal aortic aneurysm | 8 | circulatory system | 22 | 22 | 22 |
| 442.2 | Aneurysm of iliac artery | 8 | circulatory system | 2 | 2 | 2 |
| 442.3 | Aneurysm of artery of lower extremity | 8 | circulatory system | 1 | 1 | 1 |
| 442.4 | Arterial dissection | 8 | circulatory system | 1 | 1 | 1 |
| 442.8 | Aneurysm of other specified artery | 8 | circulatory system | 0 | 0 | 0 |
| 443 | Peripheral vascular disease | 8 | circulatory system | 74 | 74 | 74 |
| 443.1 | Raynaud's syndrome | 8 | circulatory system | 0 | 0 | 0 |
| 443.7 | Peripheral angiopathy in diseases classified elsewhere | 8 | circulatory system | 6 | 6 | 6 |
| 443.8 | Other specified peripheral vascular diseases | 8 | circulatory system | 0 | 0 | 0 |
| 443.9 | Peripheral vascular disease, unspecified | 8 | circulatory system | 35 | 35 | 35 |
| 444 | Arterial embolism and thrombosis | 8 | circulatory system | 7 | 7 | 7 |
| 444.1 | Arterial embolism and thrombosis of lower extremity artery | 8 | circulatory system | 2 | 2 | 2 |
| 444.2 | Embolism and thrombosis of abdominal aorta | 8 | circulatory system | 0 | 0 | 0 |
| 444.5 | Atheroembolism | 8 | circulatory system | 0 | 0 | 0 |
| 446 | Polyarteritis nodosa and allied conditions | 8 | circulatory system | 10 | 10 | 10 |
| 446.1 | Thromboangiitis obliterans | 8 | circulatory system | 0 | 0 | 0 |
| 446.2 | Acute febrile mucocutaneous lymph node syndrome (Kawasaki disease) | 8 | circulatory system | 0 | 0 | 0 |
| 446.3 | Hypersensitivity angiitis | 8 | circulatory system | 0 | 0 | 0 |
| 446.4 | Wegener's granulomatosis | 8 | circulatory system | 1 | 1 | 1 |
| 446.5 | Giant cell arteritis | 8 | circulatory system | 3 | 3 | 3 |
| 446.6 | Polyarteritis nodosa | 8 | circulatory system | 0 | 0 | 0 |
| 446.7 | Takayasu's disease | 8 | circulatory system | 0 | 0 | 0 |
| 446.8 | Thrombotic microangiopathy | 8 | circulatory system | 0 | 0 | 0 |
| 446.9 | Arteritis NOS | 8 | circulatory system | 3 | 3 | 3 |
| 447 | Other disorders of arteries and arterioles | 8 | circulatory system | 54 | 54 | 54 |

| | | | | | | |
|---|---|---|---|---|---|---|
| 447.1 | Stricture of artery | 8 | circulatory system | 6 | 6 | 6 |
| 447.7 | Aortic ectasia | 8 | circulatory system | 24 | 24 | 24 |
| 448 | Disease of capillaries | 8 | circulatory system | 1 | 1 | 1 |
| 450 | Noninfectious disorders of lymphatic channels | 8 | circulatory system | 22 | 22 | 22 |
| 451 | Phlebitis and thrombophlebitis | 8 | circulatory system | 23 | 23 | 23 |
| 451.2 | Phlebitis and thrombophlebitis of lower extremities | 8 | circulatory system | 8 | 8 | 8 |
| 452 | Other venous embolism and thrombosis | 8 | circulatory system | 130 | 130 | 130 |
| 452.1 | Iatrogenic pulmonary embolism and infarction | 8 | circulatory system | 5 | 5 | 5 |
| 452.2 | Deep vein thrombosis [DVT] | 8 | circulatory system | 72 | 72 | 72 |
| 452.8 | Postphlebitic syndrome | 8 | circulatory system | 2 | 2 | 2 |
| 453 | Chronic venous hypertension | 8 | circulatory system | 3 | 3 | 3 |
| 454 | Varicose veins | 8 | circulatory system | 57 | 57 | 57 |
| 454.1 | Varicose veins of lower extremity | 8 | circulatory system | 50 | 50 | 50 |
| 454.11 | Varicose veins of lower extremity, symptomtic | 8 | circulatory system | 13 | 13 | 13 |
| 455 | Hemorrhoids | 8 | circulatory system | 89 | 89 | 89 |
| 456 | Chronic venous insufficiency [CVI] | 8 | circulatory system | 24 | 24 | 24 |
| 457 | Encounter for long-term (current) use of anticoagulants, antithrombotics, aspirin | 8 | circulatory system | 66 | 66 | 66 |
| 457.2 | Encounter for long-term (current) use of antiplatelets/antithrombotics | 8 | circulatory system | 1 | 1 | 1 |
| 457.3 | Encounter for long-term (current) use of aspirin | 8 | circulatory system | 42 | 42 | 42 |
| 458 | Hypotension | 8 | circulatory system | 70 | 70 | 70 |
| 458.1 | Orthostatic hypotension | 8 | circulatory system | 0 | 0 | 0 |
| 458.2 | Iatrogenic hypotension | 8 | circulatory system | 2 | 2 | 2 |
| 458.9 | Hypotension NOS | 8 | circulatory system | 48 | 48 | 48 |
| 459 | Other disorders of circulatory system | 8 | circulatory system | 31 | 31 | 31 |
| 459.1 | Hemorrhage NOS | 8 | circulatory system | 0 | 0 | 0 |
| 459.7 | Blood vessel replaced | 8 | circulatory system | 1 | 1 | 1 |
| 459.9 | Circulatory disease NEC | 8 | circulatory system | 24 | 24 | 24 |
| 464 | Acute sinusitis | 9 | respiratory | 696 | 696 | 696 |
| 465 | Acute upper respiratory infections of multiple or unspecified sites | 9 | respiratory | 1296 | 1296 | 1296 |
| 465.2 | Acute pharyngitis | 9 | respiratory | 747 | 747 | 747 |
| 465.4 | Acute laryngitis and tracheitis | 9 | respiratory | 30 | 30 | 30 |
| 470 | Septal Deviations/Turbinate Hypertrophy | 9 | respiratory | 45 | 45 | 45 |
| 471 | Nasal polyps | 9 | respiratory | 10 | 10 | 10 |
| 472 | Chronic pharyngitis and nasopharyngitis | 9 | respiratory | 28 | 28 | 28 |
| 473 | Diseases of the larynx and vocal cords | 9 | respiratory | 43 | 43 | 43 |
| 473.1 | Chronic laryngitis | 9 | respiratory | 1 | 1 | 1 |
| 473.3 | Paralysis/spasm of vocal cords or larynx | 9 | respiratory | 4 | 4 | 4 |
| 473.4 | Voice disturbance | 9 | respiratory | 34 | 34 | 34 |
| 474 | Acute and chronic tonsillitis | 9 | respiratory | 104 | 104 | 104 |
| 474.1 | Acute tonsillitis | 9 | respiratory | 66 | 66 | 66 |
| 474.2 | Chronic tonsillitis and adenoiditis | 9 | respiratory | 36 | 36 | 36 |
| 475 | Chronic sinusitis | 9 | respiratory | 456 | 456 | 456 |
| 475.9 | Postnasal drip | 9 | respiratory | 41 | 41 | 41 |
| 476 | Allergic rhinitis | 9 | respiratory | 695 | 695 | 695 |
| 477 | Epistaxis or throat hemorrhage | 9 | respiratory | 39 | 39 | 39 |
| 478 | Throat pain | 9 | respiratory | 11 | 11 | 11 |
| 479 | Other upper respiratory disease | 9 | respiratory | 250 | 250 | 250 |
| 480 | Pneumonia | 9 | respiratory | 191 | 191 | 191 |
| 480.1 | Bacterial pneumonia | 9 | respiratory | 11 | 11 | 11 |
| 480.11 | Pneumococcal pneumonia | 9 | respiratory | 4 | 4 | 4 |
| 480.12 | Pseudomonal pneumonia | 9 | respiratory | 0 | 0 | 0 |
| 480.13 | MRSA pneumonia | 9 | respiratory | 0 | 0 | 0 |
| 480.2 | Viral pneumonia | 9 | respiratory | 1 | 1 | 1 |
| 480.3 | Pneumonia due to fungus (mycoses) | 9 | respiratory | 2 | 2 | 2 |
| 480.5 | Bronchopneumonia and lung abscess | 9 | respiratory | 1 | 1 | 1 |
| 481 | Influenza | 9 | respiratory | 47 | 47 | 47 |
| 483 | Acute bronchitis and bronchiolitis | 9 | respiratory | 194 | 194 | 194 |
| 495 | Asthma | 9 | respiratory | 606 | 606 | 606 |
| 495.1 | Chronic obstructive asthma | 9 | respiratory | 16 | 16 | 16 |
| 495.11 | Chronic obstructive asthma with exacerbation | 9 | respiratory | 11 | 11 | 11 |
| 495.2 | Asthma with exacerbation | 9 | respiratory | 136 | 136 | 136 |
| 496 | Chronic airway obstruction | 9 | respiratory | 141 | 141 | 141 |
| 496.1 | Emphysema | 9 | respiratory | 33 | 33 | 33 |
| 496.2 | Chronic bronchitis | 9 | respiratory | 54 | 54 | 54 |
| 496.21 | Obstructive chronic bronchitis | 9 | respiratory | 32 | 32 | 32 |
| 496.3 | Bronchiectasis | 9 | respiratory | 8 | 8 | 8 |
| 497 | Bronchitis | 9 | respiratory | 468 | 468 | 468 |

| | | | | | | |
|---|---|---|---|---|---|---|
| 498 | Acute bronchospasm | 9 | respiratory | 83 | 83 | 83 |
| 499 | Cystic fibrosis | 9 | respiratory | 1 | 1 | 1 |
| 500 | Lung disease due to external agents | 9 | respiratory | 4 | 4 | 4 |
| 500.1 | Extrinsic allergic alveolitis | 9 | respiratory | 0 | 0 | 0 |
| 500.2 | Pneumoconiosis | 9 | respiratory | 0 | 0 | 0 |
| 501 | Pneumonitis due to inhalation of food or vomitus | 9 | respiratory | 0 | 0 | 0 |
| 502 | Postinflammatory pulmonary fibrosis | 9 | respiratory | 12 | 12 | 12 |
| 503 | Pulmonary congestion and hypostasis | 9 | respiratory | 4 | 4 | 4 |
| 504 | Other alveolar and parietoalveolar pneumonopathy | 9 | respiratory | 3 | 3 | 3 |
| 504.1 | Idiopathic fibrosing alveolitis | 9 | respiratory | 3 | 3 | 3 |
| 505 | Other pulmonary inflamation or edema | 9 | respiratory | 3 | 3 | 3 |
| 506 | Empyema and pneumothorax | 9 | respiratory | 10 | 10 | 10 |
| 507 | Pleurisy; pleural effusion | 9 | respiratory | 16 | 16 | 16 |
| 508 | Pulmonary collapse; interstitial and compensatory emphysema | 9 | respiratory | 2 | 2 | 2 |
| 509 | Respiratory failure, insufficiency, arrest | 9 | respiratory | 89 | 89 | 89 |
| 509.1 | Respiratory failure | 9 | respiratory | 63 | 63 | 63 |
| 509.2 | Respiratory insufficiency | 9 | respiratory | 0 | 0 | 0 |
| 509.3 | Pulmonary insufficiency or respiratory failure following trauma and surgery | 9 | respiratory | 5 | 5 | 5 |
| 509.5 | Respiratory arrest | 9 | respiratory | 0 | 0 | 0 |
| 509.8 | Dependence on respirator [Ventilator] or supplemental oxygen | 9 | respiratory | 33 | 33 | 33 |
| 510 | Other diseases of lung | 9 | respiratory | 36 | 36 | 36 |
| 510.2 | Lung transplant | 9 | respiratory | 0 | 0 | 0 |
| 512 | Other symptoms of respiratory system | 9 | respiratory | 1436 | 1436 | 1436 |
| 512.1 | Wheezing | 9 | respiratory | 95 | 95 | 95 |
| 512.2 | Painful respiration | 9 | respiratory | 94 | 94 | 94 |
| 512.3 | Abnormal chest sounds | 9 | respiratory | 10 | 10 | 10 |
| 512.7 | Shortness of breath | 9 | respiratory | 314 | 314 | 314 |
| 512.8 | Cough | 9 | respiratory | 785 | 785 | 785 |
| 512.9 | Other dyspnea | 9 | respiratory | 337 | 337 | 337 |
| 513 | Respiratory abnormalities | 9 | respiratory | 29 | 29 | 29 |
| 513.3 | Hypoventilation | 9 | respiratory | 14 | 14 | 14 |
| 513.31 | Apnea | 9 | respiratory | 5 | 5 | 5 |
| 513.32 | Orthopnea | 9 | respiratory | 2 | 2 | 2 |
| 513.4 | Hyperventilation | 9 | respiratory | 3 | 3 | 3 |
| 513.8 | Disorders of diaphragm | 9 | respiratory | 12 | 12 | 12 |
| 514 | Abnormal findings examination of lungs | 9 | respiratory | 71 | 71 | 71 |
| 514.1 | Abnormal results of function study of pulmonary system | 9 | respiratory | 0 | 0 | 0 |
| 514.2 | Solitary pulmonary nodule | 9 | respiratory | 85 | 85 | 85 |
| 516 | Abnormal sputum | 9 | respiratory | 12 | 12 | 12 |
| 516.1 | Hemoptysis | 9 | respiratory | 12 | 12 | 12 |
| 519 | Other diseases of respiratory system, not elsewhere classified | 9 | respiratory | 170 | 170 | 170 |
| 519.1 | Tracheostomy complications | 9 | respiratory | 1 | 1 | 1 |
| 519.2 | Respiratory complications | 9 | respiratory | 0 | 0 | 0 |
| 519.8 | Other diseases of respiratory system, NEC | 9 | respiratory | 147 | 147 | 147 |
| 519.9 | Symptoms involving respiratory system and other chest symptoms | 9 | respiratory | 18 | 18 | 18 |
| 520 | Disorders of tooth development | 10 | digestive | 0 | 0 | 0 |
| 520.1 | Hereditary disturbances in tooth structure | 10 | digestive | 0 | 0 | 0 |
| 520.2 | Disturbances in tooth eruption | 10 | digestive | 0 | 0 | 0 |
| 521 | Diseases of hard tissues of teeth | 10 | digestive | 12 | 12 | 12 |
| 521.1 | Dental caries | 10 | digestive | 5 | 5 | 5 |
| 521.2 | Dental abrasion, erosion and attrition | 10 | digestive | 1 | 1 | 1 |
| 521.4 | Tooth complications likely association with other diseases | 10 | digestive | 0 | 0 | 0 |
| 522 | Diseases of pulp and periapical tissues | 10 | digestive | 24 | 24 | 24 |
| 522.1 | Pulpitis and necrosis of tooth pulp | 10 | digestive | 0 | 0 | 0 |
| 522.5 | Periapical abscess | 10 | digestive | 24 | 24 | 24 |
| 523 | Gingival and periodontal diseases | 10 | digestive | 14 | 14 | 14 |
| 523.1 | Gingivitis | 10 | digestive | 3 | 3 | 3 |
| 523.3 | Periodontitis (acute or chronic) | 10 | digestive | 8 | 8 | 8 |
| 523.31 | Acute periodontitis | 10 | digestive | 8 | 8 | 8 |
| 523.32 | Chronic periodontitis | 10 | digestive | 0 | 0 | 0 |
| 524 | Dentofacial anomalies, including malocclusion | 10 | digestive | 2 | 2 | 2 |
| 524.3 | Anomalies of tooth position/malocclusion | 10 | digestive | 0 | 0 | 0 |
| 525 | Other diseases of the teeth and supporting structures | 10 | digestive | 37 | 37 | 37 |
| 525.1 | Loss of teeth or edentulism | 10 | digestive | 7 | 7 | 7 |
| 525.2 | Atrophy of edentulous alveolar ridge | 10 | digestive | 0 | 0 | 0 |
| 526 | Diseases of the jaws | 10 | digestive | 52 | 52 | 52 |
| 526.1 | Cysts of the jaws | 10 | digestive | 0 | 0 | 0 |

| | | | | | | |
|---|---|---|---|---|---|---|
| 526.3 | Anomalies of jaw size/symmetry | 10 | digestive | 1 | 1 | 1 |
| 526.4 | Temporomandibular joint disorders | 10 | digestive | 37 | 37 | 37 |
| 526.41 | Temporomandibular joint disorder, unspecified | 10 | digestive | 1 | 1 | 1 |
| 526.42 | Arthralgia/ankylosis of temporomandibular joint | 10 | digestive | 10 | 10 | 10 |
| 526.5 | Inflammatory conditions of jaw | 10 | digestive | 1 | 1 | 1 |
| 526.8 | Exostosis of jaw | 10 | digestive | 0 | 0 | 0 |
| 526.9 | Jaw disease NOS | 10 | digestive | 0 | 0 | 0 |
| 527 | Diseases of the salivary glands | 10 | digestive | 26 | 26 | 26 |
| 527.1 | Hypertrophy of salivary gland | 10 | digestive | 1 | 1 | 1 |
| 527.2 | Sialoadenitis | 10 | digestive | 11 | 11 | 11 |
| 527.7 | Disturbance of salivary secretion | 10 | digestive | 12 | 12 | 12 |
| 527.8 | Other specified diseases of the salivary glands | 10 | digestive | 2 | 2 | 2 |
| 528 | Diseases of the oral soft tissues, excluding lesions specific for gingiva and tongue | 10 | digestive | 61 | 61 | 61 |
| 528.1 | Stomatitis and mucositis | 10 | digestive | 18 | 18 | 18 |
| 528.11 | Stomatitis and mucositis (ulcerative) | 10 | digestive | 0 | 0 | 0 |
| 528.12 | Oral aphthae | 10 | digestive | 14 | 14 | 14 |
| 528.3 | Cellulitis and abscess of oral soft tissues | 10 | digestive | 2 | 2 | 2 |
| 528.4 | Cysts of oral soft tissues | 10 | digestive | 0 | 0 | 0 |
| 528.41 | Cyst of the salivary gland | 10 | digestive | 0 | 0 | 0 |
| 528.5 | Diseases of lips | 10 | digestive | 9 | 9 | 9 |
| 528.6 | Leukoplakia of oral mucosa | 10 | digestive | 2 | 2 | 2 |
| 528.7 | Sialolithiasis | 10 | digestive | 1 | 1 | 1 |
| 529 | Diseases and other conditions of the tongue | 10 | digestive | 19 | 19 | 19 |
| 529.1 | Glossitis | 10 | digestive | 1 | 1 | 1 |
| 529.6 | Glossodynia | 10 | digestive | 7 | 7 | 7 |
| 530 | Diseases of esophagus | 10 | digestive | 777 | 777 | 777 |
| 530.1 | Esophagitis, GERD and related diseases | 10 | digestive | 753 | 753 | 753 |
| 530.11 | GERD | 10 | digestive | 676 | 676 | 676 |
| 530.12 | Ulcer of esophagus | 10 | digestive | 0 | 0 | 0 |
| 530.13 | Barrett's esophagus | 10 | digestive | 34 | 34 | 34 |
| 530.14 | Reflux esophagitis | 10 | digestive | 91 | 91 | 91 |
| 530.15 | Eosinophilic esophagitis | 10 | digestive | 10 | 10 | 10 |
| 530.2 | Esophageal bleeding (varices/hemorrhage) | 10 | digestive | 5 | 5 | 5 |
| 530.3 | Stricture and stenosis of esophagus | 10 | digestive | 11 | 11 | 11 |
| 530.5 | Disorders of esophageal motility | 10 | digestive | 8 | 8 | 8 |
| 530.6 | Diverticulum of esophagus, acquired | 10 | digestive | 0 | 0 | 0 |
| 530.7 | Gastroesophageal laceration-hemorrhage syndrome | 10 | digestive | 0 | 0 | 0 |
| 530.9 | Heartburn | 10 | digestive | 54 | 54 | 54 |
| 531 | Peptic ulcer (excl. esophageal) | 10 | digestive | 35 | 35 | 35 |
| 531.1 | Hemorrhage from gastrointestinal ulcer | 10 | digestive | 2 | 2 | 2 |
| 531.2 | Gastric ulcer | 10 | digestive | 6 | 6 | 6 |
| 531.3 | Duodenal ulcer | 10 | digestive | 0 | 0 | 0 |
| 531.4 | Peptic ulcer, site unspecified | 10 | digestive | 27 | 27 | 27 |
| 531.5 | Gastrojejunal ulcer | 10 | digestive | 0 | 0 | 0 |
| 532 | Dysphagia | 10 | digestive | 122 | 122 | 122 |
| 535 | Gastritis and duodenitis | 10 | digestive | 62 | 62 | 62 |
| 535.1 | Acute gastritis | 10 | digestive | 0 | 0 | 0 |
| 535.2 | Atrophic gastritis | 10 | digestive | 4 | 4 | 4 |
| 535.6 | Duodenitis | 10 | digestive | 3 | 3 | 3 |
| 535.8 | Other specified gastritis | 10 | digestive | 1 | 1 | 1 |
| 535.9 | Gastritis and duodenitis, NOS | 10 | digestive | 42 | 42 | 42 |
| 536 | Disorders of function of stomach | 10 | digestive | 104 | 104 | 104 |
| 536.3 | Gastroparesis | 10 | digestive | 13 | 13 | 13 |
| 536.7 | Complications of gastrostomy, colostomy and enterostomy | 10 | digestive | 1 | 1 | 1 |
| 536.8 | Dyspepsia and other specified disorders of function of stomach | 10 | digestive | 89 | 89 | 89 |
| 537 | Other disorders of stomach and duodenum | 10 | digestive | 5 | 5 | 5 |
| 537.1 | Lesions of stomach and duodenum | 10 | digestive | 1 | 1 | 1 |
| 539 | Bariatric surgery | 10 | digestive | 83 | 83 | 83 |
| 540 | Appendiceal conditions | 10 | digestive | 36 | 36 | 36 |
| 540.1 | Appendicitis | 10 | digestive | 35 | 35 | 35 |
| 540.11 | Acute appendicitis | 10 | digestive | 22 | 22 | 22 |
| 550 | Abdominal hernia | 10 | digestive | 256 | 256 | 256 |
| 550.1 | Inguinal hernia | 10 | digestive | 60 | 60 | 60 |
| 550.2 | Diaphragmatic hernia | 10 | digestive | 103 | 103 | 103 |
| 550.3 | Femoral hernia | 10 | digestive | 0 | 0 | 0 |
| 550.4 | Umbilical hernia | 10 | digestive | 39 | 39 | 39 |
| 550.5 | Ventral hernia | 10 | digestive | 25 | 25 | 25 |

| | | | | | | |
|---|---|---|---|---|---|---|
| 550.6 | Incisional hernia | 10 | digestive | 18 | 18 | 18 |
| 555 | Inflammatory bowel disease and other gastroenteritis and colitis | 10 | digestive | 57 | 57 | 57 |
| 555.1 | Regional enteritis | 10 | digestive | 29 | 29 | 29 |
| 555.2 | Ulcerative colitis | 10 | digestive | 31 | 31 | 31 |
| 555.21 | Ulcerative colitis (chronic) | 10 | digestive | 9 | 9 | 9 |
| 556 | Ulceration of the lower GI tract | 10 | digestive | 4 | 4 | 4 |
| 556.1 | Ulceration of intestine | 10 | digestive | 4 | 4 | 4 |
| 556.11 | Angiodysplasia of intestine (without mention of hemorrhage) | 10 | digestive | 4 | 4 | 4 |
| 557 | Intestinal malabsorption (non-celiac) | 10 | digestive | 14 | 14 | 14 |
| 557.1 | Celiac disease | 10 | digestive | 0 | 0 | 0 |
| 558 | Noninfectious gastroenteritis | 10 | digestive | 158 | 158 | 158 |
| 559 | Ileostomy status | 10 | digestive | 7 | 7 | 7 |
| 560 | Intestinal obstruction without mention of hernia | 10 | digestive | 47 | 47 | 47 |
| 560.1 | Paralytic ileus | 10 | digestive | 13 | 13 | 13 |
| 560.2 | Impaction of intestine | 10 | digestive | 2 | 2 | 2 |
| 560.3 | Peritoneal or intestinal adhesions | 10 | digestive | 2 | 2 | 2 |
| 560.4 | Other intestinal obstruction | 10 | digestive | 30 | 30 | 30 |
| 561 | Symptoms involving digestive system | 10 | digestive | 43 | 43 | 43 |
| 562 | Diverticulosis and diverticulitis | 10 | digestive | 158 | 158 | 158 |
| 562.1 | Diverticulosis | 10 | digestive | 20 | 20 | 20 |
| 562.2 | Diverticulitis | 10 | digestive | 80 | 80 | 80 |
| 564 | Functional digestive disorders | 10 | digestive | 209 | 209 | 209 |
| 564.1 | Irritable Bowel Syndrome | 10 | digestive | 93 | 93 | 93 |
| 564.9 | Personal history of diseases of digestive system | 10 | digestive | 1 | 1 | 1 |
| 565 | Anal and rectal conditions | 10 | digestive | 34 | 34 | 34 |
| 565.1 | Anal and rectal polyp | 10 | digestive | 0 | 0 | 0 |
| 567 | Peritonitis and retroperitoneal infections | 10 | digestive | 8 | 8 | 8 |
| 568 | Other disorders of peritoneum | 10 | digestive | 7 | 7 | 7 |
| 568.1 | Peritoneal adhesions (postoperative) (postinfection) | 10 | digestive | 0 | 0 | 0 |
| 569 | Other disorders of intestine | 10 | digestive | 29 | 29 | 29 |
| 569.1 | Toxic gastroenteritis and colitis | 10 | digestive | 0 | 0 | 0 |
| 569.2 | Gastrointestinal complications | 10 | digestive | 5 | 5 | 5 |
| 571 | Chronic liver disease and cirrhosis | 10 | digestive | 98 | 98 | 98 |
| 571.5 | Other chronic nonalcoholic liver disease | 10 | digestive | 91 | 91 | 91 |
| 571.51 | Cirrhosis of liver without mention of alcohol | 10 | digestive | 9 | 9 | 9 |
| 571.6 | Primary biliary cirrhosis | 10 | digestive | 5 | 5 | 5 |
| 571.8 | Liver abscess and sequelae of chronic liver disease | 10 | digestive | 8 | 8 | 8 |
| 571.81 | Portal hypertension | 10 | digestive | 4 | 4 | 4 |
| 572 | Ascites (non malignant) | 10 | digestive | 6 | 6 | 6 |
| 573 | Other disorders of liver | 10 | digestive | 40 | 40 | 40 |
| 573.1 | Chronic passive congestion of liver | 10 | digestive | 0 | 0 | 0 |
| 573.2 | Liver replaced by transplant | 10 | digestive | 2 | 2 | 2 |
| 573.3 | Hepatomegaly | 10 | digestive | 11 | 11 | 11 |
| 573.4 | Acute and subacute necrosis of liver | 10 | digestive | 3 | 3 | 3 |
| 573.5 | Jaundice (not of newborn) | 10 | digestive | 9 | 9 | 9 |
| 573.6 | Nonspecific elevation of levels of transaminase or lactic acid dehydrogenase [LDH] | 10 | digestive | 71 | 71 | 71 |
| 573.7 | Abnormal results of function study of liver | 10 | digestive | 19 | 19 | 19 |
| 573.9 | Abnormal serum enzyme levels | 10 | digestive | 110 | 110 | 110 |
| 574 | Cholelithiasis and cholecystitis | 10 | digestive | 122 | 122 | 122 |
| 574.1 | Cholelithiasis | 10 | digestive | 88 | 88 | 88 |
| 574.11 | Cholelithiasis with acute cholecystitis | 10 | digestive | 1 | 1 | 1 |
| 574.12 | Cholelithiasis with other cholecystitis | 10 | digestive | 6 | 6 | 6 |
| 574.2 | Calculus of bile duct | 10 | digestive | 16 | 16 | 16 |
| 574.3 | Cholecystitis without cholelithiasis | 10 | digestive | 9 | 9 | 9 |
| 575 | Other biliary tract disease | 10 | digestive | 61 | 61 | 61 |
| 575.1 | Cholangitis | 10 | digestive | 1 | 1 | 1 |
| 575.2 | Obstruction of bile duct | 10 | digestive | 1 | 1 | 1 |
| 575.6 | Cholesterolosis of gallbladder | 10 | digestive | 7 | 7 | 7 |
| 575.7 | Other disorders of gallbladder | 10 | digestive | 26 | 26 | 26 |
| 575.8 | Other disorders of biliary tract | 10 | digestive | 6 | 6 | 6 |
| 575.9 | Nonspecific abnormal findings on radiological and other examination of biliary tract | 10 | digestive | 5 | 5 | 5 |
| 577 | Diseases of pancreas | 10 | digestive | 57 | 57 | 57 |
| 577.1 | Acute pancreatitis | 10 | digestive | 11 | 11 | 11 |
| 577.2 | Chronic pancreatitis | 10 | digestive | 3 | 3 | 3 |
| 577.3 | Cyst and pseudocyst of pancreas | 10 | digestive | 11 | 11 | 11 |
| 578 | Gastrointestinal hemorrhage | 10 | digestive | 128 | 128 | 128 |
| 578.1 | Hematemesis | 10 | digestive | 0 | 0 | 0 |

| | | | | | | |
|---|---|---|---|---|---|---|
| 578.2 | Blood in stool | 10 | digestive | 45 | 45 | 45 |
| 578.8 | Hemorrhage of rectum and anus | 10 | digestive | 48 | 48 | 48 |
| 578.9 | Hemorrhage of gastrointestinal tract | 10 | digestive | 33 | 33 | 33 |
| 579 | Other symptoms involving abdomen and pelvis | 10 | digestive | 327 | 327 | 327 |
| 579.2 | Splenomegaly | 10 | digestive | 15 | 15 | 15 |
| 579.8 | Nonspecific abnormal findings in stool contents | 10 | digestive | 23 | 23 | 23 |
| 580 | Nephritis; nephrosis; renal sclerosis | 11 | genitourinary | 17 | 17 | 17 |
| 580.1 | Glomerulonephritis | 11 | genitourinary | 1 | 1 | 1 |
| 580.11 | Proliferative glomerulonephritis | 11 | genitourinary | 0 | 0 | 0 |
| 580.12 | Non-proliferative glomerulonephritis | 11 | genitourinary | 1 | 1 | 1 |
| 580.13 | Acute glomerulonephritis, NOS | 11 | genitourinary | 0 | 0 | 0 |
| 580.14 | Chronic glomerulonephritis, NOS | 11 | genitourinary | 0 | 0 | 0 |
| 580.2 | Nephrotic syndrome without mention of glomerulonephritis | 11 | genitourinary | 0 | 0 | 0 |
| 580.3 | Nephritis and nephropathy without mention of glomerulonephritis | 11 | genitourinary | 14 | 14 | 14 |
| 580.31 | Nephritis and nephropathy in diseases classified elsewhere | 11 | genitourinary | 11 | 11 | 11 |
| 580.32 | Nephritis and nephropathy with pathological lesion | 11 | genitourinary | 2 | 2 | 2 |
| 580.4 | Renal sclerosis, NOS | 11 | genitourinary | 2 | 2 | 2 |
| 585 | Renal failure | 11 | genitourinary | 223 | 223 | 223 |
| 585.1 | Acute renal failure | 11 | genitourinary | 78 | 78 | 78 |
| 585.2 | Renal failure NOS | 11 | genitourinary | 6 | 6 | 6 |
| 585.3 | Chronic renal failure [CKD] | 11 | genitourinary | 141 | 141 | 141 |
| 585.31 | Renal dialysis | 11 | genitourinary | 1 | 1 | 1 |
| 585.32 | End stage renal disease | 11 | genitourinary | 2 | 2 | 2 |
| 585.33 | Chronic Kidney Disease, Stage III | 11 | genitourinary | 120 | 120 | 120 |
| 585.34 | Chronic Kidney Disease, Stage IV | 11 | genitourinary | 10 | 10 | 10 |
| 585.4 | Chronic kidney disease, Stage I or II | 11 | genitourinary | 42 | 42 | 42 |
| 586 | Other disorders of the kidney and ureters | 11 | genitourinary | 177 | 177 | 177 |
| 586.1 | Anatomical abnormatilies of kidney and ureters | 11 | genitourinary | 0 | 0 | 0 |
| 586.11 | Small kidney | 11 | genitourinary | 0 | 0 | 0 |
| 586.12 | Vesicoureteral reflux | 11 | genitourinary | 0 | 0 | 0 |
| 586.2 | Cyst of kidney, acquired | 11 | genitourinary | 27 | 27 | 27 |
| 586.3 | Vascular disorders of kidney/hypertrophy | 11 | genitourinary | 2 | 2 | 2 |
| 586.4 | Stricture/obstruction of ureter | 11 | genitourinary | 5 | 5 | 5 |
| 587 | Kidney replaced by transpant | 11 | genitourinary | 0 | 0 | 0 |
| 588 | Disorders resulting from impaired renal function | 11 | genitourinary | 4 | 4 | 4 |
| 588.1 | Renal osteodystrophy | 11 | genitourinary | 0 | 0 | 0 |
| 588.2 | Secondary hyperparathyroidism (of renal origin) | 11 | genitourinary | 4 | 4 | 4 |
| 589 | Abnormal results of function study of kidney | 11 | genitourinary | 25 | 25 | 25 |
| 590 | Pyelonephritis | 11 | genitourinary | 52 | 52 | 52 |
| 591 | Urinary tract infection | 11 | genitourinary | 87 | 87 | 87 |
| 592 | Cystitis and urethritis | 11 | genitourinary | 214 | 214 | 214 |
| 592.1 | Cystitis | 11 | genitourinary | 72 | 72 | 72 |
| 592.11 | Acute cystitis | 11 | genitourinary | 16 | 16 | 16 |
| 592.12 | Chronic cystitis | 11 | genitourinary | 1 | 1 | 1 |
| 592.13 | Chronic interstitial cystitis | 11 | genitourinary | 13 | 13 | 13 |
| 592.2 | Urethritis and urethral syndrome | 11 | genitourinary | 12 | 12 | 12 |
| 592.21 | Urethral syndrome | 11 | genitourinary | 7 | 7 | 7 |
| 592.3 | Urethral stricture due to infecton | 11 | genitourinary | 0 | 0 | 0 |
| 593 | Hematuria | 11 | genitourinary | 174 | 174 | 174 |
| 593.1 | Gross hematuria | 11 | genitourinary | 21 | 21 | 21 |
| 593.2 | Microscopic hematuria | 11 | genitourinary | 40 | 40 | 40 |
| 594 | Urinary calculus | 11 | genitourinary | 217 | 217 | 217 |
| 594.1 | Calculus of kidney | 11 | genitourinary | 14 | 14 | 14 |
| 594.2 | Calculus of lower urinary tract | 11 | genitourinary | 2 | 2 | 2 |
| 594.3 | Calculus of ureter | 11 | genitourinary | 51 | 51 | 51 |
| 594.8 | Renal colic | 11 | genitourinary | 0 | 0 | 0 |
| 595 | Hydronephrosis | 11 | genitourinary | 28 | 28 | 28 |
| 596 | Other disorders of bladder | 11 | genitourinary | 53 | 53 | 53 |
| 596.1 | Bladder neck obstruction | 11 | genitourinary | 0 | 0 | 0 |
| 596.5 | Functional disorders of bladder | 11 | genitourinary | 31 | 31 | 31 |
| 597 | Other disorders of urethra and urinary tract | 11 | genitourinary | 503 | 503 | 503 |
| 597.1 | Urethral stricture (not specified as infectious) | 11 | genitourinary | 1 | 1 | 1 |
| 597.2 | Urinary complications NEC | 11 | genitourinary | 1 | 1 | 1 |
| 597.8 | Urethral hypermobility/ISD | 11 | genitourinary | 0 | 0 | 0 |
| 598 | Abnormal findings on examination of urine | 11 | genitourinary | 26 | 26 | 26 |
| 598.4 | Other cells and casts in urine | 11 | genitourinary | 8 | 8 | 8 |
| 599 | Other symptoms/disorders or the urinary system | 11 | genitourinary | 657 | 657 | 657 |

| | | | | | | |
|---|---|---|---|---|---|---|
| 599.1 | Urinary obstruction | 11 | genitourinary | 12 | 12 | 12 |
| 599.2 | Retention of urine | 11 | genitourinary | 48 | 48 | 48 |
| 599.3 | Dysuria | 11 | genitourinary | 254 | 254 | 254 |
| 599.4 | Urinary incontinence | 11 | genitourinary | 97 | 97 | 97 |
| 599.5 | Frequency of urination and polyuria | 11 | genitourinary | 142 | 142 | 142 |
| 599.6 | Oliguria and anuria | 11 | genitourinary | 0 | 0 | 0 |
| 599.7 | Urethral discharge | 11 | genitourinary | 4 | 4 | 4 |
| 599.8 | Other symptoms involving urinary system | 11 | genitourinary | 52 | 52 | 52 |
| 599.9 | Other abnormality of urination | 11 | genitourinary | 55 | 55 | 55 |
| 600 | Hyperplasia of prostate | 11 | genitourinary | 170 | 170 | 170 |
| 601 | Inflammatory diseases of prostate | 11 | genitourinary | 23 | 23 | 23 |
| 601.1 | Prostatitis | 11 | genitourinary | 4 | 4 | 4 |
| 601.11 | Acute prostatitis | 11 | genitourinary | 0 | 0 | 0 |
| 601.12 | Chronic prostatitis | 11 | genitourinary | 1 | 1 | 1 |
| 601.3 | Orchitis and epididymitis | 11 | genitourinary | 9 | 9 | 9 |
| 601.4 | Balanoposthitis | 11 | genitourinary | 4 | 4 | 4 |
| 601.8 | Other inflammatory disorders of male genital organs | 11 | genitourinary | 1 | 1 | 1 |
| 602 | Other disorders of prostate | 11 | genitourinary | 4 | 4 | 4 |
| 602.3 | Dysplasia of prostate | 11 | genitourinary | 0 | 0 | 0 |
| 603 | Other disorders of testis | 11 | genitourinary | 12 | 12 | 12 |
| 603.1 | Hydrocele | 11 | genitourinary | 8 | 8 | 8 |
| 603.2 | Spermatocele | 11 | genitourinary | 2 | 2 | 2 |
| 604 | Disorders of penis | 11 | genitourinary | 9 | 9 | 9 |
| 604.1 | Redundant prepuce and phimosis/BXO | 11 | genitourinary | 0 | 0 | 0 |
| 604.2 | Vascular disorders of penis | 11 | genitourinary | 0 | 0 | 0 |
| 604.3 | Peyronie's disease | 11 | genitourinary | 2 | 2 | 2 |
| 605 | Erectile dysfunction [ED] | 11 | genitourinary | 113 | 113 | 113 |
| 608 | Other disorders of male genital organs | 11 | genitourinary | 30 | 30 | 30 |
| 609 | Male infertility and abnormal spermatozoa | 11 | genitourinary | 3 | 3 | 3 |
| 609.1 | Infertility, male | 11 | genitourinary | 1 | 1 | 1 |
| 609.11 | Azoospermia and oligospermia | 11 | genitourinary | 0 | 0 | 0 |
| 609.2 | Abnormal spermatozoa | 11 | genitourinary | 2 | 2 | 2 |
| 610 | Benign mammary dysplasias | 11 | genitourinary | 78 | 78 | 78 |
| 610.1 | Cystic mastopathy | 11 | genitourinary | 35 | 35 | 35 |
| 610.2 | Fibroadenosis of breast | 11 | genitourinary | 1 | 1 | 1 |
| 610.3 | Fibrosclerosis of breast | 11 | genitourinary | 1 | 1 | 1 |
| 610.4 | Benign neoplasm of breast | 11 | genitourinary | 9 | 9 | 9 |
| 610.8 | Other specified benign mammary dysplasias | 11 | genitourinary | 6 | 6 | 6 |
| 611 | Abnormal findings on mammogram or breast exam | 11 | genitourinary | 574 | 574 | 574 |
| 611.1 | Abnormal mammogram | 11 | genitourinary | 249 | 249 | 249 |
| 611.11 | Mammographic microcalcification | 11 | genitourinary | 22 | 22 | 22 |
| 611.3 | Lump or mass in breast | 11 | genitourinary | 204 | 204 | 204 |
| 612 | Breast conditions, congenital or relating to hormones | 11 | genitourinary | 31 | 31 | 31 |
| 612.1 | Galactorrhea | 11 | genitourinary | 4 | 4 | 4 |
| 612.2 | Hypertrophy of breast (Gynecomastia) | 11 | genitourinary | 23 | 23 | 23 |
| 612.3 | Congenital anomalies of breast | 11 | genitourinary | 0 | 0 | 0 |
| 613 | Other nonmalignant breast conditions | 11 | genitourinary | 180 | 180 | 180 |
| 613.1 | Inflammatory disease of breast | 11 | genitourinary | 1 | 1 | 1 |
| 613.5 | Mastodynia | 11 | genitourinary | 97 | 97 | 97 |
| 613.7 | Other signs and symptoms in breast | 11 | genitourinary | 28 | 28 | 28 |
| 613.8 | Other specified disorders of breast | 11 | genitourinary | 11 | 11 | 11 |
| 613.9 | Breast disorder NOS | 11 | genitourinary | 10 | 10 | 10 |
| 614 | Inflammatory diseases of female pelvic organs | 11 | genitourinary | 178 | 178 | 178 |
| 614.1 | Pelvic peritoneal adhesions, female (postoperative) (postinfection) | 11 | genitourinary | 0 | 0 | 0 |
| 614.3 | Pelvic inflammatory disease (PID) | 11 | genitourinary | 5 | 5 | 5 |
| 614.31 | Acute inflammatory pelvic disease | 11 | genitourinary | 3 | 3 | 3 |
| 614.32 | Chronic inflammatory pelvic disease | 11 | genitourinary | 0 | 0 | 0 |
| 614.33 | Pelvic inflammatory disease, NOS | 11 | genitourinary | 1 | 1 | 1 |
| 614.4 | Inflammatory diseases of uterus, except cervix | 11 | genitourinary | 3 | 3 | 3 |
| 614.5 | Inflammatory disease of cervix, vagina, and vulva | 11 | genitourinary | 167 | 167 | 167 |
| 614.51 | Cervicitis and endocervicitis | 11 | genitourinary | 0 | 0 | 0 |
| 614.52 | Vaginitis and vulvovaginitis | 11 | genitourinary | 146 | 146 | 146 |
| 614.53 | Cyst or abscess of Bartholin's gland | 11 | genitourinary | 4 | 4 | 4 |
| 614.54 | Abscess or ulceration of vulva | 11 | genitourinary | 8 | 8 | 8 |
| 615 | Endometriosis | 11 | genitourinary | 41 | 41 | 41 |
| 618 | Genital prolapse | 11 | genitourinary | 49 | 49 | 49 |
| 618.1 | Prolapse of vaginal walls | 11 | genitourinary | 30 | 30 | 30 |

| | | | | | | |
|---|---|---|---|---|---|---|
| 618.2 | Uterine/Uterovaginal prolapse | 11 | genitourinary | 4 | 4 | 4 |
| 618.5 | Prolapse of vaginal vault after hysterectomy | 11 | genitourinary | 0 | 0 | 0 |
| 618.6 | Vaginal enterocele, congenital or acquired | 11 | genitourinary | 0 | 0 | 0 |
| 619 | Noninflammatory female genital disorders | 11 | genitourinary | 133 | 133 | 133 |
| 619.1 | Noninflammatory disorders of ovary, fallopian tube, and broad ligament | 11 | genitourinary | 5 | 5 | 5 |
| 619.2 | Disorders of uterus, NEC | 11 | genitourinary | 9 | 9 | 9 |
| 619.3 | Noninflammatory disorders of cervix | 11 | genitourinary | 2 | 2 | 2 |
| 619.4 | Noninflammatory disorders of vagina | 11 | genitourinary | 31 | 31 | 31 |
| 619.5 | Noninflammatory disorders of vulva and perineum | 11 | genitourinary | 7 | 7 | 7 |
| 620 | Dysplasia of female genital organs | 11 | genitourinary | 6 | 6 | 6 |
| 620.1 | Dysplasia of cervix | 11 | genitourinary | 3 | 3 | 3 |
| 621 | Endometrial hyperplasia | 11 | genitourinary | 6 | 6 | 6 |
| 622 | Polyp of female genital organs | 11 | genitourinary | 6 | 6 | 6 |
| 622.1 | Polyp of corpus uteri | 11 | genitourinary | 0 | 0 | 0 |
| 622.2 | Mucous polyp of cervix | 11 | genitourinary | 5 | 5 | 5 |
| 623 | Hypertrophy of female genital organs | 11 | genitourinary | 14 | 14 | 14 |
| 624 | Symptoms involving female genital tract | 11 | genitourinary | 36 | 36 | 36 |
| 624.1 | Dystrophy of female genital tract | 11 | genitourinary | 1 | 1 | 1 |
| 624.2 | Atrophy of female genital tract | 11 | genitourinary | 1 | 1 | 1 |
| 624.9 | stress incontinence, female | 11 | genitourinary | 5 | 5 | 5 |
| 625 | Pain and other symptoms associated with female genital organs | 11 | genitourinary | 208 | 208 | 208 |
| 625.1 | Dyspareunia | 11 | genitourinary | 2 | 2 | 2 |
| 626 | Disorders of menstruation and other abnormal bleeding from female genital tract | 11 | genitourinary | 435 | 435 | 435 |
| 626.1 | Irregular menstrual cycle/bleeding | 11 | genitourinary | 315 | 315 | 315 |
| 626.11 | Absent or infrequent menstruation | 11 | genitourinary | 16 | 16 | 16 |
| 626.12 | Excessive or frequent menstruation | 11 | genitourinary | 111 | 111 | 111 |
| 626.13 | Irregular menstrual cycle | 11 | genitourinary | 48 | 48 | 48 |
| 626.14 | Irregular menstrual bleeding | 11 | genitourinary | 17 | 17 | 17 |
| 626.15 | Infertility, female, associated with anovulation | 11 | genitourinary | 0 | 0 | 0 |
| 626.2 | Dysmenorrhea | 11 | genitourinary | 47 | 47 | 47 |
| 626.21 | Mittelschmerz | 11 | genitourinary | 3 | 3 | 3 |
| 626.4 | Premenstrual tension syndromes | 11 | genitourinary | 15 | 15 | 15 |
| 626.8 | Infertility, female | 11 | genitourinary | 23 | 23 | 23 |
| 627 | Menopausal and postmenopausal disorders | 11 | genitourinary | 419 | 419 | 419 |
| 627.1 | Postmenopausal bleeding | 11 | genitourinary | 35 | 35 | 35 |
| 627.2 | Symptomatic menopause | 11 | genitourinary | 269 | 269 | 269 |
| 627.21 | Symptomatic artificial menopause | 11 | genitourinary | 3 | 3 | 3 |
| 627.22 | Need for Hormone replacement therapy (postmenopausal) | 11 | genitourinary | 92 | 92 | 92 |
| 627.3 | Postmenopausal atrophic vaginitis | 11 | genitourinary | 56 | 56 | 56 |
| 627.4 | Premenopausal menorrhagia | 11 | genitourinary | 0 | 0 | 0 |
| 627.5 | Premature menopause and other ovarian failure | 11 | genitourinary | 10 | 10 | 10 |
| 628 | Ovarian cyst | 11 | genitourinary | 122 | 122 | 122 |
| 634 | Miscarriage; stillbirth | 12 | pregnancy complications | 34 | 34 | 34 |
| 634.1 | Missed abortion/Hydatidiform mole | 12 | pregnancy complications | 4 | 4 | 4 |
| 634.3 | Ectopic pregnancy | 12 | pregnancy complications | 7 | 7 | 7 |
| 635 | Hemorrhage during pregnancy; childbirth and postpartum | 12 | pregnancy complications | 11 | 11 | 11 |
| 635.2 | Antepartum hemorrhage, abruptio placentae, and placenta previa | 12 | pregnancy complications | 11 | 11 | 11 |
| 635.3 | Placenta previa and abruptio placenta | 12 | pregnancy complications | 0 | 0 | 0 |
| 636 | Early or threatened labor; hemorrhage in early pregnancy | 12 | pregnancy complications | 82 | 82 | 82 |
| 636.1 | Threatened premature labor | 12 | pregnancy complications | 2 | 2 | 2 |
| 636.2 | Early onset of delivery | 12 | pregnancy complications | 4 | 4 | 4 |
| 636.3 | Hemorrhage in early pregnancy | 12 | pregnancy complications | 44 | 44 | 44 |
| 636.8 | Cervical incompetence | 12 | pregnancy complications | 3 | 3 | 3 |
| 637 | Short gestation; low birth weight; and fetal growth retardation | 12 | pregnancy complications | 0 | 0 | 0 |
| 642 | Hypertension complicating pregnancy, childbirth, and the puerperium | 12 | pregnancy complications | 32 | 32 | 32 |
| 642.1 | Preeclampsia and eclampsia | 12 | pregnancy complications | 9 | 9 | 9 |
| 643 | Excessive vomiting in pregnancy | 12 | pregnancy complications | 12 | 12 | 12 |
| 643.1 | Hyperemesis gravidarum | 12 | pregnancy complications | 1 | 1 | 1 |
| 644 | Anemia during pregnancy | 12 | pregnancy complications | 3 | 3 | 3 |
| 646 | Other complications of pregnancy NEC | 12 | pregnancy complications | 29 | 29 | 29 |
| 647 | Infectious and parasitic complications affecting pregnancy | 12 | pregnancy complications | 6 | 6 | 6 |
| 647.1 | Infections of genitourinary tract during pregnancy | 12 | pregnancy complications | 3 | 3 | 3 |
| 647.3 | Major puerperal infection | 12 | pregnancy complications | 0 | 0 | 0 |
| 649 | Other conditions or status of the mother complicating pregnancy, childbirth, or the puerperium | 12 | pregnancy complications | 46 | 46 | 46 |
| 649.1 | Diabetes or abnormal glucose tolerance complicating pregnancy | 12 | pregnancy complications | 27 | 27 | 27 |
| 653 | Problems associated with amniotic cavity and membranes | 12 | pregnancy complications | 18 | 18 | 18 |
| 654 | Other and unspecified complications of birth; puerperium affecting management of mother | 12 | pregnancy complications | 0 | 0 | 0 |

| | | | | | | |
|---|---|---|---|---|---|---|
| 654.1 | Abnormality of organs and soft tissues of pelvis complicating pregnancy, childbirth, or the puerperium | 12 | pregnancy complications | 4 | 4 | 4 |
| 654.2 | Rhesus isoimmunization in pregnancy | 12 | pregnancy complications | 4 | 4 | 4 |
| 655 | Known or suspected fetal abnormality affecting management of mother | 12 | pregnancy complications | 75 | 75 | 75 |
| 655.1 | Abnormality in fetal heart rate or rhythm | 12 | pregnancy complications | 1 | 1 | 1 |
| 656 | Other perinatal conditions of fetus or newborn | 12 | pregnancy complications | 0 | 0 | 0 |
| 656.1 | Isoimmunization of fetus or newborn | 12 | pregnancy complications | 0 | 0 | 0 |
| 656.2 | Respiratory conditions of fetus and newborn | 12 | pregnancy complications | 0 | 0 | 0 |
| 656.22 | Interstitial emphysema and related conditions of newborn | 12 | pregnancy complications | 0 | 0 | 0 |
| 656.26 | Transitory tachypnea or apnea of newborn | 12 | pregnancy complications | 0 | 0 | 0 |
| 656.3 | Endocrine and metabolic disturbances of fetus and newborn | 12 | pregnancy complications | 0 | 0 | 0 |
| 656.4 | Hemorrhage of fetus or newborn | 12 | pregnancy complications | 0 | 0 | 0 |
| 656.5 | Hematological disorders of newborn | 12 | pregnancy complications | 0 | 0 | 0 |
| 656.6 | Perinatal disorders of digestive system | 12 | pregnancy complications | 0 | 0 | 0 |
| 656.7 | Conditions involving the integument and temperature regulation of fetus and newborn | 12 | pregnancy complications | 0 | 0 | 0 |
| 656.8 | Perinatal jaundice | 12 | pregnancy complications | 0 | 0 | 0 |
| 656.9 | Neonatal bradycardia or tachycardia | 12 | pregnancy complications | 0 | 0 | 0 |
| 657 | Infections specific to the perinatal period | 12 | pregnancy complications | 0 | 0 | 0 |
| 669 | Complications of labor and delivery NEC | 12 | pregnancy complications | 2 | 2 | 2 |
| 671 | Venous/cerebrovascular complications embolism in pregnancy and the puerperium | 12 | pregnancy complications | 11 | 11 | 11 |
| 674 | Other complications of the puerperium NEC | 12 | pregnancy complications | 0 | 0 | 0 |
| 676 | Other disorders of the breast associated with childbirth and disorders of lactation | 12 | pregnancy complications | 2 | 2 | 2 |
| 681 | Superficial cellulitis and abscess | 13 | dermatologic | 282 | 282 | 282 |
| 681.1 | Cellulitis and abscess of fingers/toes | 13 | dermatologic | 35 | 35 | 35 |
| 681.2 | Cellulitis and abscess of face/neck | 13 | dermatologic | 8 | 8 | 8 |
| 681.3 | Cellulitis and abscess of arm/hand | 13 | dermatologic | 22 | 22 | 22 |
| 681.5 | Cellulitis and abscess of leg, except foot | 13 | dermatologic | 44 | 44 | 44 |
| 681.6 | Cellulitis and abscess of foot, toe | 13 | dermatologic | 12 | 12 | 12 |
| 681.7 | Cellulitis and abscess of trunk | 13 | dermatologic | 55 | 55 | 55 |
| 686 | Other local infections of skin and subcutaneous tissue | 13 | dermatologic | 67 | 67 | 67 |
| 686.1 | Carbuncle and furuncle | 13 | dermatologic | 11 | 11 | 11 |
| 686.2 | Impetigo | 13 | dermatologic | 6 | 6 | 6 |
| 686.3 | Pilonidal cyst | 13 | dermatologic | 4 | 4 | 4 |
| 686.4 | Pyogenic granuloma | 13 | dermatologic | 0 | 0 | 0 |
| 686.5 | Pyoderma | 13 | dermatologic | 0 | 0 | 0 |
| 687 | Symptoms affecting skin | 13 | dermatologic | 51 | 51 | 51 |
| 687.1 | Rash and other nonspecific skin eruption | 13 | dermatologic | 224 | 224 | 224 |
| 687.3 | Changes in skin texture | 13 | dermatologic | 16 | 16 | 16 |
| 687.4 | Disturbance of skin sensation | 13 | dermatologic | 59 | 59 | 59 |
| 689 | Disorder of skin and subcutaneous tissue NOS | 13 | dermatologic | 243 | 243 | 243 |
| 690 | Erythematosquamous dermatosis | 13 | dermatologic | 11 | 11 | 11 |
| 690.1 | Seborrheic dermatitis | 13 | dermatologic | 11 | 11 | 11 |
| 691 | Congenital anomalies of skin | 13 | dermatologic | 4 | 4 | 4 |
| 691.1 | Ichthyosis congenita | 13 | dermatologic | 0 | 0 | 0 |
| 691.3 | Congenital pigmentary anomalies of skin | 13 | dermatologic | 1 | 1 | 1 |
| 694 | Dyschromia and Vitiligo | 13 | dermatologic | 29 | 29 | 29 |
| 694.1 | Vitiligo | 13 | dermatologic | 4 | 4 | 4 |
| 694.2 | Other dyschromia | 13 | dermatologic | 23 | 23 | 23 |
| 694.3 | Vascular disorders of skin | 13 | dermatologic | 2 | 2 | 2 |
| 695 | Erythematous conditions | 13 | dermatologic | 94 | 94 | 94 |
| 695.1 | Toxic erythema | 13 | dermatologic | 3 | 3 | 3 |
| 695.2 | Bullous dermatoses | 13 | dermatologic | 2 | 2 | 2 |
| 695.21 | Dermatitis herpetiformis | 13 | dermatologic | 0 | 0 | 0 |
| 695.22 | Pemphigus and pemphigoid | 13 | dermatologic | 2 | 2 | 2 |
| 695.3 | Rosacea | 13 | dermatologic | 43 | 43 | 43 |
| 695.4 | Lupus (localized and systemic) | 13 | dermatologic | 21 | 21 | 21 |
| 695.41 | Cutaneous lupus erythematosus | 13 | dermatologic | 6 | 6 | 6 |
| 695.42 | Systemic lupus erythematosus | 13 | dermatologic | 10 | 10 | 10 |
| 695.7 | Prurigo and Lichen | 13 | dermatologic | 10 | 10 | 10 |
| 695.8 | Other specified erythematous conditions | 13 | dermatologic | 10 | 10 | 10 |
| 695.81 | Erythema nodosum | 13 | dermatologic | 3 | 3 | 3 |
| 695.9 | Unspecified erythematous condition | 13 | dermatologic | 3 | 3 | 3 |
| 696 | Psoriasis and related disorders | 13 | dermatologic | 46 | 46 | 46 |
| 696.2 | Parapsoriasis | 13 | dermatologic | 0 | 0 | 0 |
| 696.3 | Pityriasis | 13 | dermatologic | 0 | 0 | 0 |
| 696.4 | Psoriasis | 13 | dermatologic | 42 | 42 | 42 |
| 696.41 | Psoriasis vulgaris | 13 | dermatologic | 40 | 40 | 40 |
| 696.42 | Psoriatic arthropathy | 13 | dermatologic | 1 | 1 | 1 |

| | | | | | | |
|---|---|---|---|---|---|---|
| 697 | Sarcoidosis | 13 | dermatologic | 13 | 13 | 13 |
| 698 | Pruritus and related conditions | 13 | dermatologic | 80 | 80 | 80 |
| 700 | Corns and callosities | 13 | dermatologic | 14 | 14 | 14 |
| 701 | Other hypertrophic and atrophic conditions of skin | 13 | dermatologic | 96 | 96 | 96 |
| 701.1 | Keratoderma, acquired | 13 | dermatologic | 1 | 1 | 1 |
| 701.2 | Scar conditions and fibrosis of skin | 13 | dermatologic | 5 | 5 | 5 |
| 701.3 | Circumscribed scleroderma | 13 | dermatologic | 0 | 0 | 0 |
| 701.4 | Keloid scar | 13 | dermatologic | 3 | 3 | 3 |
| 701.5 | Abnormal granulation tissue | 13 | dermatologic | 1 | 1 | 1 |
| 701.6 | Acquired acanthosis nigricans | 13 | dermatologic | 2 | 2 | 2 |
| 702 | Degenerative skin conditions and other dermatoses | 13 | dermatologic | 141 | 141 | 141 |
| 702.1 | Actinic keratosis | 13 | dermatologic | 0 | 0 | 0 |
| 702.2 | Seborrheic keratosis | 13 | dermatologic | 62 | 62 | 62 |
| 702.4 | Degenerative skin disorders | 13 | dermatologic | 0 | 0 | 0 |
| 703 | Diseases of nail, NOS | 13 | dermatologic | 46 | 46 | 46 |
| 703.1 | Ingrowing nail | 13 | dermatologic | 1 | 1 | 1 |
| 704 | Diseases of hair and hair follicles | 13 | dermatologic | 105 | 105 | 105 |
| 704.1 | Alopecia | 13 | dermatologic | 13 | 13 | 13 |
| 704.11 | Alopecia Areata | 13 | dermatologic | 1 | 1 | 1 |
| 704.12 | Telogen effluvium | 13 | dermatologic | 0 | 0 | 0 |
| 704.2 | Hirsutism | 13 | dermatologic | 13 | 13 | 13 |
| 704.8 | Other specified diseases of hair and hair follicles | 13 | dermatologic | 9 | 9 | 9 |
| 705 | Disorders of sweat glands | 13 | dermatologic | 16 | 16 | 16 |
| 705.1 | Dyshidrosis | 13 | dermatologic | 8 | 8 | 8 |
| 705.3 | Hidradenitis | 13 | dermatologic | 5 | 5 | 5 |
| 705.8 | Hyperhidrosis | 13 | dermatologic | 34 | 34 | 34 |
| 706 | Diseases of sebaceous glands | 13 | dermatologic | 153 | 153 | 153 |
| 706.1 | Acne | 13 | dermatologic | 96 | 96 | 96 |
| 706.2 | Sebaceous cyst | 13 | dermatologic | 53 | 53 | 53 |
| 706.3 | Seborrhea | 13 | dermatologic | 0 | 0 | 0 |
| 706.8 | Other specified diseases of sebaceous glands | 13 | dermatologic | 1 | 1 | 1 |
| 707 | Chronic ulcer of skin | 13 | dermatologic | 27 | 27 | 27 |
| 707.1 | Decubitus ulcer | 13 | dermatologic | 8 | 8 | 8 |
| 707.2 | Chronic ulcer of leg or foot | 13 | dermatologic | 15 | 15 | 15 |
| 707.3 | Chronic ulcer of unspecified site | 13 | dermatologic | 8 | 8 | 8 |
| 709 | Diffuse diseases of connective tissue | 13 | dermatologic | 40 | 40 | 40 |
| 709.2 | Sicca syndrome | 13 | dermatologic | 20 | 20 | 20 |
| 709.3 | Systemic sclerosis | 13 | dermatologic | 3 | 3 | 3 |
| 709.4 | Polymyositis | 13 | dermatologic | 1 | 1 | 1 |
| 709.5 | Dermatomyositis | 13 | dermatologic | 0 | 0 | 0 |
| 709.6 | Other specified diffuse diseases of connective tissue | 13 | dermatologic | 1 | 1 | 1 |
| 709.7 | Unspecified diffuse connective tissue disease | 13 | dermatologic | 6 | 6 | 6 |
| 710 | Osteomyelitis, periostitis, and other infections involving bone | 14 | musculoskeletal | 19 | 19 | 19 |
| 710.1 | Osteomyelitis | 14 | musculoskeletal | 19 | 19 | 19 |
| 710.11 | Acute osteomyelitis | 14 | musculoskeletal | 3 | 3 | 3 |
| 710.12 | Chronic osteomyelitis | 14 | musculoskeletal | 2 | 2 | 2 |
| 710.19 | Unspecified osteomyelitis | 14 | musculoskeletal | 14 | 14 | 14 |
| 710.2 | Periostitis | 14 | musculoskeletal | 0 | 0 | 0 |
| 710.3 | Osteopathy resulting from poliomyelitis | 14 | musculoskeletal | 0 | 0 | 0 |
| 711 | Arthropathy associated with infections | 14 | musculoskeletal | 8 | 8 | 8 |
| 711.1 | Pyogenic arthritis | 14 | musculoskeletal | 2 | 2 | 2 |
| 711.2 | Reiter's disease | 14 | musculoskeletal | 2 | 2 | 2 |
| 711.3 | Behcet's syndrome | 14 | musculoskeletal | 0 | 0 | 0 |
| 712 | Infective connective tissue disorders | 14 | musculoskeletal | 1 | 1 | 1 |
| 713 | Arthropathy associated with other disorders classified elsewhere | 14 | musculoskeletal | 3 | 3 | 3 |
| 713.5 | Arthropathy associated with neurological disorders | 14 | musculoskeletal | 0 | 0 | 0 |
| 714 | Rheumatoid arthritis and other inflammatory polyarthropathies | 14 | musculoskeletal | 73 | 73 | 73 |
| 714.1 | Rheumatoid arthritis | 14 | musculoskeletal | 37 | 37 | 37 |
| 714.2 | Juvenile rheumatoid arthritis | 14 | musculoskeletal | 1 | 1 | 1 |
| 715 | Other inflammatory spondylopathies | 14 | musculoskeletal | 34 | 34 | 34 |
| 715.1 | Sacroiliitis NEC | 14 | musculoskeletal | 12 | 12 | 12 |
| 715.2 | Ankylosing spondylitis | 14 | musculoskeletal | 2 | 2 | 2 |
| 715.3 | Spinal enthesopathy | 14 | musculoskeletal | 3 | 3 | 3 |
| 716 | Other arthropathies | 14 | musculoskeletal | 268 | 268 | 268 |
| 716.1 | Unspecified polyarthropathy or polyarthritis | 14 | musculoskeletal | 10 | 10 | 10 |
| 716.2 | Unspecified monoarthritis | 14 | musculoskeletal | 0 | 0 | 0 |
| 716.3 | Kaschin-Beck disease | 14 | musculoskeletal | 0 | 0 | 0 |

| | | | | | | |
|---|---|---|---|---|---|---|
| 716.8 | Palindromic rheumatism | 14 | musculoskeletal | 0 | 0 | 0 |
| 716.9 | Arthropathy NOS | 14 | musculoskeletal | 245 | 245 | 245 |
| 717 | Polymyalgia Rheumatica | 14 | musculoskeletal | 5 | 5 | 5 |
| 720 | Spinal stenosis | 14 | musculoskeletal | 145 | 145 | 145 |
| 720.1 | Spinal stenosis of lumbar region | 14 | musculoskeletal | 126 | 126 | 126 |
| 721 | Spondylosis and allied disorders | 14 | musculoskeletal | 291 | 291 | 291 |
| 721.1 | Spondylosis without myelopathy | 14 | musculoskeletal | 217 | 217 | 217 |
| 721.2 | Spondylosis with myelopathy | 14 | musculoskeletal | 9 | 9 | 9 |
| 721.8 | Other allied disorders of spine | 14 | musculoskeletal | 5 | 5 | 5 |
| 722 | Intervertebral disc disorders | 14 | musculoskeletal | 519 | 519 | 519 |
| 722.1 | Displacement of intervertebral disc | 14 | musculoskeletal | 162 | 162 | 162 |
| 722.3 | Schmorl's nodes | 14 | musculoskeletal | 4 | 4 | 4 |
| 722.6 | Degeneration of intervertebral disc | 14 | musculoskeletal | 328 | 328 | 328 |
| 722.7 | Intervertebral disc disorder with myelopathy | 14 | musculoskeletal | 9 | 9 | 9 |
| 722.8 | Postlaminectomy syndrome | 14 | musculoskeletal | 15 | 15 | 15 |
| 722.9 | Other and unspecified disc disorder | 14 | musculoskeletal | 32 | 32 | 32 |
| 723 | Other disorders of cervical region | 14 | musculoskeletal | 48 | 48 | 48 |
| 723.1 | Torticollis | 14 | musculoskeletal | 14 | 14 | 14 |
| 724 | Other and unspecified disorders of back | 14 | musculoskeletal | 133 | 133 | 133 |
| 724.1 | Disorders of sacrum | 14 | musculoskeletal | 25 | 25 | 25 |
| 724.2 | Disorders of coccyx | 14 | musculoskeletal | 8 | 8 | 8 |
| 724.8 | Other symptoms referable to back | 14 | musculoskeletal | 82 | 82 | 82 |
| 724.9 | Other unspecified back disorders | 14 | musculoskeletal | 4 | 4 | 4 |
| 726 | Peripheral enthesopathies and allied syndromes | 14 | musculoskeletal | 371 | 371 | 371 |
| 726.1 | Enthesopathy | 14 | musculoskeletal | 172 | 172 | 172 |
| 726.2 | Synoviopathy | 14 | musculoskeletal | 22 | 22 | 22 |
| 726.3 | Bursitis | 14 | musculoskeletal | 39 | 39 | 39 |
| 726.4 | Calcaneal spur; Exostosis NOS | 14 | musculoskeletal | 7 | 7 | 7 |
| 727 | Other disorders of synovium, tendon, and bursa | 14 | musculoskeletal | 218 | 218 | 218 |
| 727.1 | Synovitis and tenosynovitis | 14 | musculoskeletal | 95 | 95 | 95 |
| 727.2 | Bursitis disorders | 14 | musculoskeletal | 4 | 4 | 4 |
| 727.4 | Ganglion and cyst of synovium, tendon, and bursa | 14 | musculoskeletal | 23 | 23 | 23 |
| 727.5 | Rupture of synovium | 14 | musculoskeletal | 17 | 17 | 17 |
| 727.6 | Rupture of tendon, nontraumatic | 14 | musculoskeletal | 26 | 26 | 26 |
| 727.7 | Contracture of tendon (sheath) | 14 | musculoskeletal | 1 | 1 | 1 |
| 727.8 | Plica syndrome | 14 | musculoskeletal | 1 | 1 | 1 |
| 728 | Disorders of muscle, ligament, and fascia | 14 | musculoskeletal | 3 | 3 | 3 |
| 728.1 | Muscular calcification and ossification | 14 | musculoskeletal | 0 | 0 | 0 |
| 728.2 | Laxity of ligament or hypermobility syndrome | 14 | musculoskeletal | 2 | 2 | 2 |
| 728.7 | Fasciitis | 14 | musculoskeletal | 79 | 79 | 79 |
| 728.71 | Contracture of palmar fascia [Dupuytren's disease] | 14 | musculoskeletal | 7 | 7 | 7 |
| 729 | Other disorders of soft tissues | 14 | musculoskeletal | 13 | 13 | 13 |
| 729.1 | Rheumatism, unspecified and fibrositis | 14 | musculoskeletal | 0 | 0 | 0 |
| 729.3 | Panniculitis | 14 | musculoskeletal | 2 | 2 | 2 |
| 729.7 | Nontraumatic compartment syndrome | 14 | musculoskeletal | 0 | 0 | 0 |
| 731 | Osteitis deformans and osteopathies associated with other disorders classified elsewhere | 14 | musculoskeletal | 0 | 0 | 0 |
| 731.1 | Osteitis deformans [Paget's disease of bone] | 14 | musculoskeletal | 0 | 0 | 0 |
| 732 | Osteochondropathies | 14 | musculoskeletal | 5 | 5 | 5 |
| 732.1 | Juvenile osteochondrosis | 14 | musculoskeletal | 1 | 1 | 1 |
| 732.7 | Osteochondritis dissecans | 14 | musculoskeletal | 1 | 1 | 1 |
| 733 | Other disorders of bone and cartilage | 14 | musculoskeletal | 383 | 383 | 383 |
| 733.2 | Cyst of bone | 14 | musculoskeletal | 6 | 6 | 6 |
| 733.4 | Aseptic necrosis of bone | 14 | musculoskeletal | 4 | 4 | 4 |
| 733.6 | Costochondritis | 14 | musculoskeletal | 27 | 27 | 27 |
| 733.8 | Malunion and nonunion of fracture | 14 | musculoskeletal | 6 | 6 | 6 |
| 733.9 | Chondromalacia | 14 | musculoskeletal | 4 | 4 | 4 |
| 735 | Acquired foot deformities | 14 | musculoskeletal | 95 | 95 | 95 |
| 735.1 | Flat foot | 14 | musculoskeletal | 8 | 8 | 8 |
| 735.2 | Acquired toe deformities | 14 | musculoskeletal | 26 | 26 | 26 |
| 735.21 | Hammer toe (acquired) | 14 | musculoskeletal | 15 | 15 | 15 |
| 735.22 | Claw toe (acquired) | 14 | musculoskeletal | 0 | 0 | 0 |
| 735.23 | Hallux rigidus | 14 | musculoskeletal | 5 | 5 | 5 |
| 735.3 | Hallux valgus (Bunion) | 14 | musculoskeletal | 32 | 32 | 32 |
| 736 | Other acquired deformities of limbs | 14 | musculoskeletal | 11 | 11 | 11 |
| 736.1 | Acquired deformities of forearm | 14 | musculoskeletal | 0 | 0 | 0 |
| 736.2 | Acquired deformities of finger | 14 | musculoskeletal | 3 | 3 | 3 |
| 736.3 | Acquired deformities of hip | 14 | musculoskeletal | 0 | 0 | 0 |

| | | | | | | |
|---|---|---|---|---|---|---|
| 736.4 | Genu valgum or varum (acquired) | 14 | musculoskeletal | 3 | 3 | 3 |
| 736.5 | Acquired deformities of knee | 14 | musculoskeletal | 0 | 0 | 0 |
| 736.6 | Unequal leg length (acquired) | 14 | musculoskeletal | 3 | 3 | 3 |
| 737 | Curvature of spine | 14 | musculoskeletal | 77 | 77 | 77 |
| 737.1 | Kyphosis (acquired) | 14 | musculoskeletal | 9 | 9 | 9 |
| 737.2 | Lordosis (acquired) | 14 | musculoskeletal | 0 | 0 | 0 |
| 737.3 | Kyphoscoliosis and scoliosis | 14 | musculoskeletal | 60 | 60 | 60 |
| 738 | Other acquired musculoskeletal deformity | 14 | musculoskeletal | 56 | 56 | 56 |
| 738.4 | Acquired spondylolisthesis | 14 | musculoskeletal | 49 | 49 | 49 |
| 739 | Contracture of joint | 14 | musculoskeletal | 6 | 6 | 6 |
| 740 | Osteoarthrosis | 14 | musculoskeletal | 516 | 516 | 516 |
| 740.1 | Osteoarthritis; localized | 14 | musculoskeletal | 318 | 318 | 318 |
| 740.11 | Osteoarthrosis, localized, primary | 14 | musculoskeletal | 270 | 270 | 270 |
| 740.12 | Osteoarthrosis, localized, secondary | 14 | musculoskeletal | 8 | 8 | 8 |
| 740.2 | Osteoarthrosis, generalized | 14 | musculoskeletal | 72 | 72 | 72 |
| 740.3 | Osteoarthrosis involving more than one site, but not specified as generalized | 14 | musculoskeletal | 10 | 10 | 10 |
| 740.9 | Osteoarthrosis NOS | 14 | musculoskeletal | 196 | 196 | 196 |
| 741 | Symptoms and disorders of the joints | 14 | musculoskeletal | 134 | 134 | 134 |
| 741.1 | Ankylosis of joint | 14 | musculoskeletal | 1 | 1 | 1 |
| 741.2 | Stiffness of joint | 14 | musculoskeletal | 9 | 9 | 9 |
| 741.3 | Difficulty in walking | 14 | musculoskeletal | 4 | 4 | 4 |
| 741.4 | Joint effusions | 14 | musculoskeletal | 70 | 70 | 70 |
| 741.5 | Hemarthrosis | 14 | musculoskeletal | 2 | 2 | 2 |
| 741.6 | Villonodular synovitis | 14 | musculoskeletal | 0 | 0 | 0 |
| 742 | Derangement of joint, non-traumatic | 14 | musculoskeletal | 31 | 31 | 31 |
| 742.1 | Loose body in joint | 14 | musculoskeletal | 0 | 0 | 0 |
| 742.2 | Pathological, developmental or recurrent dislocation | 14 | musculoskeletal | 3 | 3 | 3 |
| 742.8 | Articular cartilage disorder | 14 | musculoskeletal | 8 | 8 | 8 |
| 742.9 | Other derangement of joint | 14 | musculoskeletal | 17 | 17 | 17 |
| 743 | Osteoporosis, osteopenia and pathological fracture | 14 | musculoskeletal | 172 | 172 | 172 |
| 743.1 | Osteoporosis | 14 | musculoskeletal | 34 | 34 | 34 |
| 743.11 | Osteoporosis NOS | 14 | musculoskeletal | 1 | 1 | 1 |
| 743.12 | Senile osteoporosis | 14 | musculoskeletal | 21 | 21 | 21 |
| 743.13 | Other specified osteoporosis | 14 | musculoskeletal | 8 | 8 | 8 |
| 743.2 | Pathologic fracture | 14 | musculoskeletal | 13 | 13 | 13 |
| 743.21 | Pathologic fracture of vertebrae | 14 | musculoskeletal | 8 | 8 | 8 |
| 743.22 | Pathologic fracture of femur | 14 | musculoskeletal | 0 | 0 | 0 |
| 743.4 | Stress fracture | 14 | musculoskeletal | 5 | 5 | 5 |
| 743.9 | Osteopenia or other disorder of bone and cartilage | 14 | musculoskeletal | 16 | 16 | 16 |
| 745 | Pain in joint | 14 | musculoskeletal | 1400 | 1400 | 1400 |
| 747 | Cardiac and circulatory congenital anomalies | 15 | congenital anomalies | 36 | 36 | 36 |
| 747.1 | Cardiac congenital anomalies | 15 | congenital anomalies | 28 | 28 | 28 |
| 747.11 | Cardiac shunt/ heart septal defect | 15 | congenital anomalies | 14 | 14 | 14 |
| 747.12 | Valvular heart disease/ heart chambers | 15 | congenital anomalies | 11 | 11 | 11 |
| 747.13 | Congenital anomalies of great vessels | 15 | congenital anomalies | 3 | 3 | 3 |
| 747.2 | Congenital anomalies of peripheral vascular system | 15 | congenital anomalies | 6 | 6 | 6 |
| 748 | Anomalies of respiratory system, congenital | 15 | congenital anomalies | 0 | 0 | 0 |
| 749 | Congenital anomalies of face and neck | 15 | congenital anomalies | 5 | 5 | 5 |
| 749.1 | Cleft palate | 15 | congenital anomalies | 1 | 1 | 1 |
| 749.2 | Congenital anomalies of skull and face bones | 15 | congenital anomalies | 0 | 0 | 0 |
| 750 | Digestive congenital anomalies | 15 | congenital anomalies | 13 | 13 | 13 |
| 750.1 | Upper gastrointestinal congenital anomalies | 15 | congenital anomalies | 3 | 3 | 3 |
| 750.11 | Esophageal atresia/tracheoesophageal fistula | 15 | congenital anomalies | 3 | 3 | 3 |
| 750.13 | Congenital anomalies of mouth/tongue | 15 | congenital anomalies | 0 | 0 | 0 |
| 750.14 | Congenital anomalies of esophagus | 15 | congenital anomalies | 0 | 0 | 0 |
| 750.15 | Congenital anomalies of stomach | 15 | congenital anomalies | 0 | 0 | 0 |
| 750.2 | Lower gastrointestinal congenital anomalies | 15 | congenital anomalies | 9 | 9 | 9 |
| 750.21 | Congenital anomalies of intestine | 15 | congenital anomalies | 1 | 1 | 1 |
| 750.22 | Congenital anomaly of gallbladder, bile ducts, liver, pancreas | 15 | congenital anomalies | 8 | 8 | 8 |
| 750.5 | Congenital hypertrophic pyloric stenosis | 15 | congenital anomalies | 0 | 0 | 0 |
| 751 | Genitourinary congenital anomalies | 15 | congenital anomalies | 46 | 46 | 46 |
| 751.1 | Congenital anomalies of genital organs | 15 | congenital anomalies | 3 | 3 | 3 |
| 751.11 | Congenital anomalies of female genital organs | 15 | congenital anomalies | 1 | 1 | 1 |
| 751.12 | Congenital anomalies of male genital organs | 15 | congenital anomalies | 1 | 1 | 1 |
| 751.2 | Congenital anomalies of urinary system | 15 | congenital anomalies | 35 | 35 | 35 |
| 751.21 | Cystic kidney disease | 15 | congenital anomalies | 26 | 26 | 26 |
| 751.22 | Other specified congenital anomalies of kidney | 15 | congenital anomalies | 3 | 3 | 3 |

| | | | | | | |
|---|---|---|---|---|---|---|
| 751.3 | Obstructive genitourinary defect | 15 | congenital anomalies | 2 | 2 | 2 |
| 752 | Nervous system congenital anomalies | 15 | congenital anomalies | 1 | 1 | 1 |
| 752.1 | Neural tube defects | 15 | congenital anomalies | 0 | 0 | 0 |
| 752.11 | Spina bifida | 15 | congenital anomalies | 0 | 0 | 0 |
| 752.2 | Other specified congenital anomalies of nervous system | 15 | congenital anomalies | 1 | 1 | 1 |
| 753 | Congenital anomalies of the eye | 15 | congenital anomalies | 1 | 1 | 1 |
| 753.1 | Congenital cataract and lens anomalies | 15 | congenital anomalies | 0 | 0 | 0 |
| 753.2 | Congenital anomalies of posterior segment of eye | 15 | congenital anomalies | 0 | 0 | 0 |
| 754 | Congenital musculoskeletal deformities of spine | 15 | congenital anomalies | 6 | 6 | 6 |
| 754.1 | Lumbosacral spondylolysis, congenital | 15 | congenital anomalies | 1 | 1 | 1 |
| 754.2 | Spondylolisthesis, congenital | 15 | congenital anomalies | 3 | 3 | 3 |
| 755 | Congenital anomalies of limbs | 15 | congenital anomalies | 10 | 10 | 10 |
| 755.1 | Congenital deformities of feet | 15 | congenital anomalies | 3 | 3 | 3 |
| 755.3 | Congenital anomaly of fingers/toes | 15 | congenital anomalies | 0 | 0 | 0 |
| 755.4 | Congenital anomalies of upper limb, including shoulder girdle | 15 | congenital anomalies | 1 | 1 | 1 |
| 755.6 | Other congenital anomalies of lower limb, including pelvic girdle | 15 | congenital anomalies | 5 | 5 | 5 |
| 755.61 | Congenital hip dysplasia and deformity | 15 | congenital anomalies | 5 | 5 | 5 |
| 756 | Other congenital musculoskeletal anomalies | 15 | congenital anomalies | 10 | 10 | 10 |
| 756.1 | Congenital anomalies of abdominal wall; diaphram | 15 | congenital anomalies | 0 | 0 | 0 |
| 756.2 | Pectus and other congenital anomalies of ribs/sternum | 15 | congenital anomalies | 4 | 4 | 4 |
| 756.21 | Pectus excavatum | 15 | congenital anomalies | 2 | 2 | 2 |
| 756.22 | Pectus carinatum | 15 | congenital anomalies | 0 | 0 | 0 |
| 756.3 | Congenital anomalies of muscle, tendon, fascia, and connective tissue | 15 | congenital anomalies | 5 | 5 | 5 |
| 756.5 | Congenital osteodystrophies | 15 | congenital anomalies | 0 | 0 | 0 |
| 757 | Congenital anomalies of the integument | 15 | congenital anomalies | 1 | 1 | 1 |
| 758 | Chromosomal anomalies and genetic disorders | 15 | congenital anomalies | 2 | 2 | 2 |
| 758.1 | Chromosomal anomalies | 15 | congenital anomalies | 1 | 1 | 1 |
| 759 | Other and unspecified congenital anomalies | 15 | congenital anomalies | 8 | 8 | 8 |
| 759.1 | Anomalies of endocrine glands, congenital | 15 | congenital anomalies | 2 | 2 | 2 |
| 760 | Back pain | 17 | symptoms | 1036 | 1036 | 1036 |
| 761 | Cervicalgia | 17 | symptoms | 439 | 439 | 439 |
| 763 | Thoracic or lumbosacral neuritis or radiculitis, unspecified | 17 | symptoms | 221 | 221 | 221 |
| 764 | Sciatica | 17 | symptoms | 293 | 293 | 293 |
| 765 | Cervical radiculitis | 17 | symptoms | 98 | 98 | 98 |
| 766 | Neuralgia, neuritis, and radiculitis NOS | 17 | symptoms | 53 | 53 | 53 |
| 767 | Cervicocranial/Cervicobrachial syndrome | 17 | symptoms | 0 | 0 | 0 |
| 769 | Nonallopathic lesions NEC | 17 | symptoms | 28 | 28 | 28 |
| 770 | Myalgia and myositis unspecified | 17 | symptoms | 245 | 245 | 245 |
| 771 | Musculoskeletal symptoms referable to limbs | 17 | symptoms | 25 | 25 | 25 |
| 771.1 | Swelling of limb | 17 | symptoms | 50 | 50 | 50 |
| 771.2 | Cramp of limb | 17 | symptoms | 31 | 31 | 31 |
| 772 | Symptoms of the muscles | 17 | symptoms | 9 | 9 | 9 |
| 772.1 | Muscular wasting and disuse atrophy | 17 | symptoms | 2 | 2 | 2 |
| 772.2 | Spasm of muscle | 17 | symptoms | 132 | 132 | 132 |
| 772.3 | Muscle weakness | 17 | symptoms | 27 | 27 | 27 |
| 772.4 | Rhabdomyolysis | 17 | symptoms | 2 | 2 | 2 |
| 772.6 | Facial weakness | 17 | symptoms | 2 | 2 | 2 |
| 780 | Hypothermia/Chills | 17 | symptoms | 15 | 15 | 15 |
| 781 | Symptoms involving nervous and musculoskeletal systems | 17 | symptoms | 113 | 113 | 113 |
| 781.1 | Loss of height | 17 | symptoms | 2 | 2 | 2 |
| 781.2 | Abnormal posture | 17 | symptoms | 0 | 0 | 0 |
| 782 | Symptoms involving skin and other integumentary tissue | 17 | symptoms | 236 | 236 | 236 |
| 782.3 | Edema | 17 | symptoms | 191 | 191 | 191 |
| 782.6 | Pallor and flushing | 17 | symptoms | 27 | 27 | 27 |
| 783 | Fever of unknown origin | 17 | symptoms | 234 | 234 | 234 |
| 783.1 | Postprocedural fever | 17 | symptoms | 2 | 2 | 2 |
| 785 | Abdominal pain | 17 | symptoms | 929 | 929 | 929 |
| 788 | Syncope and collapse | 17 | symptoms | 209 | 209 | 209 |
| 789 | Nausea and vomiting | 17 | symptoms | 544 | 544 | 544 |
| 789.1 | Persistent vomiting | 17 | symptoms | 9 | 9 | 9 |
| 790 | Nonspecific findings on examination of blood | 17 | symptoms | 18 | 18 | 18 |
| 790.1 | Elevated sedimentation rate | 17 | symptoms | 10 | 10 | 10 |
| 790.6 | Other abnormal blood chemistry | 17 | symptoms | 125 | 125 | 125 |
| 790.8 | Elevated C-reactive protein (CRP) | 17 | symptoms | 16 | 16 | 16 |
| 790.9 | Abnormal arterial blood gases | 17 | symptoms | 2 | 2 | 2 |
| 792 | Abnormal Papanicolaou smear of cervix and cervical HPV | 17 | symptoms | 40 | 40 | 40 |
| 792.1 | Papanicolaou smear of cervix or vagina with atypical squamous cells | 17 | symptoms | 30 | 30 | 30 |

| | | | | | | |
|---|---|---|---|---|---|---|
| 794 | Abnormal results of other function studies (bladder, pancreas, placenta, spleen, etc) | 17 | symptoms | 0 | 0 | 0 |
| 795 | Other and nonspecific abnormal cytological, histological and immunological findings | 17 | symptoms | 8 | 8 | 8 |
| 795.8 | Abnormal tumor markers | 17 | symptoms | 2 | 2 | 2 |
| 795.81 | Elevated carcinoembryonic antigen [CEA] | 17 | symptoms | 1 | 1 | 1 |
| 795.82 | Elevated cancer antigen 125 [CA 125] | 17 | symptoms | 1 | 1 | 1 |
| 796 | Elevated prostate specific antigen [PSA] | 17 | symptoms | 64 | 64 | 64 |
| 797 | Shock | 17 | symptoms | 7 | 7 | 7 |
| 797.1 | Cardiogenic shock | 17 | symptoms | 0 | 0 | 0 |
| 798 | Malaise and fatigue | 17 | symptoms | 1164 | 1164 | 1164 |
| 798.1 | Chronic fatigue syndrome | 17 | symptoms | 19 | 19 | 19 |
| 800 | Fracture of lower limb | 18 | injuries & poisonings | 56 | 56 | 56 |
| 800.1 | Fracture of neck of femur | 18 | injuries & poisonings | 18 | 18 | 18 |
| 800.2 | Fracture of unspecified part of femur | 18 | injuries & poisonings | 2 | 2 | 2 |
| 800.3 | Fracture of tibia and fibula | 18 | injuries & poisonings | 21 | 21 | 21 |
| 800.4 | Fracture of patella | 18 | injuries & poisonings | 9 | 9 | 9 |
| 801 | Fracture of ankle and foot | 18 | injuries & poisonings | 95 | 95 | 95 |
| 801.1 | Fracture of foot | 18 | injuries & poisonings | 32 | 32 | 32 |
| 802 | Fracture of pelvis | 18 | injuries & poisonings | 6 | 6 | 6 |
| 803 | Fracture of upper limb | 18 | injuries & poisonings | 91 | 91 | 91 |
| 803.1 | Fracture of humerus | 18 | injuries & poisonings | 26 | 26 | 26 |
| 803.2 | Fracture of radius and ulna | 18 | injuries & poisonings | 45 | 45 | 45 |
| 803.21 | Colles' fracture | 18 | injuries & poisonings | 3 | 3 | 3 |
| 803.3 | Fracture of clavicle or scapula | 18 | injuries & poisonings | 14 | 14 | 14 |
| 804 | Fracture of hand or wrist | 18 | injuries & poisonings | 38 | 38 | 38 |
| 805 | Fracture of vertebral column without mention of spinal cord injury | 18 | injuries & poisonings | 49 | 49 | 49 |
| 807 | Fracture of ribs | 18 | injuries & poisonings | 36 | 36 | 36 |
| 809 | Fracture of unspecified bones | 18 | injuries & poisonings | 21 | 21 | 21 |
| 816 | Cerebral laceration and contusion | 18 | injuries & poisonings | 0 | 0 | 0 |
| 817 | Concussion | 18 | injuries & poisonings | 26 | 26 | 26 |
| 818 | Intracranial hemorrhage (injury) | 18 | injuries & poisonings | 8 | 8 | 8 |
| 818.1 | Subdural hemorrhage (injury) | 18 | injuries & poisonings | 5 | 5 | 5 |
| 818.2 | Subarachnoid hemorrhage (injury) | 18 | injuries & poisonings | 0 | 0 | 0 |
| 819 | Skull and face fracture and other intercranial injury | 18 | injuries & poisonings | 41 | 41 | 41 |
| 823 | Fracture of tibia and fibula | 18 | injuries & poisonings | 0 | 0 | 0 |
| 830 | Dislocation | 18 | injuries & poisonings | 120 | 120 | 120 |
| 835 | Internal derangement of knee | 18 | injuries & poisonings | 93 | 93 | 93 |
| 836 | Traumatic arthropathy | 18 | injuries & poisonings | 2 | 2 | 2 |
| 840 | Sprains and strains | 18 | injuries & poisonings | 445 | 445 | 445 |
| 840.1 | Muscle/tendon sprain | 18 | injuries & poisonings | 10 | 10 | 10 |
| 840.2 | Rotator cuff (capsule) sprain | 18 | injuries & poisonings | 15 | 15 | 15 |
| 840.3 | Joint/ligament sprain | 18 | injuries & poisonings | 34 | 34 | 34 |
| 841 | Sprains and strains of back and neck | 18 | injuries & poisonings | 257 | 257 | 257 |
| 842 | Other sprains and strains | 18 | injuries & poisonings | 51 | 51 | 51 |
| 850 | Hemorrhage or hematoma complicating a procedure | 18 | injuries & poisonings | 9 | 9 | 9 |
| 851 | Complications of transplants and reattached limbs | 18 | injuries & poisonings | 0 | 0 | 0 |
| 853 | Complication of colostomy or enterostomy | 18 | injuries & poisonings | 4 | 4 | 4 |
| 854 | Complications of cardiac/vascular device, implant, and graft | 18 | injuries & poisonings | 2 | 2 | 2 |
| 855 | Complication of nervous system device, implant, and graft | 18 | injuries & poisonings | 0 | 0 | 0 |
| 856 | Vascular complications of surgery and medical procedures | 18 | injuries & poisonings | 0 | 0 | 0 |
| 857 | Mechanical complication of unspecified genitourinary device, implant, and graft | 18 | injuries & poisonings | 6 | 6 | 6 |
| 858 | Complication of internal orthopedic device | 18 | injuries & poisonings | 16 | 16 | 16 |
| 859 | Complication due to other implant and internal device | 18 | injuries & poisonings | 11 | 11 | 11 |
| 860 | Bone marrow or stem cell transplant | 2 | neoplasms | 2 | 2 | 2 |
| 870 | Open wounds of head; neck; and trunk | 18 | injuries & poisonings | 121 | 121 | 121 |
| 870.1 | Open wound or laceration of eye or eyelid | 18 | injuries & poisonings | 2 | 2 | 2 |
| 870.2 | Open wound of ear | 18 | injuries & poisonings | 0 | 0 | 0 |
| 870.3 | Other open wound of head and face | 18 | injuries & poisonings | 32 | 32 | 32 |
| 870.4 | Open wound of nose and sinus | 18 | injuries & poisonings | 3 | 3 | 3 |
| 870.5 | Open wound of lip and mouth | 18 | injuries & poisonings | 9 | 9 | 9 |
| 870.6 | Open wound of neck | 18 | injuries & poisonings | 1 | 1 | 1 |
| 870.8 | Open wound of genital organs | 18 | injuries & poisonings | 2 | 2 | 2 |
| 871 | Open wounds of extremities | 18 | injuries & poisonings | 243 | 243 | 243 |
| 871.1 | Open wound of hand except finger(s) | 18 | injuries & poisonings | 50 | 50 | 50 |
| 871.2 | Open wound of finger(s) | 18 | injuries & poisonings | 112 | 112 | 112 |
| 871.3 | Open wound of foot except toe(s) alone | 18 | injuries & poisonings | 14 | 14 | 14 |
| 871.4 | Open wound of toe(s) | 18 | injuries & poisonings | 16 | 16 | 16 |
| 872 | Traumatic amputation | 18 | injuries & poisonings | 4 | 4 | 4 |

| | | | | | | |
|---|---|---|---|---|---|---|
| 874 | Complication of amputation stump | 18 | injuries & poisonings | 0 | 0 | 0 |
| 875 | Non-healing surgical wound | 18 | injuries & poisonings | 2 | 2 | 2 |
| 876 | Posttraumatic wound infection not elsewhere classified | 18 | injuries & poisonings | 43 | 43 | 43 |
| 907 | Injuries to the nervous system | 18 | injuries & poisonings | 8 | 8 | 8 |
| 910 | Superficial injury, infected | 18 | injuries & poisonings | 17 | 17 | 17 |
| 911 | Blister | 18 | injuries & poisonings | 8 | 8 | 8 |
| 912 | Insect bite | 18 | injuries & poisonings | 38 | 38 | 38 |
| 913 | Toxic effect of venom | 18 | injuries & poisonings | 16 | 16 | 16 |
| 916 | Contusion | 18 | injuries & poisonings | 295 | 295 | 295 |
| 930 | Allergic reaction to food | 18 | injuries & poisonings | 38 | 38 | 38 |
| 931 | Contact dermatitis and other eczema due to plants [except food] | 18 | injuries & poisonings | 1 | 1 | 1 |
| 938 | Dermatitis due to solar radiation | 18 | injuries & poisonings | 16 | 16 | 16 |
| 938.1 | Acute dermatitis due to solar radiation | 18 | injuries & poisonings | 8 | 8 | 8 |
| 938.2 | Chronic dermatitis due to solar radiation | 18 | injuries & poisonings | 1 | 1 | 1 |
| 939 | Atopic/contact dermatitis due to other or unspecified | 13 | dermatologic | 246 | 246 | 246 |
| 939.1 | Contact and allergic dermatitis of eyelid | 13 | dermatologic | 3 | 3 | 3 |
| 941 | Adverse reaction to serum or vaccine | 18 | injuries & poisonings | 1 | 1 | 1 |
| 942 | Infusion and transfusion reaction | 18 | injuries & poisonings | 1 | 1 | 1 |
| 946 | Anaphylactic shock NOS | 18 | injuries & poisonings | 10 | 10 | 10 |
| 947 | Urticaria | 18 | injuries & poisonings | 75 | 75 | 75 |
| 949 | Allergies, other | 18 | injuries & poisonings | 216 | 216 | 216 |
| 949.1 | Diaper or napkin rash | 18 | injuries & poisonings | 1 | 1 | 1 |
| 952 | Spinal cord injury without evidence of spinal bone injury | 18 | injuries & poisonings | 2 | 2 | 2 |
| 957 | Injury to other and unspecified nerves | 18 | injuries & poisonings | 0 | 0 | 0 |
| 958 | Certain early complications of trauma or procedure | 18 | injuries & poisonings | 4 | 4 | 4 |
| 958.1 | Postoperative shock | 18 | injuries & poisonings | 1 | 1 | 1 |
| 958.2 | Traumatic and surgical subcutaneous emphysema | 18 | injuries & poisonings | 0 | 0 | 0 |
| 960 | Poisoning by antibiotics | 18 | injuries & poisonings | 130 | 130 | 130 |
| 960.1 | Adverse effects of antibacterials (not penicillins) | 18 | injuries & poisonings | 0 | 0 | 0 |
| 960.2 | Allergy/adverse effect of penicillin | 18 | injuries & poisonings | 55 | 55 | 55 |
| 960.3 | Poisoning by antifungal antibiotics | 18 | injuries & poisonings | 0 | 0 | 0 |
| 961 | Poisoning by other anti-infectives | 18 | injuries & poisonings | 1 | 1 | 1 |
| 961.1 | Poisoning/allergy of sulfonamides | 18 | injuries & poisonings | 36 | 36 | 36 |
| 962 | Poisoning by hormones and synthetic substitutes | 18 | injuries & poisonings | 12 | 12 | 12 |
| 962.1 | Adrenal cortical steroids causing adverse effects in therapeutic use | 18 | injuries & poisonings | 9 | 9 | 9 |
| 962.2 | Insulins and antidiabetic agents causing adverse effects in therapeutic use | 18 | injuries & poisonings | 2 | 2 | 2 |
| 962.3 | Hormones and synthetic substitutes causing adverse effects in therapeutic use | 18 | injuries & poisonings | 1 | 1 | 1 |
| 963 | Poisoning by primarily systemic agents | 18 | injuries & poisonings | 17 | 17 | 17 |
| 963.1 | Antineoplastic and immunosuppressive drugs causing adverse effects | 18 | injuries & poisonings | 15 | 15 | 15 |
| 964 | Poisoning by agents primarily affecting blood constituents | 18 | injuries & poisonings | 1 | 1 | 1 |
| 964.1 | Anticoagulants causing adverse effects | 18 | injuries & poisonings | 1 | 1 | 1 |
| 965 | Poisoning by analgesics, antipyretics, and antirheumatics | 18 | injuries & poisonings | 13 | 13 | 13 |
| 965.1 | Opiates and related narcotics causing adverse effects in therapeutic use | 18 | injuries & poisonings | 41 | 41 | 41 |
| 965.2 | Antirheumatics causing adverse effects in therapeutic use | 18 | injuries & poisonings | 0 | 0 | 0 |
| 965.3 | Salicylates causing adverse effects in therapeutic use | 18 | injuries & poisonings | 0 | 0 | 0 |
| 966 | Poisoning by anticonvulsants and anti-Parkinsonism drugs | 18 | injuries & poisonings | 0 | 0 | 0 |
| 967 | Adverse effects of sedatives or other central nervous system depressants and anesthetics | 18 | injuries & poisonings | 4 | 4 | 4 |
| 969 | Poisoning by psychotropic agents | 18 | injuries & poisonings | 4 | 4 | 4 |
| 971 | Poisoning by drugs primarily affecting the autonomic nervous system | 18 | injuries & poisonings | 3 | 3 | 3 |
| 972 | Poisoning by agents primarily affecting the cardiovascular system | 18 | injuries & poisonings | 19 | 19 | 19 |
| 972.1 | Cardiac rhythm regulators causing adverse effects in therapeutic use | 18 | injuries & poisonings | 1 | 1 | 1 |
| 972.2 | Antilipemic and antiarteriosclerotic drugs causing adverse effects in therapeutic use | 18 | injuries & poisonings | 3 | 3 | 3 |
| 972.6 | Antihypertensive agents causing adverse effects | 18 | injuries & poisonings | 15 | 15 | 15 |
| 973 | Poisoning by agents primarily affecting the gastrointestinal system | 18 | injuries & poisonings | 0 | 0 | 0 |
| 974 | Poisoning by water, mineral, and uric acid metabolism drugs | 18 | injuries & poisonings | 4 | 4 | 4 |
| 975 | Poisoning by agents primarily acting on the smooth and skeletal muscles and respiratory system | 18 | injuries & poisonings | 2 | 2 | 2 |
| 976 | Poisoning by agents primarily affecting skin & mucous membrane, ophthalmological, otorhinolaryngological, & dental drugs | 18 | injuries & poisonings | 1 | 1 | 1 |
| 977 | Personal history of allergy to medicinal agents | 18 | injuries & poisonings | 18 | 18 | 18 |
| 979 | Adverse drug events and drug allergies | 18 | injuries & poisonings | 49 | 49 | 49 |
| 980 | Encounter for long-term (current) use of antibiotics | 1 | infectious diseases | 2 | 2 | 2 |
| 981 | Toxic effect of (non-ethyl) alcohol and petroleum and other solvents | 18 | injuries & poisonings | 0 | 0 | 0 |
| 983 | Toxic effect of corrosive aromatics, acids, and caustic alkalis | 18 | injuries & poisonings | 2 | 2 | 2 |
| 984 | Toxic effect of lead and its compounds (including fumes) | 18 | injuries & poisonings | 0 | 0 | 0 |
| 985 | Toxic effect of other metals | 18 | injuries & poisonings | 1 | 1 | 1 |
| 986 | Toxic effect of carbon monoxide | 18 | injuries & poisonings | 0 | 0 | 0 |
| 987 | Toxic effect of other gases, fumes, or vapors | 18 | injuries & poisonings | 1 | 1 | 1 |
| 988 | Toxic effect of noxious substances eaten as food | 18 | injuries & poisonings | 1 | 1 | 1 |

| | | | | | | |
|---|---|---|---|---|---|---|
| 989 | Toxic effect of other substances, chiefly nonmedicinal as to source | 18 | injuries & poisonings | 1 | 1 | 1 |
| 990 | Effects radiation NOS | 18 | injuries & poisonings | 6 | 6 | 6 |
| 994 | Sepsis and SIRS | 18 | injuries & poisonings | 65 | 65 | 65 |
| 994.1 | Systemic inflammatory response syndrome (SIRS) | 18 | injuries & poisonings | 1 | 1 | 1 |
| 994.2 | Sepsis | 18 | injuries & poisonings | 55 | 55 | 55 |
| 994.21 | Septic shock | 18 | injuries & poisonings | 16 | 16 | 16 |
